# Generation and application of novel hES cell reporter lines for the differentiation and maturation of hPS cell-derived islet-like clusters

**DOI:** 10.1101/2024.07.15.603551

**Authors:** Elisa Zanfrini, Manuj Bandral, Luka Jarc, Maria Alejandra Ramirez-Torres, Daniela Pezzolla, Vida Kufrin, Eva Rodriguez-Aznar, Ana Karen Mojica Avila, Christian Cohrs, Stephan Speier, Katrin Neumann, Anthony Gavalas

## Abstract

The significant advances in the differentiation of human pluripotent stem (hPS) cells into pancreatic endocrine cells, including functional β-cells, have been based on a detailed understanding of the underlying developmental mechanisms. However, the final differentiation steps, leading from endocrine progenitors to mono-hormonal and mature pancreatic endocrine cells, remain to be fully understood and this is reflected in the remaining shortcomings of the hPS cell-derived islet cells (SC-islet cells), which include a lack of β-cell maturation and variability among different cell lines. Additional signals and modifications of the final differentiation steps will have to be assessed in a combinatorial manner to address the remaining issues and appropriate reporter lines would be useful in this undertaking.

Here we report the generation and functional validation of human pluripotent stem cell reporter lines that can monitor the generation of INS^+^ and GCG^+^ cells and their resolution into mono-hormonal cells (*INS ^eGFP^, INS ^eGFP^/ GCG ^mCHERRY^*) as well as β-cell maturation (*INS ^eGFP^/ MAFA ^mCHERRY^*) and function (*INS ^GCaMP^*^6^). The reporter hPS cell lines maintained strong and widespread expression of pluripotency markers and differentiated efficiently into definitive endoderm and pancreatic progenitor (PP) cells. PP cells from all lines differentiated efficiently into islet cell clusters that robustly expressed the corresponding reporters and contained glucose-responsive, insulin-producing cells.

To demonstrate the applicability of these hES cell reporter lines in a high-content live imaging approach for the identification of optimal differentiation conditions, we adapted our differentiation procedure to generate SC-islet clusters in microwells. This allowed the live confocal imaging of multiple SC-islets for a single condition and, using this approach, we found that the use of the N21 supplement in the last stage of the differentiation increased the number of monohormonal β-cells without affecting the number of α-cells in the SC-islets.

The hPS cell reporter lines and the high-content live imaging approach described here will enable the efficient assessment of multiple conditions for the optimal differentiation and maturation of SC-islets.

## INTRODUCTION

Diabetes is a global epidemic that affects > 9% of the global population ^1^. Restoration of insulin independence with the maintenance of normoglycemia and prevention of diabetic complications are the ultimate goals of diabetes therapy. Improvements in the synthesis and delivery of recombinant insulin as well as the development of several other drugs have substantially decreased the morbidity and mortality associated with the disease. However, these life-long regimens do not result in a definitive cure and > 400 million people worldwide suffer from devastating secondary diabetes complications. In cases of severe diabetes, particularly Type 1, whole pancreas transplantation or pancreatic islet transplantation are recommended as they can restore good metabolic control and significantly reduce or eliminate insulin dependence. However, both approaches are critically limited by the scarcity of donor tissue and the requirement for immunosuppression which adversely affects the already limited β-cell self-renewal and has severe side effects ^2^. The genetic origins of β-cell contribution to both T1D and T2D are diverse and only partially understood ^3–5^. The availability of β-cells genetically identical to patients would provide an additional tool to better understand the molecular mechanisms of the disease and develop drugs for personalized treatments.

Advances in the generation of human pluripotent stem cell-derived pancreatic islet-like (SC-islet) cells suggested that this approach could provide an alternative to donor tissue transplantations. The differentiation is based on the application of developmental mechanisms whereby hPS cells are first differentiated into definitive endoderm (DE) and then, through additional intermediates, into pancreas progenitors (PP), pancreatic endocrine progenitors (PEP) and, finally, stem cell-derived islet (SC-islet) cells, including functional β-cells ^6–9^. However, these β-cells do not progress to full maturity, as indicated by their lower insulin production and secretion in response to physiological stimuli such as glucose or incretins, and the conversion efficiency of hPS cells into SC-islet cells remains relatively low. The final cultures retain considerable numbers of duct-like and enteroendocrine cells which inhibit functional maturation ^7,8,10–13^.

A recent study employed the combined use of the ISX9 NeuroD1 inducer and the general Wnt inhibitor Wnt-C59 in the generation of PEP cells to increase their conversion into SC-islet cells ^14^. In another recent study, differentiation efficiency and maturation were dramatically improved by modifying the final differentiation stage, accelerating mitotic exit with an antiproliferative aurora kinase inhibitor and extending the final stage of the differentiation for several weeks. The derived SC-islets showed robust glucose-stimulated insulin secretion (GSIS), including second-phase secretion and incretin response as well as cytoplasmic Ca^2+^ currents, a balanced proportion of α- and β- cells and higher, but still divergent, transcriptome similarity to human islets. Additionally, these SC-islets maintained features of immature cells with respect to glycolysis and mitochondrial metabolism in comparison to adult human islets ^15^. Maturation of β-cells is a gradual process occurring postnatally and is accompanied by enhanced expression and induction of essential transcription factors. In humans, postnatal maturation extends until the end of puberty ^16,17^ and it is, to a large extent, driven by the gradual upregulation of the transcription factor MAFA ^18^. In humans, reduced *MAFA* expression or missense MAFA mutations have been linked with poor GSIS and T2D ^19–21^. MAFA expression is particularly low in current hPS cell-derived β-cells ^11^ but very little is known about conditions that could enable *MAFA* upregulation. Finally, there is substantial variation in the differentiation efficiency of hPS cell lines, suggesting that differentiation procedures are still not sufficiently robust. Recent findings indicate that there are additional players, including signaling pathways, mechanical forces and metabolism, affecting endocrine lineage specification and endocrine cell maturation, that have not yet been widely leveraged in these procedures ^22–31^.

In summary, additional changes, particularly at the later stages, leading from PP cells to PEP cells and finally to SC-islets, are still necessary to establish a highly efficient differentiation procedure that will be reproducible across different hPS cell lines and will yield mature β-cells in the shortest time frame possible. The challenge arising is a large number of possible combinations when modifying different stages, with different compounds in variable concentrations and for application in different time windows. Fluorescent reporter lines used for high-content experiments would address this limitation. Reporter lines for the progenitor markers *PDX1*, *NKX6.1*, *SOX9* and *NEUROG3* have been generated and used to understand the generation and differentiation paths of pancreas progenitors in order to optimize the differentiation of hPS cells into SC-islet cells ^32–37^. However, the existing lines for the late differentiation stages are limited to single reporter lines for the *INS* ^34,38–40^, *PDX1* ^36^ and *ARX* loci ^41^ and they do not address the need to effectively assess differentiation efficiency, maturation and function. There is a report of an *INS ^eGFP^ / SOM ^RFP^* double reporter line that was generated using the Mel1 *INS ^GFP^* line ^39^ but the capacity of these cells to differentiate into functional β-cells was not reported ^42^.

Here we report the generation and functional assessment of heterozygous hPS cell reporter lines, including double reporter lines, suitable to monitor the generation of INS^+^ and GCG^+^ cells, their resolution into mono-hormonal cells (*INS ^eGFP^, INS ^eGFP^/ GCG ^mCHERRY^*) as well as β-cell maturation (*INS ^eGFP^/ MAFA ^mCHERRY^*) and function (*INS ^GCaMP^*^6^). GCaMP6 is an optimized fluorescent Ca^2+^ detecting protein^43^ and the *INS ^GCaMP^*^6^ line will allow bypassing the considerable drawbacks of dye Ca^2+^ indicators, such as staining of non-β-cells, photobleaching, leakage and uneven loading.

The lines were generated using CRISPR-Cas9 mediated recombination to target the loci in H1 hES cells. The reporter lines maintained strong and widespread expression of pluripotency markers and differentiated efficiently into DE and PP cells. PP cells from all lines differentiated into SC-islet clusters that robustly expressed the corresponding reporters and contained glucose-responsive, insulin-producing cells. We demonstrated that these hES cell lines, in combination with their differentiation into SC-islets in microwells, can be used in live imaging, high-content approaches, to identify better conditions for the differentiation and maturation of SC-islets. We found that the inclusion of the N21 supplement, in the last stage of differentiation, increased by more than 40% the number of monohormonal β-cells and their responsiveness.

The hPS cell reporter lines and the high-content live imaging approach described here will enable the efficient assessment of multiple conditions for the optimal differentiation and maturation of SC-islets while providing statistical power and minimizing the number of cells necessary to carry out these screens.

## MATERIALS AND METHODS

### Generation of the targeting vectors

All constructs were assembled using the Gibson assembly method of the linearized backbone vector and four PCR-generated fragments that corresponded to the 5’ homology arm, the T2A – reporter transgene, the selection cassette and the 3’ HA. Overlapping sequences and PCR primers were designed using the SnapGene software. The generation or origin of the plasmids is described in Table S1 ^43–45^ and for the assembly reactions we used the NEBuilder HiFi Assembly cloning kit (NEB E5520S). PCR reactions were carried out using the Q5 High-Fidelity DNA polymerase (NEB MO515) and plasmids were purified using the Qiagen EndoFree Plasmid Maxi Kit (Qiagen 12362).

### Maintenance and karyotyping of human pluripotent stem cells

The H1 hES cell line was purchased from WiCell (Wisconsin, USA) and was used to derive all reporter lines. Cells were maintained on cell culture dishes coated with hES cell qualified Corning Matrigel (BD Bioscience, 354277) diluted 1:50 with DMEM/F-12 (Gibco, 21331-020) and daily changes of mTeSR1 medium (STEMCELL Technologies, 85850) supplemented with 1x penicillin/streptomycin (Gibco, 15140-122). Cells were passaged at around 70% confluency, approximately every 4 days at a ratio of 1:6 to 1:9, as small aggregates using ReLeSR (STEMCELL Technologies, 05872). Cells from the parental cell line and the derived reporter lines used for the experiments here were analyzed for chromosomal abnormalities using standard G banding karyotyping. Cells were treated with 100 ng/ml KaryoMAX Colcemid solution (Thermo Fisher Scientific 15212012) for 4 h at 37°C, harvested and enlarged with 0.075 M KCl solution (Thermo Fisher Scientific 10575090) for 20 min at 37°C. After fixing with 3:1 methanol (VWR 20846.326) : glacial acetic acid (VWR 20102.292), cells were spread onto glass slides, stained with Giemsa and G-bandings of at least 20 metaphases were analyzed at the Institute of Human Genetics, Jena University, Germany. Cells were routinely tested for mycoplasma contamination by PCR as published previously ^46^.

### Gene editing of hES cells

Cells at passage 35-60 were harvested as a single cell suspension using TrypLE Express and 800K cells were nucleofected in 100 μl final volume containing either 2 μg of the Cas9 nickase expression vector ^47^ with 1 μg of each of the two sgRNAs and 10 μg of the targeting vector (for the *INS ^eGFP^* allele) or 6 μl Cas9-NLS protein (120 pmol, NEB, M0646M) with 2 μg of each of the sgRNAs (Table S2) and 10 μg linearized targeting vector (all other alleles). Nucleofections were done using Amaxa 4D and the H9 hES nucleofection program (CB150). Two sgRNAs were used to increase efficiency and were placed either closely on either side of the STOP codon of the target gene or immediately downstream or overlapping with it. The specificity of the sgRNAs was assessed using both the ChopChop (http://chopchop.cbu.uib.no/) ^48^ and CCTop (https://cctop.cos.uni-heidelberg.de/) ^49^ algorithms. For all sgRNAs used (Table S2), there were no likely off-target sites detected from either tool because alternative sites usually had a minimum of three mismatches. Nucleofected cells were equally seeded in three wells of a 6-well plate. Drug selection started three days after the nucleofection and lasted for six days. Geneticin (Fisher Scientific, 11558616) was used at 50, 75 or 100 ug/ml while hygromycin B (Thermo Fisher Scientific, 10687010) was used at 30, 40 or 50 μg/ml and single colonies were isolated 11-17 days after the nucleofection. 48-70 single colonies were picked for each targeting event and correctly targeted clones were first identified by junction-long PCR using the REDTaq PCR reaction mix (Sigma R2523) or the Phusion High-Fidelity PCR Master Mix with GC Buffer (Thermo Scientific, 10094387). Junction-long PCR (LPCR) was performed using primers residing outside of the respective HAs (Table S3). LPCR products, from selected heterozygous clones and from both the targeted and wt alleles, were Sanger sequenced (Microsynth) using custom-designed internal primers which resulted in overlapping sequencing reads so that the full extent of the insert was read. Two or three heterozygous clones of each targeting event, fully verified by junction PCR and Sanger sequencing, were further processed for the removal of the selection cassette. Cells were dissociated in single-cell suspension by TrypLE Express and 200K cells were nucleofected in 20 μl total volume containing 1 ug of the pCAGGS-Flpo-IRES-Puro plasmid ^50^ (gift from K. Anastassiadis, TU Dresden). Cells were seeded in three different concentrations (1:30, 1:10 and the rest) into three wells of a 6-well plate. Puromycin (Gibco, A1113803) was added one day after the nucleofection at 1 µg/ml for one day. On days 11-14 after the nucleofection, 6-10 clones for each independent initial clone were picked and analyzed for the removal of the selection cassette by genomic PCR. The correctness of the targeted and wt alleles was then verified by junction LPCR and Sanger sequencing as explained above. The sequences of the targeted alleles, including the 5’ and 3’ homology arms, have been submitted to GenBank with accession numbers OR729124 (*INS ^eGFP^*), OR727484 (*GCG ^mCHERRY^*), OR727485 (*MAFA ^mCHERRY^*) and OR727486 (*INS ^GCaMP^*^6^). Two independent clones, free of the selection cassette and sequence-verified for both alleles, were expanded and cryopreserved and one of them (Table S4) was used for the subsequent experiments described here. DNA was isolated using the Quick-DNA Miniprep Plus Kit (Zymo Research, D4069), LPCR was performed using LongAmp® Taq 2X Master Mix (NEB, M0287L), Sanger sequencing by Microsynth and alignments were performed using SnapGene or Geneious Software.

### Differentiation of hPS cell lines into SC-islets

H1 parental and modified hES cells were differentiated into PP cells using the STEMdiff™ Pancreatic Progenitor Kit (STEMCELL Technologies, 05120) according to the manufacturer’s instructions or using an adaptation of published procedures ^8,10,51^ with changes as described (Table S5). For the first procedure, hES cell colonies were dissociated into single cells using TrypLE Express (Gibco, 12604-013) and seeded on Matrigel-coated plates as described above at a concentration of 95K cells/cm^2^ in mTeSR1 supplemented with 20 µM ROCKi. The medium was replaced the next day by mTeSR1 and differentiation was initiated by replacing it with the S1d1 differentiation medium (Table S5) when cells were 60-70% confluent, typically two days after the initial seeding. Daily washes with DPBS (Gibco, 14190250) and media changes were done until S4d5 when the cells reached the end of the PP stage. For the second procedure, 220K cells/cm^2^ in mTeSR1 supplemented with 20 µM ROCKi were seeded, the medium was replaced the next day by mTeSR1 and differentiation was initiated six hours later at a confluence of 90% by replacing it with the S1d1 differentiation medium (Table S5). The monolayer of PP cells was dissociated using TrypLE Express (Gibco, 12604-013) at 37°C for 2-3 mins and seeded in microwells of AggreWell 800 plates (STEMCELL Technologies, 34815), using the recommended procedure by the supplier and at a density of 5000 cells per microwell. Media for S5-S7 were also an adaptation of published procedures and half of the medium was changed daily ^8,10,15,51^ (Table S5). S7 medium was modified by supplementing it with 1 x N21-MAX insulin-free supplement (R&D SYSTEMS, AR010) during S7 (S7D1-S7D14).

### Immunofluorescence analyses

For immunofluorescence (IF) on coverslips, cells were cultured on 12 mm diameter Matrigel-coated coverslips (Carl Roth, P231.1) placed in 24-well wells (Corning, CLS3524) and differentiated as described above. Cells were then fixed in 4% paraformaldehyde (PFA) for 20 min at RT and washed with PBS. Cells were blocked and permeabilized for 1 hr at RT using a 5% serum / 0.3% Triton X-100 PBS solution. Samples were then incubated overnight under gentle rotation at 4°C in 2.5% serum / 0.3% Triton X-100 in PBS containing the primary antibodies in the appropriate concentration (Table S6). The following day, samples were incubated for 1 hr at RT in 2.5% serum / 0.3% Triton X-100 in PBS containing the appropriate secondary antibodies, conjugated with either Alexa 488, 568 or 647, at a 1:500 dilution (Table S7). The coverslip with the stained cells was then placed on a microscope slide, covered with ProLong™ Gold Antifade mounting medium with DAPI (Invitrogen, P36931) and overlayed with a rectangular coverslip. Due to the multiple steps some cells would detach resulting in occasional patches on the coverslip that would be devoid of cells.

For immunofluorescence on cryosections, SC-islets were fixed in 4% PFA for 1h at 4 °C, washed once in PBS, resuspended in a 20% sucrose / PBS solution and kept at 4 °C overnight. SC-islets were then resuspended in OCT (Sakura, 4583), moved into a mold, frozen on dry ice and kept at -80 °C. The OCT bocks were cryosectioned at a thickness of 8 μm on a glass slide (Thermo Fischer Scientific, 15438060) and stored at -80 °C. Before staining, the glass slides were left to warm up and dry at RT for approximately 5-10 min. The slides were quickly washed three times with PBS. SC-islets were permeabilized with 0.3% Triton X-100 PBS for 30 min at RT and then blocked for 1 hr at RT with 5% serum / 0.3% Triton X-100 PBS. Slides were incubated at 4 °C with the primary antibodies, appropriately diluted (Table S7) in PBS containing 2.5% fetal bovine serum and 0.3% Triton X-100. The following day, slides were incubated for 1 hr at RT in the dark with the appropriate secondary antibodies (Table S8), diluted 1:500 in PBS containing 2.5% fetal bovine serum and 0.3% Triton X-100. After extensive PBS washes, the slides were mounted with ProLong Gold Antifade with DAPI (Thermo Fischer Scientific, P36931) and covered with a glass coverslip (Engelbrecht, K12450).

IF images were acquired using a Zeiss Axio Observer Z1 microscope coupled with the Apotome 2.0 imaging system and with consistent exposure times for the Alexa 488, 568 and 647 channels in between passages and conditions to allow for direct comparison of the signal intensities.

### Flow cytometry (FC) and fluorescence-activated cell sorting (FACS) of hPS cell-derived cells

Cells were first washed with PBS and then dissociated into single cells using TrypLE Express (Gibco, 12604-013). Following dissociation, the cells were counted using the Countess II Automated Cell Counter (Thermo Fischer Scientific) and Trypan Blue. Then, cells were washed with PBS and fixed at a concentration of 4 million/ml in 4% PFA for 10 min at 4°C. For staining with transcription factor antibodies, cells were washed with PBS and permeabilized using the Foxp3 Transcription Factor Staining Buffer Set (Invitrogen, 00-5523-00) for 1 h in the dark at 4°C. Cells were then washed again using the 1x permeabilization buffer. Thereafter, 2x10^6^ cells/sample were blocked using 100 μl of 5% serum in 1x permeabilization buffer and then incubated with primary antibodies at the appropriate concentration (Table S6) in the same buffer overnight at 4°C at 1200 rpm. Then, they were washed twice with 1x permeabilization buffer and incubated with secondary antibodies at the appropriate concentration (Table S7) in 100 μl of 1x permeabilization buffer containing 5% serum at RT for 1 hour at 1200 rpm in the dark. For cytoplasmic factors, 2x10^6^ cells/sample were blocked for 30 mins at RT in 100 μl PBS containing 2% serum and 0.3% Triton X-100. Then cells were washed with 0.1% Triton X-100 in PBS solution and incubated at 4°C with primary antibodies at the appropriate concentrations (Table S6) in the same buffer overnight. Thereafter, cells were washed twice with 0.1% Triton X-100 in PBS solution and incubated with secondary antibodies at the appropriate concentration (Table S7) at RT in the dark for 1 hour. FC data were acquired using BD FACSCanto™ II and analyzed using the FlowJo software.

For live cell analyses, SC-islets were washed with PBS and then dissociated into single cells with a solution of 1x non-enzymatic cell dissociation solution (Sigma, C5789), DNase I (40 μg/ml) (Roche, 11284932001) and 0.01% Trypsin-EDTA (Invitrogen, 15400054). Dissociation takes generally up to 6 min in a 37 °C water bath. Every 2 min, the clusters were pipetted to break them up and after the 6 min, the process was stopped by adding 2% BSA in DMEM/F-12. Cells were washed with PBS, spun down and resuspended in 350 μl of PBS along with 0.05% FCS and 0.1 mM EDTA. DRAQ7 (Biostatus, DR70250) was added to cell suspension at a final concentration of 0.8 μM and data were acquired using the BD FACSCanto™ II. For FACS, cells were dissociated as above and resuspended in 350 μl of PBS containing 0.05% FCS and 0.1 mM EDTA. DRAQ7 was added to the cell suspension and cells were sorted using the BD FACS Aria III and collected in RLT buffer of the RNeasy kit (Qiagen, 74004) for RNA isolation and RT-qPCR analyses.

### Static glucose-stimulated insulin secretion (GSIS) and insulin content assays

At the end of S7 (S7d14 – S7d16) 150 clusters were collected, washed twice with fresh pre-warmed (37°C) Krebs-Ringer solution buffered with HEPES (KRBH) (2.5 mM CaCl_2_, 129 mM NaCl, 4.8 mM KCl, 1.2 mM MgSO_4_, 1.2 mM KH_2_PO_4_, 1 mM Na_2_HPO_4_, 5 mM NaHCO_3_, 10 mM HEPES, 0.1% BS, pH adjusted to 7.4 with 5 M NaOH) and then incubated for 1 h in 2 ml of KRBH as a starvation step. After starvation, clusters were incubated with low (2.8 mM) glucose KRBH for 1 h, then with high (16.7 mM) glucose KRBH for 1 h and finally in KCl / high glucose (30 mM / 16.7 mM) KRBH for 1 h. Supernatants were collected after each incubation and -frozen at 80 °C. Cell pellets were immediately used for insulin isolation using the acid ethanol extraction method ^52^ and genomic DNA isolation using the DNeasy Blood & Tissue kit (Qiagen, 69504). The human C-peptide enzyme-linked immunosorbent assay (ELISA) (Mercodia, 10-1141-01) or the human insulin homogeneous time-resolved fluorescence (HTRF) assays (Cisbio, 62IN2PEG) were used to determine C-peptide and insulin, respectively. DNA quantification was done using a Nanodrop spectrophotometer and the insulin content of the SC-islets was normalized to the corresponding DNA content.

### RNA isolation and RT-qPCR

Total RNA was prepared using the RNeasy kit with on-column genomic DNA digestion (Qiagen, 74004) following the manufacturer’s instructions. First-strand cDNA was prepared using the TAKARA PrimeScript RT Master Mix (TAKARA RR036A). Real-time PCR primers (Table S8) were designed using the Primer 3 software (SimGene), their specificity was ensured by *in silico* PCR and they were further evaluated by inspection of the dissociation curve. Reactions were performed with the FastStart Essential DNA Green Master mix (Roche 06924204001) using the Roche LightCycler 480 and primary results were analyzed using the onboard software. Reactions were carried out in technical triplicates from at least three independent biological samples. Relative expression values were calculated using the *TBP* housekeeping gene and the ΔΔCt method by normalizing to the expression levels of the corresponding undifferentiated hPS cells.

### Live Ca^2+^ imaging

Before imaging, *INS^GCaMP6^* SC-islets were embedded in a fibrinogen gel for stabilization. Briefly, embedding was done by first washing SC-islets in the HBSS buffer. SC-islets were then submerged in 2 µl fibrinogen (100 mg/ml) (Sigma, F8630) and this 2 µl volume was then applied to a mixture of 6 µL HBSS and 2 µL thrombin (Sigma, T7513). Ca^2+^ imaging was performed on an LSM780 in an open bath chamber (JG-23, Warner Instruments) using a 20x objective (W Plan-Apochromat 20x/1,0 DIC M27 70 mm). Islets were perfused with KRBH buffer (137mM NaCl, 5.36mM KCl, 0.34mM Na_2_HPO_4_, 0.81mM MgSO_4_, 4.17mM NaHCO_3_, 1.26mM CaCl_2_, 0.44mM KH_2_PO_4_, 10mM HEPES, 0.1% BSA, pH 7.3) containing 3mM glucose (resting) or 16.7mM glucose + 60mM KCl (depolarizing condition) using a peristaltic pump at a flow rate of 0.5 ml/min. 60, rather than 30 mM KCl has been used because previous experiments with native islets showed a slightly higher peak response to 60 mM KCl, therefore making the peak identification easier. Images were taken at a resolution of 512 x 512 pixels at 0.2 frames/second (z-stack) or 2 frames/second (single plane). Image analysis and assessment of traces were performed using Fiji. Registration of the time series images was performed using the descriptor-based registration plugin and intensity traces of single *INS^GCaMP6+^* cells were collected after manual ROI contouring and the use of the Time Series Analyzer V3 plugin. Traces were then normalized to basal activity F_0_ (F_0_ = mean intensity value of the first 2 minutes at 3 mM glucose).

### Automated confocal imaging and analysis

SC-islets were distributed on GRID3 96-well plates (Sun Biosciences) and stained with 10 mg/ml Hoechst 33342 dye (ThermoFisher H1399) overnight at 37 C and 5% CO2. Samples were acquired using the automated confocal CV7000 Yokogawa platform with a 0.75 NA 20X objective and appropriate lasers and filters for the fluorescent markers: Hoechst 33342 for the nuclei, GFP and RFP for the cellular markers. A Z volume of 120 microns, taking one image every 20 microns, was probed. Images were analyzed in a multistep approach: Stardist was used to analyze the confocal images by initially identifying spots of very strong DAPI signal which were computationally expanded until they met a neighboring expanding area and this produced a segmentation pattern with each segment approximating the cell area which was then scored for GFP or RFP signal. The labeled cell areas were then imported in a Cell Profiler pipeline, with which the SC-islets were identified, the cell nuclei were related to the appropriated SC-islet and the intensity profiles of the SC-islets and the cells were measured in the GFP and RFP channels ^53^. The pipeline was adjusted to allow parallelization of the calculations and as such it will apply to high content high throughput analyses with minimal adjustments. The resulting data was analyzed and plotted using KNIME (https://www.knime.com).

### Statistical analysis and graphs

Data were analyzed for statistical significance and plotted using Prism 9.

## RESULTS

### Generation of hES cell reporter lines using CRISPR/Cas9 mediated homologous recombination

To ensure the specificity of the reporter lines we opted for a knock-in strategy via homologous recombination so that the reporters would be expressed under the control of all the regulatory elements of the cognate gene. Since homologous recombination in human ES cells is extremely inefficient ^54^, we used CRISPR/Cas9 mediated DNA cleavage to enhance the incorporation of the transgenes in the target loci ^55^. To minimize disruption of the targeted gene function, the coding sequence was fully preserved and the homologous arms were designed so that a T2A self-cleaving peptide sequence ^56^ would be introduced in frame with the last amino acid codon of the endogenous gene and in-frame with the first codon of the following reporter. The mCHERRY cDNA carried nuclear localization signals whereas the eGFP and GCaMP6 cDNAs encoded cytoplasmic proteins. The reporter sequence was followed by a resistance selection cassette flanked by the flippase recognition target (FRT) sites ^57^ (Figure 1A). The targeting vectors were assembled using the Gibson method ^58^ and DNA fragments were commercially synthesized or isolated from available vectors ^43–45^ by PCR or conventional restriction enzyme digestion (Table S1).

**Figure 1.**
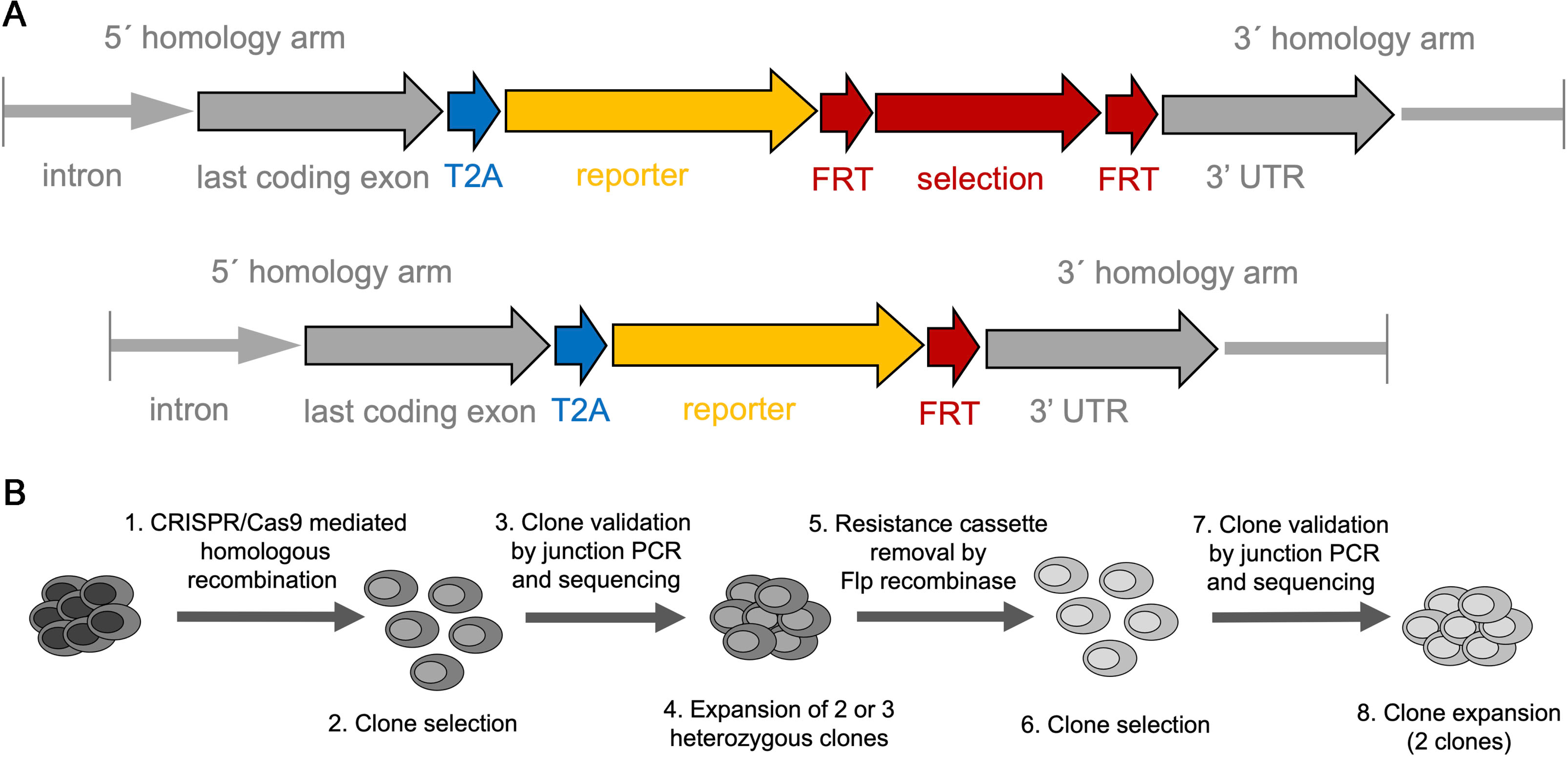
Strategy for the derivation of the reporter lines. (A) The general structure of the targeted loci is shown before and after the removal of the selection cassette by Flp recombinase. Homology arms used for the targeting are shown in grey. The last coding exon of the gene of interest (grey) lacks the STOP codon and is in frame with the self-cleavage peptide T2A sequence (blue) and the reporter cDNA (yellow). These are followed by the selection cassette (red) flanked by FRT sites (red). After removal of the selection cassette, only one FRT site remains upstream of the 3’UTR of the targeted gene (grey). (B) Outline of the procedures for the reporter line generation. H1 hES cells were targeted using CRISPR/Cas9 mediated homologous recombination, resistant clones were selected, validated by junction PCR and sequencing and correctly targeted heterozygous clones were expanded (1-4). The selection cassette was then removed by transient transfection of Flp recombinase and selected through the transient expression of a drug-resistance gene. Selected clones were confirmed by junction PCR and sequencing for both the modified and targeted allele.

To enhance the targeting specificity, two, rather than one, sites within 50 bp of the STOP codon were targeted with either nickase CRISPR/Cas9 ^47^ or wt CRISPR/Cas9 ^47^ (Table S2). H1 cells that show very good efficiency towards endoderm differentiation were then nucleofected with the donor vector and sgRNAs (Table S2) as well as either a nickase CRISPR/Cas9 encoding plasmid ^47^ or wt CRISPR/Cas9-NLS protein (NEB). Following antibiotic selection, resistant clones were analyzed by junction-long PCR using primers (Table S3) outside the homology arms (HAs) and Sanger sequencing of the entire product(s) for both the targeted and wt alleles. Two correctly targeted, heterozygous clones, were then nucleofected with FRT recombinase for the deletion of the selection cassette and subclones from each of the initial parental clones were again analyzed by junction-long PCR and Sanger sequencing genotyping for both the targeted and wt alleles as above. The deletion of the selection cassette was necessary to ensure that the strong promoter driving expression of the resistance marker would not interfere with the expression of the locus. Two independently derived and sequence-verified clones were then expanded and cryopreserved (Figure 1B). One of them (Table S4) was used for the subsequent differentiation experiments described below. The *GCG* and *MAFA* loci were targeted using the *INS ^eGFP^* reporter line to generate the *INS ^eGFP^/ GCG ^mCHERRY^* and *INS ^eGFP^/ MAFA ^mCHERRY^* double reporter lines, respectively. Detailed maps of the resulting targeted loci, including the 5’ and 3’ HAs, are provided (Figure S1A-D) as well as the corresponding annotated sequences (Files S1-4). To perform the experiments that follow, we used a single clone for each reporter line (Table S4) which was also karyotyped to exclude the appearance of chromosomal abnormalities (Figure S2A-E). The pluripotency of each used cell clone was assessed qualitatively by immunofluorescence for the expression of the pluripotency markers SOX2, OCT4 and SSEA4 (Figure S2F-J) and quantitatively by flow cytometry for OCT4 / SOX2 (Figure S2K-P). As controls, hES cells of the corresponding cell line were stained with secondary antibodies. No differences were seen between the parental H1 line the derived reporter lines with respect to OCT4 / SOX2 expression.

### The reporter lines differentiate efficiently into definitive endoderm and pancreatic progenitor cells

The hES lines were differentiated into pancreas progenitors (PP) and we compared the differentiation efficiency of the generated reporter lines with the parental hPS cell line H1, initially towards definitive endoderm (DE) cells and subsequently into PP cells. DE differentiation was initially assessed by immunofluorescence for the expression of SOX17 and FOXA2, two transcription factors essential for the specification of DE (D’Amour et al. 2005). These experiments suggested that the derived reporter lines differentiated into DE with similar efficiency to the parental hES cell line (Figure 2A-E). These results were confirmed by flow cytometry for SOX17 and FOXA2, which showed that the generated reporter lines maintained similar capacities to differentiate into DE ranging from 81% to 99 % SOX17^+^/FOXA2^+^ cells (Figure 2F-J, representative control in S2Q).

**Figure 2.**
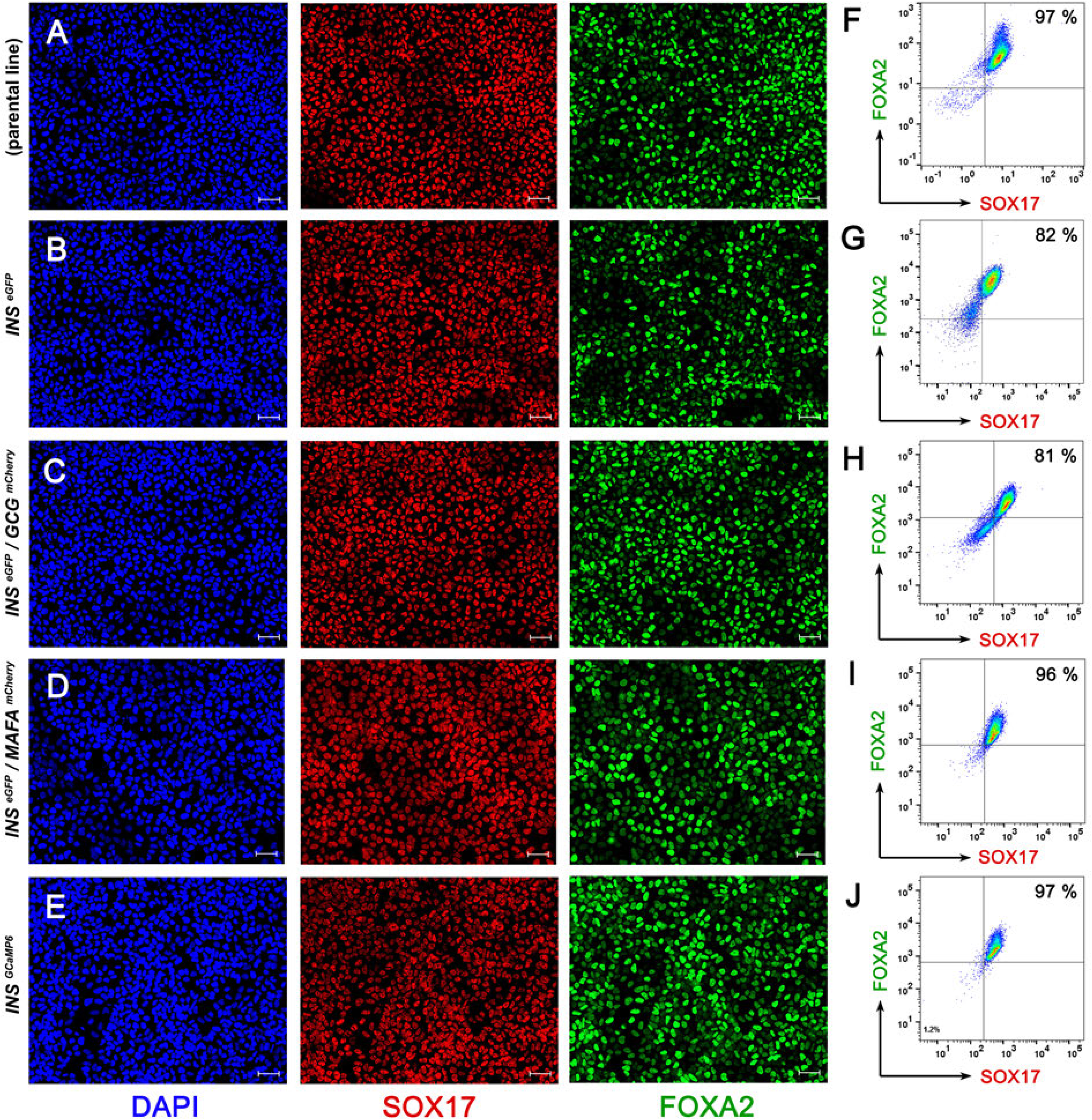
Reporter lines differentiate efficiently into definitive endoderm. (A-E) Representative images of immunofluorescent staining of DE cells derived from H1 (parental line) (A), H1 *INS ^eGFP^* (B), H1 *INS ^eGFP^*/*GCG ^mCherry^* (C), H1 *INS ^eGFP^*/*MAFA ^mCherry^* (D) and H1 *INS ^GCaMP6^* (E) hES cells for the expression of the DE transcription factors SOX17 (red) and FOXA2 (green). (F-J) Flow Cytometry analyses of DE cells derived from H1 (parental line) (F), H1 *INS^eGFP^* (G), H1 *INS ^eGFP^*/*GCG ^mCherry^* (H), H1 *INS ^eGFP^*/*MAFA ^mCherry^* (I) and H1 *INS ^GCaMP6^* (J) hES cells for the expression of SOX17 and FOXA2 showed that between 81% and 97% of the cells were SOX17^+^/FOXA2^+^. Scale bars correspond to 50 µm.

DE cells derived from H1 cells and the reporter lines were then sequentially differentiated into primitive gut tube (PGT) cells, posterior foregut (PF) cells and finally into PP cells. PP cells were initially analyzed by immunofluorescence, flow cytometry analyses and RT-RT-qPCR for expression of *PDX1*, *SOX9* and *NKX6.1*. These transcription factors are essential, during development, for the specification of PP cells and their competence to subsequently differentiate into pancreatic endocrine cells ^59–61^ .

Immunofluorescence analyses suggested similarly high levels of conversion of all lines into PDX1^+^/SOX9^+^/NKX6-1^+^ PP cells ^62^ (Figure 3A-E). This was further supported by flow cytometry analyses in which PDX1^+^/SOX9^+^ cells were first identified based on the gates set by the corresponding control stainings (Figure S3A). Τhen NKX6.1^+^ cells were scored in these pre-gated PDX1^+^/SOX9^+^ cells based on the gate set by the corresponding control staining (Figure S3B). As controls, PP cells of the corresponding cell line were stained with secondary antibodies. In independent differentiations, the percentage of PDX1^+^/SOX9^+^ cells ranged from 94 % to 99 % (Figure S3C-H, control in S3A) and the percentage of PDX1^+^/SOX9^+^/NKX6-1^+^ cells ranged between 42 % and 74% (Figure 3F-K, control in S3B). As a result of the gating strategy, there are no cells in the lower two quadrants in Figure 3F-J.

**Figure 3.**
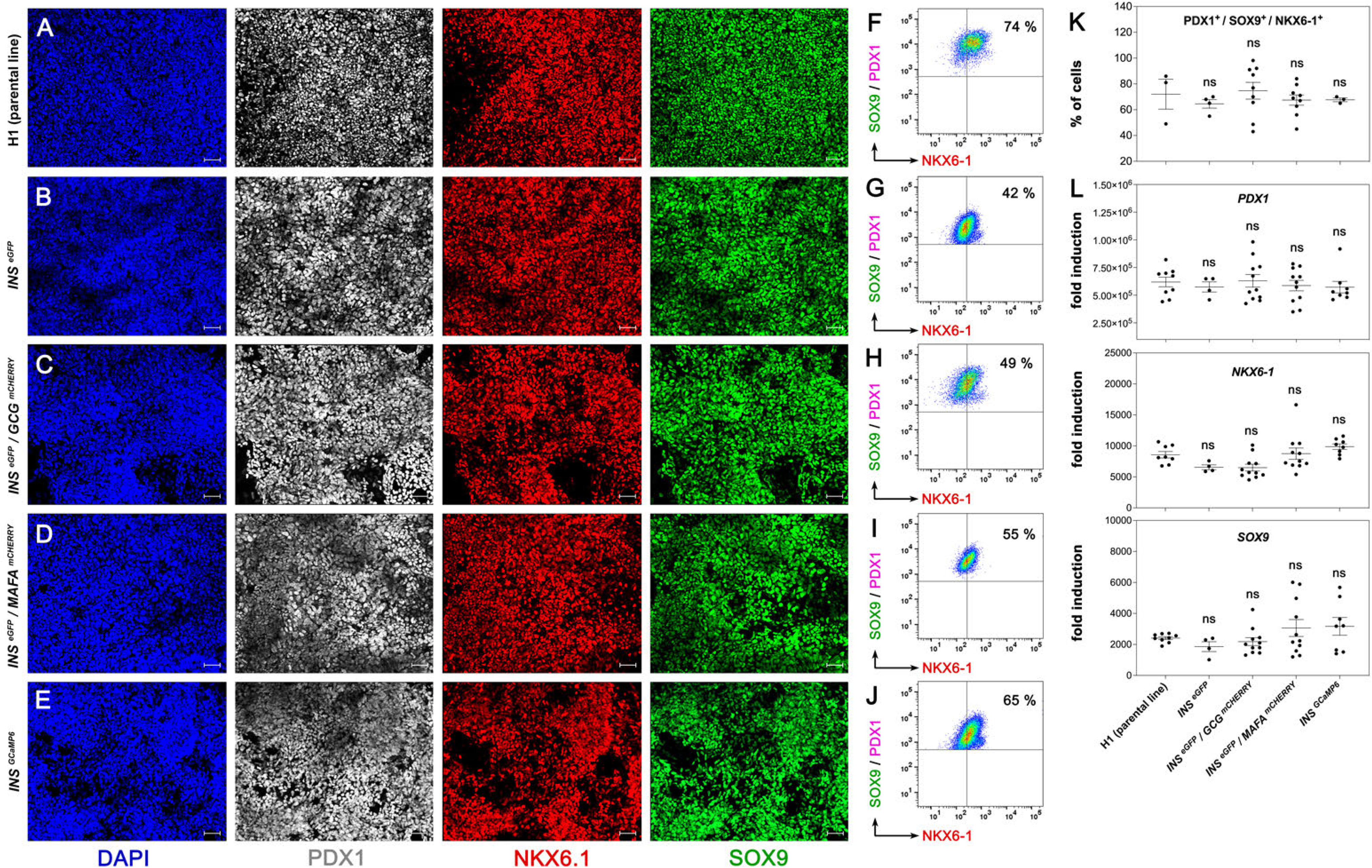
Reporter lines differentiate efficiently into pancreatic progenitors. (A-E) Representative images of immunofluorescent staining of PP cells derived from H1 (parental line) (A), H1 *INS ^eGFP^* (B), H1 *INS ^eGFP^*/*GCG ^mCherry^* (C), H1 *INS ^eGFP^*/*MAFA ^mCherry^* (D) and H1 *INS ^GCaMP6^* (E) hES cells for the key PP transcription factors PDX1 (grey), NKX6.1 (red) and SOX9 (green). (F-J) Selected cytometry analyses of PP cells derived from H1 (parental line) (F), H1 *INS ^eGFP^* (G), H1 *INS ^eGFP^*/*GCG ^mCherry^* (H), H1 *INS ^eGFP^*/*MAFA ^mCherry^* (I) and H1 *INS ^GCaMP6^* (J) hES cells for the expression of PDX1, SOX9 and NKX6.1. (K) Summary of flow cytometry analyses of PDX1^+^ / SOX9^+^ / NKX6.1^+^ PP cells derived from the H1 (parental line), H1 *INS ^eGFP^*, H1 *INS ^eGFP^*/*GCG ^mCherry^*, H1 *INS ^eGFP^*/*MAFA ^mCherry^* and H1 *INS ^GCaMP6^* hES cells. Dots represent values from independent differentiation experiments. (L) Gene expression analyses by RT-qPCR for key pancreatic progenitor markers in PP cells derived from H1 (parental line), *INS ^eGFP^*, *INS ^eGFP^*/*GCG ^mCherry^*, *IN S^eGFP^*/*MAFA ^mCherry^* and *INS ^GCaMP6^* hES cells. Expression levels are expressed as fold induction relative to those in undifferentiated hES cells. Dots represent values from independent differentiation experiments Statistical analyses were performed by the non-parametric Kruskal-Wallis test using the H1 (parent line) as the control for the comparison with *p* ≤ 0.05 (*), *p* ≤ 0.005 (**) and *p* ≤ 0.0005 (***). Horizontal lines in the graphs represent the mean ± standard error of the mean (SEM). Scale bars correspond to 50 µm.

Gene expression analyses by RT-qPCR suggested that there were no statistically significant differences between the parental H1 line and the different reporter lines in the expression of *PDX1*, *SOX9* and *NKX6-1* (Figure 3L). Expression of *PTF1A*, a transcription factor expressed at the PP stage, did not appear to be significantly different between the H1 parental line and the derived reporter lines (Figure S3I). We then examined the expression of the alternative fate liver and gut markers, *AFP* and *CDX2* respectively. AFP expression was significantly higher only in the *INS ^eGFP^ / MAFA ^mCHERRY^*, but, in balance, *CDX2* expression was significantly lower in the same line. There were no significant changes in any of the other cell lines (Figure S3I).

Overall, these results suggested that the derived reporter lines differentiated into PP cells with similar efficiency to that of the parental H1 line.

### Differentiation into SC-islets and functionality of the reporter lines

PP cells derived from the H1 hES cell line and all reporter lines were clustered in microwells and further differentiated, in several independent experiments for each line and using an adaptation of published media ^8,10,51^, successively into pancreatic endocrine progenitors (PEP), immature SC-islets (Stage 6) and SC-islets (Stage 7) (Table S5).

At the PEP stage, the expression of *PDX1* and *SOX9* remained at similar levels as that at the PP stage for all lines whereas, as expected, the expression of *NKX6.1* increased substantially compared to the PP stage (Figure 3L and Figure S3J). There were no statistically significant differences between the expression of these genes in the PEP cells derived from the H1 parental line and those derived from the reporter lines (Figure S3K). Expression of the endocrine lineage inducing *NEUROG3* transcription factor ^63^ and its downstream target *NEUROD1* transcription factor ^64^ was readily detectable in both the H1 parental line and derived reporter lines with no detectable differences across the different lines (Figure S3K). In line with all the other markers, expression of the respective liver and gut lineage markers, *AFP* and *CDX2*, appeared very similar in the parental and all derived reporter lines (Figure S3K). Using the *INS ^eGFP^/ GCG ^mCHERRY^* reporter line and whole-mount immunofluorescence imaging we found that there was no expression of either the *INS ^eGFP^* or the *GCG ^mCherry^* reporter at the PP stage. Reporter expression was gradually increased through the PEP stage and S6, reaching maximum expression at S7 (Figure S3L).

Then, we assessed the differentiation efficiency of the reporter lines into SC-islets and β-cell functionality of the generated SC-islets. We first evaluated the relative differentiation efficiency of the reporter lines by RT-qPCR analyses. Expression levels of the key transcription factors *PDX1* and *NKX6-1* were similar in all lines (Figure 4A). The expression of the hormone genes *INS*, *GCG* and *SST* was variable among independent experiments but there were no statistically significant differences between the reporter lines and the parental line (Figure 4A, S4A). Genes expressed specifically in β-cells, and important for their function, were also generally expressed at similar levels in H1 and reporter line-derived SC-islets except for lower *GK* expression in the *INS ^eGFP^* SC-islets and lower *PCSK1* expression in the *INS ^eGFP^/ GCG ^mCHERRY^* and *INS ^GCaMP6^* SC-islets (Figure 4A, S4A). Overall, the extensive RT-qPCR analyses at this stage suggested that reporter lines differentiated with comparable efficiencies with one another and with the H1 parental line. We then compared the functionality of β-cells in SC-islets derived from the parental H1 line and the reporter lines by static glucose stimulation insulin secretion (GSIS) assays. The stimulation indexes in both cases were very similar between H1- and reporter line-derived SC-islets (Figure 4B).

**Figure 4.**
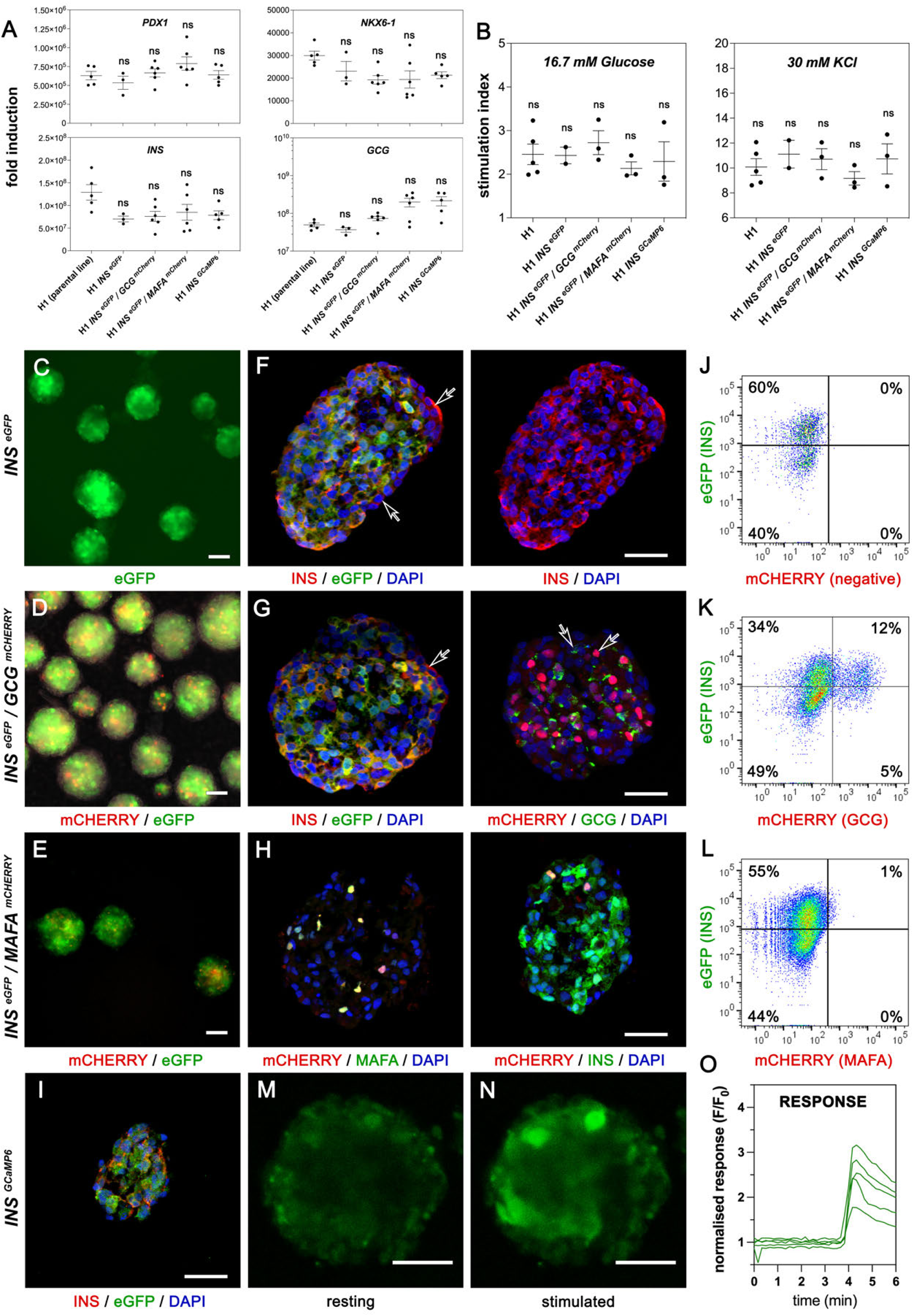
Reporter lines differentiate efficiently into SC-islets and reporters are functional. (A) Gene expression analysis by RT-qPCR of SC-islets derived from all the reporter lines and compared to H1-derived SC-islets for the β-cell markers *PDX1, NKX6.1, INS* and *GCG*. Gene expression levels are expressed as fold induction relative to those in undifferentiated hES cells. Dots represent values from independent differentiation experiments. (B) Static GSIS assays measuring C-peptide secretion after stimulation with high glucose (16.7 mM) and high glucose with 30 mM KCl from SC-islets derived from H1 (parental line), *INS ^eGFP^*, *INS ^eGFP^*/*GCG ^mCherry^*, *INS^eGFP^*/*MAFA ^mCherry^* and *INS ^GCaMP6^* hES cells. The stimulation index is determined by the ratio of secretion under these conditions to secretion in basal low glucose (2.8 mM). Dots represent values from independent experiments. (C-E) Whole mount imaging of GFP (green) and mCherry (red) fluorescence from live SC-islets from *INS ^eGFP^* (C), *INS ^eGFP^*/*GCG ^mCherry^* (D) and *INS ^eGFP^*/*MAFA ^mCherry^* (E) hES cells. (F-I) Representative images of immunostainings on cryosections showing extensive co-localization of the reporters with the corresponding genes in for *INS ^eGFP^* (F), *INS ^eGFP^*/*GCG ^mCherry^* (G), *INS ^eGFP^*/*MAFA ^mCherry^* (H) and *INS ^GCaMP6^* (I) SC-islets. GFP expression was visualized through immunostaining whereas mCherry expression was visualized either directly (*GCG ^mCherry^*) or following immunostaining (*MAFA ^mCherry^*). Occasional non-coinciding stainings are indicated with open arrows (F, G). (J-L) Live flow cytometry analyses for reporter expression to detect GFP^+^ cells from *INS ^eGFP^* SC-islets (J), GFP^+^ and mCHERRY^+^ cells from *INS ^eGFP^*/*GCG ^mCherry^* SC-islets (K) as well as GFP^+^ and mCHERRY^+^ cells from *INS ^eGFP^*/*MAFA ^mCherry^* SC-islets (L). (M-O) Perfusion live imaging of *INS ^GCaMP6^* SC-islets showed low fluorescent signal at low glucose (2.8 mM) (M) but a strong signal in response to 16.7 mM Glucose + 30 mM KCl (N). Average traces of responding cells in individual *INS ^GCaMP6+^* SC-islets (n=5) are shown (O). Statistical analyses were performed by the non-parametric Kruskal-Wallis test using the H1 (parent line) as the control for the comparison with *p* ≤ 0.05 (*), *p* ≤ 0.005 (**) and *p* ≤ 0.0005 (***). Horizontal lines on the graphs represent the mean ± SEM. Scale bars correspond to 50 μm.

We then assessed the functionality and faithfulness of the reporters at the SC-islet stage by whole-mount live fluorescence, immunofluorescence on cryosections, flow cytometry and RT-qPCRs on FACS-isolated cells. Both cytoplasmic eGFP and nuclear mCHERRY fluorescence were strong and easily detectable in whole SC-islets of the *INS ^eGFP^*, *INS ^eGFP^/ GCG ^mCHERRY^* and *INS ^eGFP^/ MAFA ^mCHERRY^* lines with the partial exception for *MAFA ^mCHERRY^* fluorescence which was weaker due to the very small number of MAFA^+^ cells (Figure 4C-E, see also below). The concordance of reporter expression with the corresponding protein was confirmed by double immunofluorescence of the protein and the corresponding reporter on cryosections. Importantly, this was also confirmed for the rare MAFA^+^ cells since all detected MAFA^+^ cells were also mCHERRY^+^ (Figure 4F-I, S4B, C). There were rare instances where cells appeared stained for the reporter but not for the corresponding protein and vice versa (Figure 4F, G open arrows). This could be due to staining and imaging limitations but it is worth noting that, at this stage, cells are still not fully mature and hormone expression is likely to fluctuate, leading to differences arising from differential protein stabilities. We also examined if these reporter lines could be used with live cells to provide a quantitative evaluation of the differentiation efficiency and β-cell functionality. SC-islets derived from the *INS ^eGFP^*, *INS ^eGFP^/ GCG ^mCHERRY^* and *INS ^eGFP^/ MAFA ^mCHERRY^* lines were dissociated into single cells and flow cytometry was used to evaluate the detection of INS^+^, GCG^+,^ INS^+^/GCG^+^, INS^+^/MAFA^+^ cells and MAFA^+^ cells. These experiments demonstrated that such cells were readily detectable using the live reporters (Figure 4J-L, control in S4D). The functionality of the GCaMP6 reporter in the *INS ^GCaMP6^* line was assessed using SC-islets in perfusion live imaging assays where SC-islets in resting conditions (3 mM Glucose) were challenged with 16.7 mM Glucose + 60 mM KCl. The fluorescence of individual cells was traced demonstrating a strong increase upon KCl stimulation (Figure 4L-N, Video S1) which was found to be detectable in 67.6% ± 9.3% within *INS^GCaMP6+^* cells (n= 5 individual SC-islets, 203 cells in total)(controls in Figure 5B, C, G, and H).

**Figure 5.**
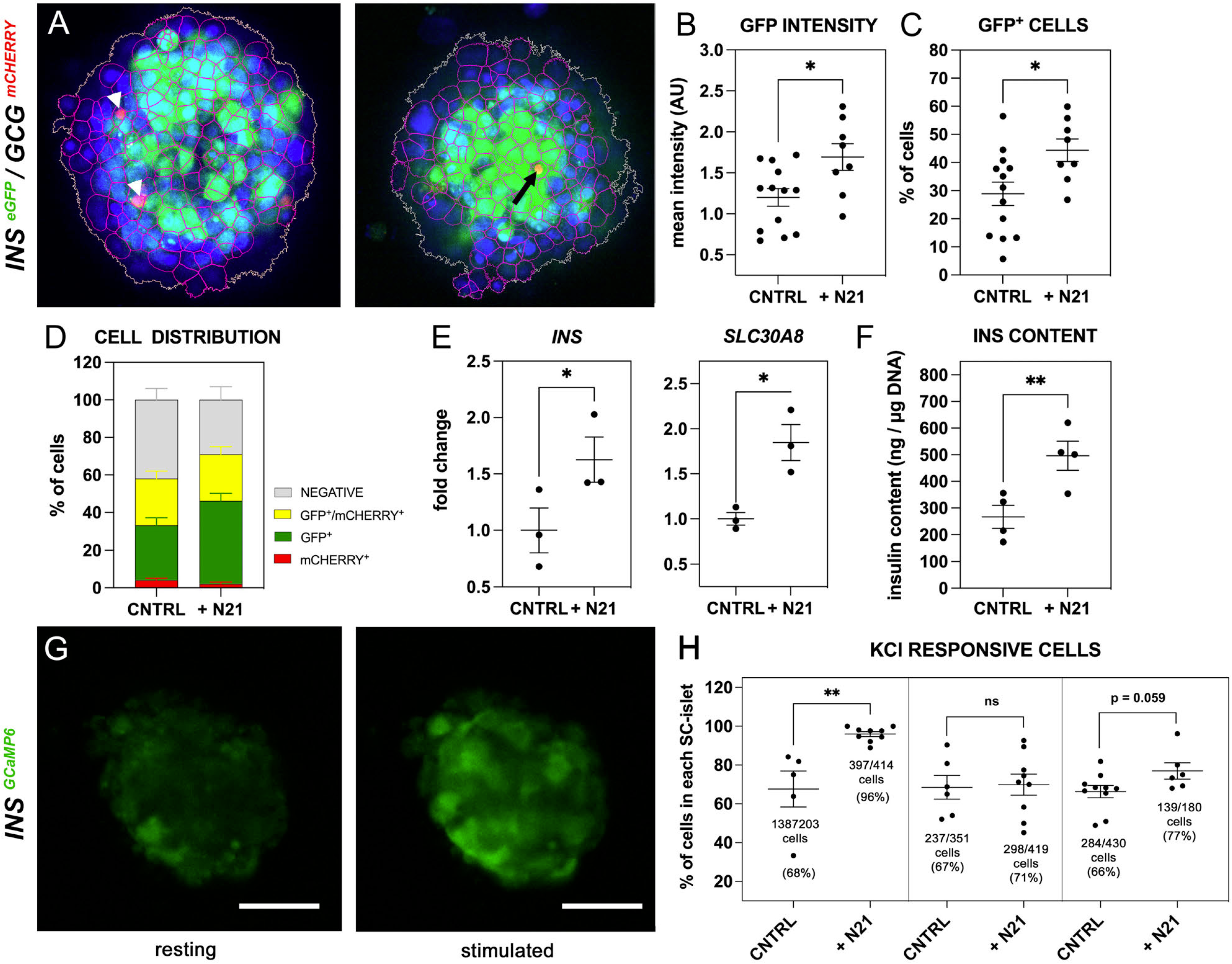
N21 promotes β-cell formation. (A) Cell segmentation of confocal images to identify negative, GFP^+^, mCHERRY^+^ (arrowheads) and mCHERRY^+^/GFP^+^ (arrow) individual cells. (B-D) Mean intensity per SC-islet (B), % of β-cells per islet (C) and distribution of SC-islets (D) (n=8) after N21 treatment during S7. Dots in B, C correspond to individual SC-islets. (E) Changes in the expression of β-cell genes *INS* and *SLC30A8* upon supplementation of S7 medium with N21 as compared to control conditions (CNTRL). Dots represent values from independent differentiation experiments. (F) DNA normalized insulin content of SC-islets differentiated under control (CNTRL) and N21 supplemented S7 conditions and determined following static GSIS assays. Dots represent values from independent differentiation experiments. (G, H) Snapshots from the perifusion live imaging of *INS ^GCaMP6^* SC-islets, differentiated in the presence of N21 during S7, at low glucose (3mM) (E) and in response to 16.7 mM Glucose + 60 mM KCl (G). The % of cells responsive to 16.7 mM Glucose + 60 mM KCl in individual *INS ^GCaMP6^* SC-islets are shown for control CNTRL and the N21 *INS ^GCaMP6^* SC-islets. Individual rectangles correspond to independent differentiation experiments and dots correspond to individual SC-islets. Statistical analyses were performed by the non-parametric Kruskal-Wallis test using the H1 (parent line) as the control for the comparison with *p* ≤ 0.05 (*), *p* ≤ 0.005 (**) and *p* ≤ 0.0005 (***). Horizontal lines on the graphs represent the mean ± SEM. Scale bars correspond to 50 μm.

In summary, these experiments suggested that all reporter lines maintain a similar capacity to the parental line in differentiating into SC-islets containing functional β-cells and that they can indeed be used in live assays to assess the efficiency of differentiation (*INS ^eGFP^*, *INS ^eGFP^/ GCG ^mCHERRY^*), maturation (*INS ^eGFP^/ MAFA ^mCHERRY^*) and functionality (*INS ^GCaMP6^*).

### The inclusion of N21 during S7 increases the number of monohormonal β-cells in SC-islets

We then examined whether we could use these reporter lines to identify conditions that may improve the differentiation of the SC-islets. N21 is a defined cell culture supplement that has been used extensively to promote the survival, maturation and function of neurons in culture as well as to improve 3D culture conditions ^65^. Pancreatic endocrine cells and neurons share functional similarities and thus we reasoned that N21 may have a beneficial effect on the differentiation of SC-islets. Accordingly, and to provide proof of concept that these reporter lines can be used to identify improved differentiation conditions, we used N21 as an additional supplement during S7 of the differentiation. To also establish a pipeline for future high-content analyses, to be used for the evaluation of distinct differentiation conditions, *INS ^eGFP^* / *GCG ^mCherry^* SC-islets were differentiated as single clusters, each in a separate microwell over 24 days (S5-S7) in Gri3D 96-well plates (Sunbioscience). To assess the effect of N21, on *INS* and *GCG* expression as well as on the segregation of INS^+^ and GCG^+^ cells in SC-islets, images were acquired using an automated confocal imaging platform. For each SC-islet, a z-volume of 120 microns was acquired and images were analyzed using a multistep approach to segment the images into individual cells and assign mCHERRY and GFP intensity values to each cell (Figure 5A).

The initial analysis showed that, upon N21 addition, there was a significant increase in the mean eGFP signal intensity per SC-islet. This was a specific β-cell effect because the mean mCHERRY signal intensity did not change (Figure 5B, S5A). Cell segmentation of control (n=13) and N21 differentiated (n=8) SC-islets revealed that this was accompanied by a significant 50% increase in the percentage of the single GFP^+^ cells (from 28.9 ± 4.2 to 44.4 ± 4.0) but no change in the percentage of single mCHERRY^+^ cells, or eGFP^+^/mCHERRY^+^ cells (Figure 5C, D, S5B). RT-qPCR analyses showed that *INS* and *SLC30A8* were upregulated, whereas the expression of *GCG* and *SOM* was not affected confirming that the N21 effects were indeed β-cell specific (Figure 5E, S5D). Interestingly, the expression of *PDX1* and *NKX6.1* was not affected (Figure S5D) suggesting that N21 affected specific aspects of the β-cell formation. *MAFA* expression was also not affected (Figure S5D) by the inclusion of N21 and therefore we did not use the *INS ^eGFP^/ MAFA ^mCHERRY^* reporter line to assess possible effects on the number of MAFA^+^ cells.

We then evaluated whether the insulin content of the SC-islets was increased and whether β-cell functionality changed following N21 treatment. Consistent with the findings described above, we found that insulin content increased nearly two-fold in N21-treated SC-islets derived from independent differentiations (n=4) (Figure 5F). On the other hand, static GSIS assays did not show increased insulin secretion in either high (16.7 mM) glucose concentration or additional high (30 mM) KCl concentration (Figure S5C). To assess whether N21 treatment resulted in changes in Ca^2+^ mobilization, we used *INS ^GCaMP6^* derived SC-islets from three independent differentiations and perifusion live imaging assays, during which SC-islets in resting conditions (3 mM Glucose) were challenged with 16.7 mM Glucose + 60 mM KCl (Figure 5G, Video S1, S2). Results were variable since we found that, in one of the differentiations, there was a significant 40% increase in the percentage of KCL-responsive β-cells (GCaMP6^+^ cells) per SC-islet (from 67.6 ± 9.3 to 95.2 ± 1.7) (Figure 5H) in the N21 differentiated SC-islets. However, in a second differentiation, we could detect no difference (69.2 ± 6.8 to 71.7 ± 5.1) and, in a third differentiation, there was only a small, not statistically significant increase (from 63.0 ± 4.3 to 77.3 ± 4.7) (Figure 5H).

Thus, using two of the hES cell reporter lines that we developed, we established that the presence of the N21 supplement in S7 increased significantly the % of monohormonal β-cells. This approach will be widely applicable to test simultaneously several supplements, or their combinations and for different time windows, aiming for the generation of mature SC-islets and with the correct composition of endocrine cells. It also allows the simultaneous assessment of several individual SC-islets per condition, bringing statistical rigor to the approach, while minimizing the necessary number of cells to be analyzed.

## DISCUSSION

The significant advances in the differentiation of hPS cells into pancreatic endocrine cells were based on a detailed understanding of the developmental mechanisms underlying the induction of definitive endoderm, its regionalization and the subsequent induction and differentiation of pancreas progenitors into endocrine progenitors ^62,66,67^. However, the final differentiation steps leading from endocrine progenitors to mono-hormonal pancreatic endocrine cells as well as the subsequent maturation steps that result in full endocrine cell functionality remain to be fully understood. This is inevitably reflected in the remaining shortcomings of the hPS cell differentiation procedure into SC-islets such as the persistence of features of alternative lineages, incomplete conversion into endocrine cells, incomplete maturation, particularly of β-cells, as well as variability of the differentiation among experiments and across different cell lines. Ongoing work on the developmental mechanisms, implicated in endocrine lineage specification and endocrine cell maturation, indicated that additional players, including signaling pathways, mechanical forces and metabolism, affect these processes ^22–30^.

These and additional findings, which will undoubtedly be uncovered in the near future, could help resolve the remaining shortcomings of the differentiation procedure. Ideally, the final SC-islets would contain only pancreatic endocrine cells at appropriate ratios that would have reached full maturation. Endocrine cells in islets act as a community to regulate blood glucose levels and it is now appreciated that cross-talk among different cell types is essential for proper function ^68–71^. To achieve this level of sophistication, additional modifications in the steps leading to PEP cells and then from PEP cells to SC-islets will be needed. The latter steps last several days, usually two weeks in procedures that utilize suspension culture and differentiation in large volumes ^7,9,11,13^ and more than three weeks in procedures that employ smaller culture volumes ^8,10,15^. It is conceivable that additional modifications would optimally be applied only during specific time windows. For example, maturation signals would be needed towards the end of the procedure whereas signals that would contribute to enhancing endocrine specification and eliminating other lineages would be necessary at the early stages of the process. All these possibilities add up to a substantial number of alternative procedures that could only be assessed longitudinally using live imaging and multiple appropriate reporters. In anticipation of this need, we have generated reporter lines that allow longitudinal live cell imaging and demonstrated that they can be used to generate SC-islets containing functional β-cells in arrayed microwells. It is important to note that all reporters have been incorporated in the corresponding gene loci ensuring that they are under the control of all corresponding regulatory elements. Additionally, we have taken all possible precautions, such as the use of a self-cleaving peptide and the removal of the selection cassette, to minimize interference to the targeted locus and we analyzed heterozygote lines to minimize the effects of the targeting on the expression levels of corresponding proteins.

The *INS ^eGFP^/ GCG ^mCHERRY^* line can be employed to assess the degree of endocrine specification as well as the early steps in the maturation process with the resolution of INS^+^ / GCG^+^ cells. Targeting the *SOM* allele with a distinct fluorescent reporter, for example, the blue fluorescent protein (BFP), in the same line, will extend the scope of this analysis to identifying optimal conditions for a balanced ratio of the three endocrine types to ensure optimal function of the derived SC-islets.

An important functional maturation parameter is intracellular Ca^2+^ fluxes and oscillations in β-cells of the SC-islets in response to glucose or other secretagogues. The *INS ^GCaMP6^* line that we developed allows the recording of individual β-cells, the calculation of the percentages of responding β-cells as well as the amplitude and synchronicity of their response. Since live Ca^2+^ combined with glucose stimulation is less amenable to automation, this line will be useful in assessing specific conditions that would be identified by high-content screens.

The functional maturation process continues after the resolution of polyhormonal cells and a suitable reporter line to follow the late steps of SC-islet maturation is still lacking. In humans, postnatal maturation of β–cells extends until the end of puberty ^16,17^ and it is, to a large extent, driven by the gradual upregulation of the transcription factor MAFA ^18^. In humans, reduced *MAFA* expression or missense MAFA mutations have been linked to poor GSIS and T2D ^19–21^. MAFA expression is particularly low in currently hPS cell-derived β cells and substantial upregulation of this gene, along with other genes marking advanced maturation, was only seen six months after engraftment ^72^. Very little is known regarding the factors that promote this final maturation step and the generation of the *INS ^eGFP^/ MAFA ^mCHERRY^* reporter which will allow the assessment of several possibilities. The full concordance of the mCHERRY expression with MAFA immunofluorescence on cryosections suggests that the line will be useful in identifying such conditions.

An additional application that would likely be very insightful is the transplantation of SC-islets, derived from these lines, to follow their maturation in vivo. The reporters will enable the sorting of the cells and their analysis at any point after transplantation. Moreover, the transplantation of these SC-islets in a natural body window, such as the anterior chamber of the eye, will enable the longitudinal in vivo investigation of the transplanted cells with sufficient cellular resolution ^73,74,75^. This approach would provide high-quality data on the engraftment and maturation of SC-islets after their transplantation and allow the comparison of different in vitro differentiation protocols as well as the impact of different systemic environments on cell maturation and function.

We have adapted the differentiation procedure to generate SC-islets in 24-well wells, each containing a few hundred microwells, each carrying a single SC-islet. This offers the opportunity to assess several conditions in parallel – one condition applied per well – with a relatively small number of cells. In this application, we used 1.5 million PP cells per well but this number can be further reduced for smaller microwells. An additional, important advantage of the approach is that high-content live imaging of multiple single SC-islets provides statistically rigorous results and an accurate estimate of the variability among SC-islets differentiated under the same conditions. This is an important consideration because uniformity of SC-islets is expected to provide more reliable clinical outcomes. Live imaging also offers the possibility of longitudinal assessment of different conditions to identify optimal time windows.

Here we demonstrated the feasibility of the approach and found that supplementing the S7 medium with the N21 supplement significantly increases the number of monohormonal β-cells by 50% and this was reflected in the nearly two-fold increase in insulin content. Despite this increase and no changes in insulin secretion, expression of *PDX1* and *NKX6-1* did not increase suggesting that PDX1 and NKX6-1 protein stability has increased following N21 treatment. Interestingly, the supplement does not affect the numbers of bi-hormonal or single α-cells and thus the net effect is a corresponding increase in the number of endocrine cells in the SC-islets to the detriment of non-endocrine cell types. The persistence of bi-hormonal cells suggests that the conversion process of progenitor cells, which may become other cell types, to endocrine progenitors, to bi-hormonal cells and, finally, to β-cells, remained incomplete and this could be due to either or both of two possibilities. Etither here was not enough time, in which case S7 should be included in the S6 medium as well, or some N21 components might be present in lower concentrations than necessary. In the latter case, different N21 formulations will need to be tested and the approach described here would be ideal for this task. N21 is a complex supplement but it is serum-free and composed of twenty-one defined substances that can be produced under GMP conditions. Thus, it is suitable for the generation of SC-islets for clinical applications but, to simplify its application, it will be of interest to define exactly which of the components are essential and which are dispensable.

The approach described here could be used to identify conditions that have a similar effect on α- and δ-cells, so that mature SC-islets, containing exclusively mature endocrine cells in the appropriate ratios, can be generated efficiently and cost-effectively.

### Data availability

The sequences of the targeted alleles, including the 5’ and 3’ homology arms, have been submitted to GenBank with accession numbers OR729124 (*INS ^eGFP^*), OR727484 (*GCG ^mCHERRY^*), OR727485 (*MAFA^mCHERRY^*) and OR727486 (*INS ^GCaMP6^*).

## Author contributions

AG conceptualized the study, EZ, LJ, MB, DP, MAR-T, ER-A, VK, AKM-A, CC and KN performed experiments, AG, EZ, LJ, MB, CC, SS and KN analysed the data, AG wrote the manuscript with input and editing from EZ, LJ, MB and KN. AG supervised the study and acquired funding.

## Acknowledgments

Research in the AG laboratory was supported by grants from the German Center for Diabetes Research (DZD) (grant 82DZD00101) and the German Research Foundation (DFG) (grants GA-2004/3-1 and IRTG 2251). This study was also supported by a Seed CRTD Grant to IG, DP and CC.

---

**Figure S1.**
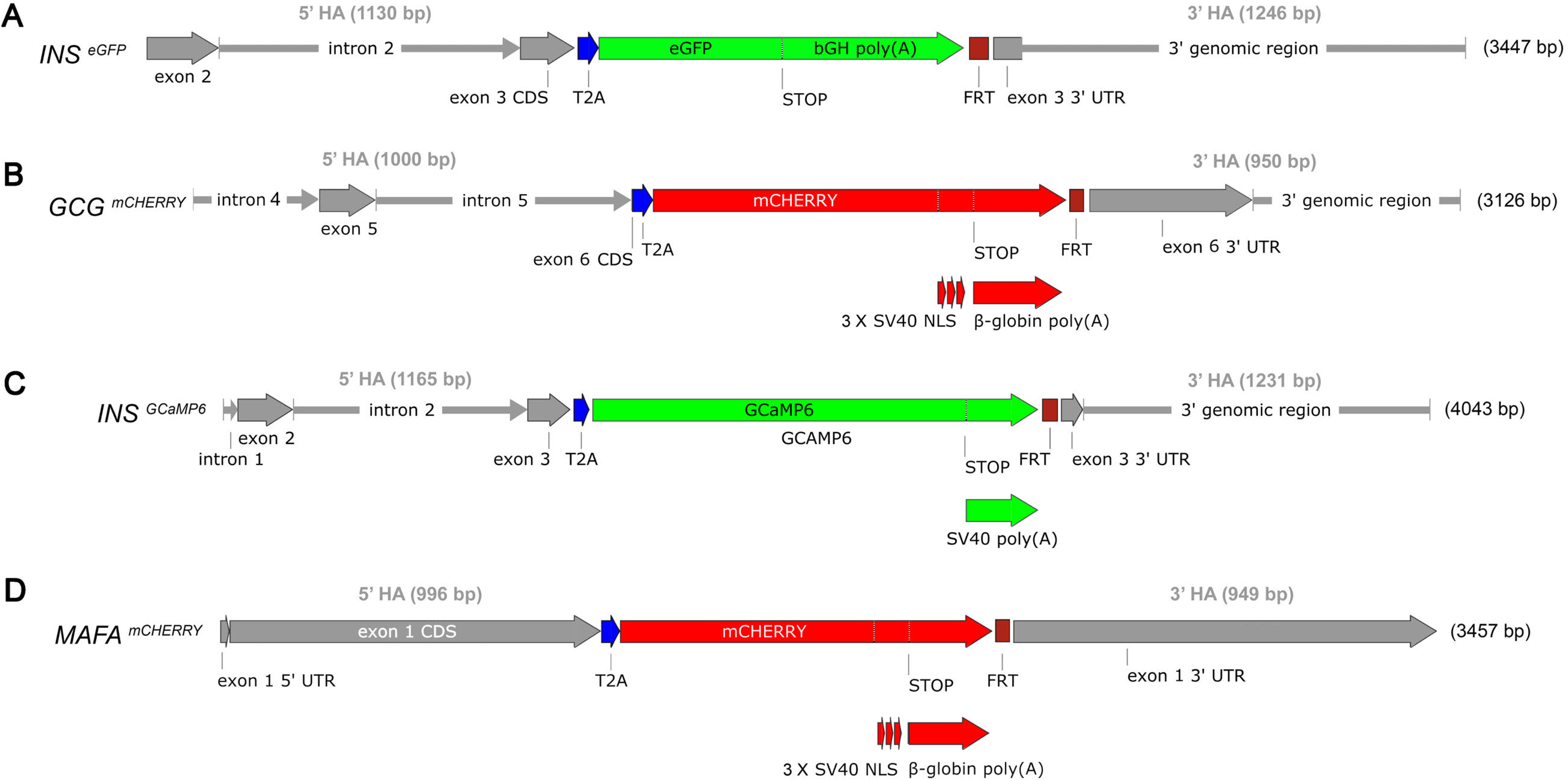
Detailed maps of the targeted alleles. (A-D) Representation of the loci of the genes of interest after the targeting and the removal of the selection cassette. 5’ and 3’ homology arms (HA) containing introns and coding exons, or 3’ UTRs and downstream genomic regions, respectively, are shown in grey. T2A coding sequences (blue) are inserted in-frame with the amino acid coding part of the last exon of the corresponding gene. The reporter is inserted in-frame with the T2A and is followed by a polyA (red or green). mCherry CDS includes a nuclear localization signal (NLS) at its 3’. The single FRT site left after the Flp recombinase-mediated removal of the selection cassette is shown in purple.

**Figure S2.**
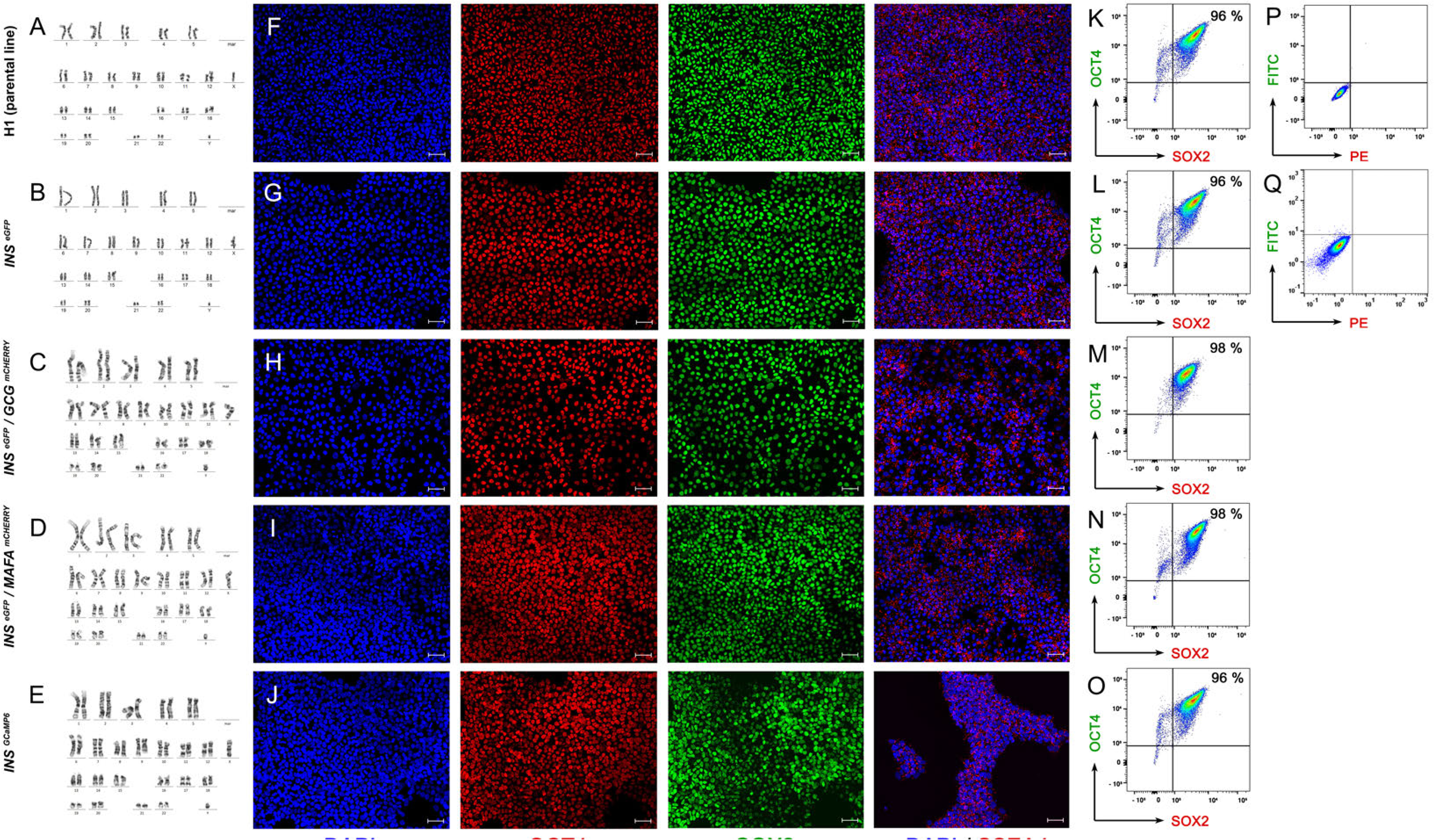
Expression of pluripotency and definitive endoderm markers in H1 cells, derived reporter lines and differentiated cells into definitive endoderm. (A-E) Karyotyping by G-banding for the parental H1 line (A) and in the derived reporter lines H1 *INS^eGFP^* (B), H1 *INS ^eGFP^*/*GCG ^mCherry^* (C), H1 *INS ^eGFP^*/*MAFA ^mCherry^* (D) and H1 *INS ^GCaMP6^* (E). (F-J) Fluorescent immunostainings show strong and broad expression of the pluripotent markers OCT4, SOX2 and SSEA4 in the parental H1 (F) and in the derived reporter lines H1 *INS ^eGFP^* (G), H1 *INS ^eGFP^*/*GCG ^mCherry^* (H), H1 *INS ^eGFP^*/*MAFA ^mCherry^* (I) and H1 *INS ^GCaMP6^* (J). (K-P) Flow Cytometry analyses of H1 (parental line) (K), H1 *INS ^eGFP^* (L), H1 *INS ^eGFP^*/*GCG ^mCherry^* (M), H1 *INS ^eGFP^*/*MAFA ^mCherry^* (N) and H1 *INS ^GCaMP6^* (O) hES cells using antibodies for OCT4 and SOX2. (P) Shows a representative flow cytometry control sample stained only with the secondary antibodies used for the detection of OCT4 and SOX2. (Q) Representative flow cytometry control sample of DE cells stained only with the secondary antibodies used for the detection of SOX17 and FOXA2. Scale bar corresponds to 50 µm.

**Figure S3.**
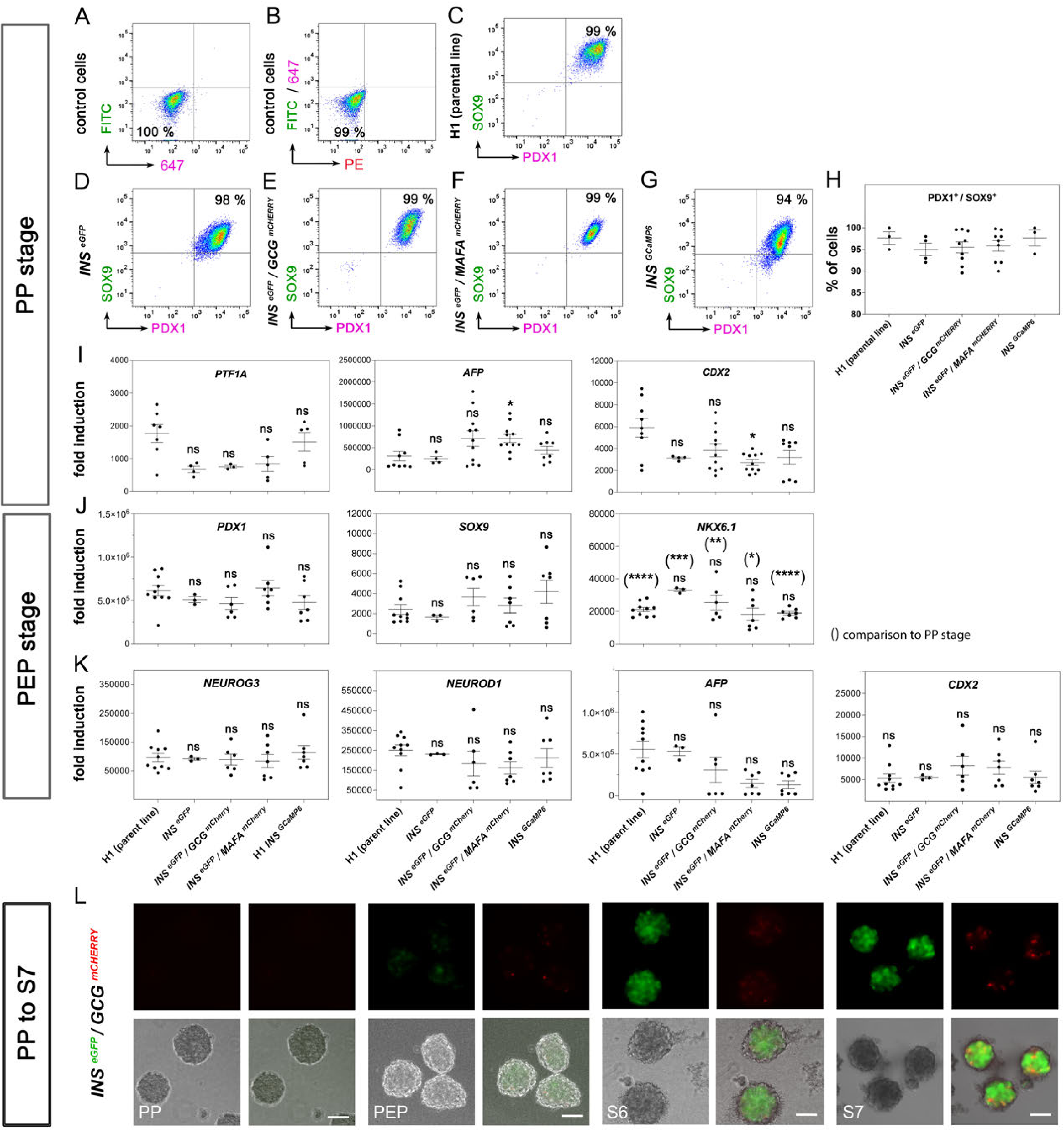
Differentiation efficiency into pancreatic progenitors and pancreatic endocrine progenitors. (A, B) Flow Cytometry analyses of PP cells stained only with the secondary antibodies providing the gating for PDX1^+^/NKX6.1^+^ cells (A) and the gating for NKX6.1^+^ cells (B). (C-G) Selected flow cytometry analyses showing the % of double PDX1^+^/SOX9^+^ cells in H1 (C), H1 *INS ^eGFP^* (D), H1 *INS ^eGFP^*/*GCG ^mCherry^* (E), H1 *INS ^eGFP^*/*MAFA ^mCherry^* (F) and H1 *INS ^GCaMP6^* (G) derived PP cells. (H) Flow cytometry analyses of PDX1^+^ / SOX9^+^ PP cells derived from the H1 (parental line), H1 *INS ^eGFP^*, H1 *INS ^eGFP^*/*GCG ^mCherry^*, H1 *INS ^eGFP^*/*MAFA ^mCherry^* and H1 *INS ^GCaMP6^* hES cells. Dots represent values from independent experiments and horizontal lines represent the mean ± standard error of the mean (SEM). (I) Gene expression analyses by RT-qPCR of PP cells, derived from H1 as well as the reporter hPS cell lines, showing the expression of the pancreatic progenitor marker *PTF1A*, the liver marker *AFP* and the gut *CDX2* marker expression. Gene expression levels are shown as fold change relative to those in undifferentiated H1. Dots represent values from independent experiments and horizontal lines represent the mean ± SEM. (J, K) Gene expression analyses by RT-qPCR of PEP cells, derived from H1 as well as the reporter hPS cell lines, showing the expression of *PDX1*, *SOX9* and *NKX6.1* genes (K), expression of the key endocrine progenitor genes *NEUROG3* and *NEUROD1* in all the lines as well as expression of the liver marker *AFP* and the intestinal marker *CDX2* (L). Gene expression levels are shown as fold change relative to those in undifferentiated H1. Dots represent values from independent experiments and horizontal lines represent the mean ± SEM. (L) Time course of the *INS ^eGFP^* and *GCG ^mCherry^* reporter expression by whole mount live immunofluorescence one day after clustering of the PP cells (PP) and at the end PEP, S6 and S7 stages. Statistical analyses were performed using the non-parametric Kruskal-Wallis test, using the H1 (parent line) as the basis for the comparison. For the comparison of the *NKX6.1* expression between PEP and PP cells of the same line the Welsh’s t-test was used. *p* ≤ 0.05 (*), *p* ≤ 0.005 (**), *p* ≤ 0.0005 (***) and *p* ≤ 0.0001 (****). Scale bars correspond to 100 μm

**Figure S4.**
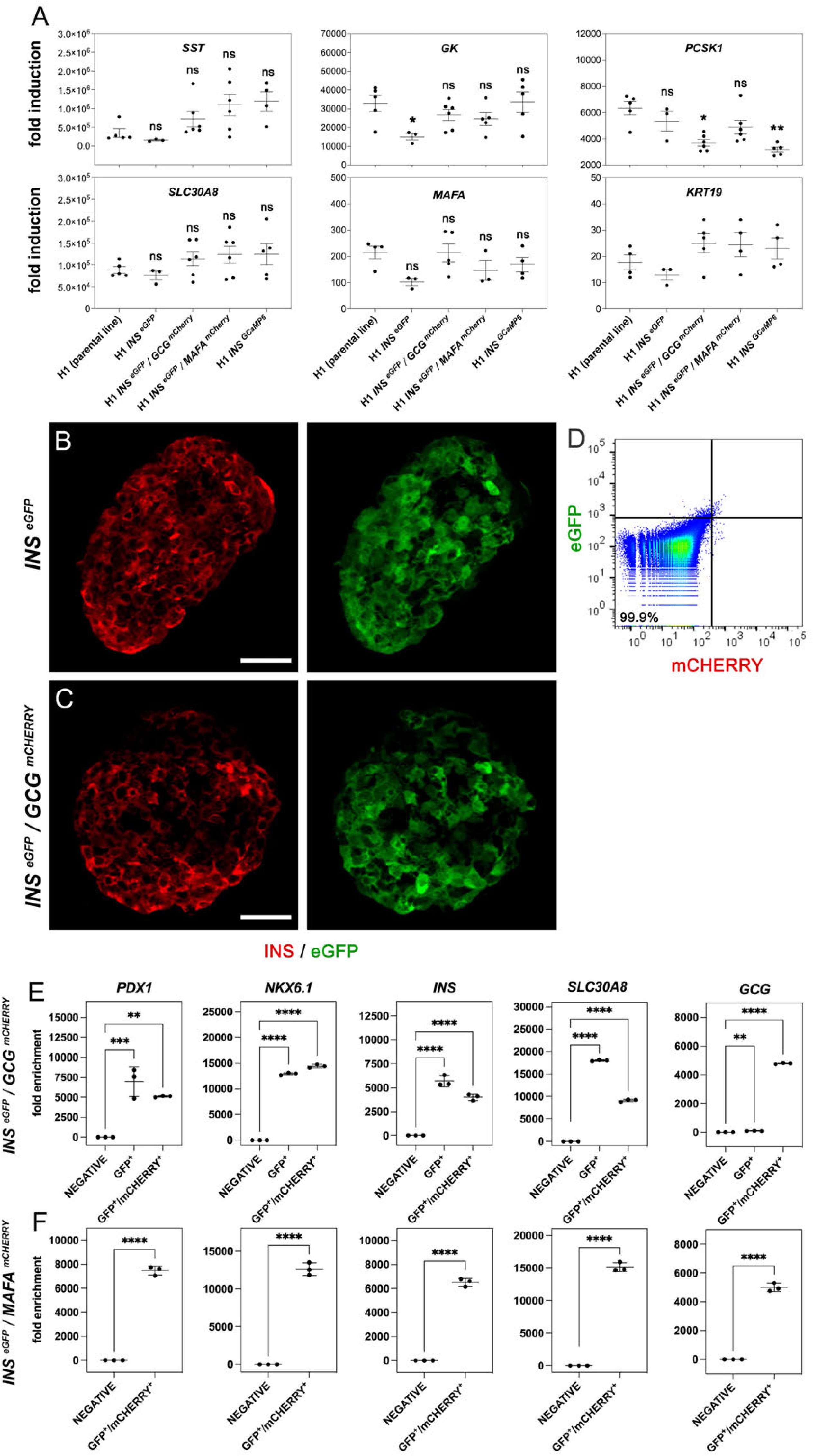
Differentiation efficiency into SC-islets and reporter functionality. (A) Gene expression analyses by RT-qPCR of SC-islets for the hormones *GCG* and *SST*, genes involved in the β-cell maturation and functionality, *PAX4, ZnT8, PC1/3, MAFA* as well as for the hepatic gene marker *AFP,* the gut marker *CDX2* and the duct marker *CK19*. Data are expressed as fold change relative to undifferentiated H1. Dots represent values from independent experiments and horizontal lines represent the mean ± SEM. (B) Representative immunofluorescent staining of *INS ^eGFP^* SC-islet cryosections for INS and eGFP. (C) Representative immunofluorescent staining of *INS ^eGFP^*/*GCG ^mCherry^* SC-islet cryosections for INS and eGFP. (D) Representative flow cytometry analyses of live H1 parental line SC-islet cells used to set the gating for the analyses of the reporter lines. (E) Fold-enrichment of *PDX1*, *NKX6.1*, *INS*, *GCG* and *ZnT8* expression in GFP^+^ and GFP^+^/mCHERRY^+^ FACS isolated cells from *INS ^eGFP^*/*GCG ^mCherry^* SC-islets. Expression in control cells was set as 1. Dots represent values from independent experiments and horizontal lines represent the mean ± SEM. (F) Fold-enrichment of *PDX1*, *NKX6.1*, *INS*, *GCG* and *ZnT8* expression in GFP^+^ FACS isolated cells from *INS ^eGFP^*/*MAFA ^mCherry^* SC-islets. Expression in control cells was set at 1. Dots represent values from independent experiments and horizontal lines represent the mean ± SEM. Statistical analyses were performed using the non-parametric Kruskal-Wallis test, using the H1 (parental line) as control for the comparison with *p* ≤ 0.05 (*), *p* ≤ 0.005 (**) and *p* ≤ 0.0005 (***). Scale bars correspond to 50 μm.

**Figure S5.**
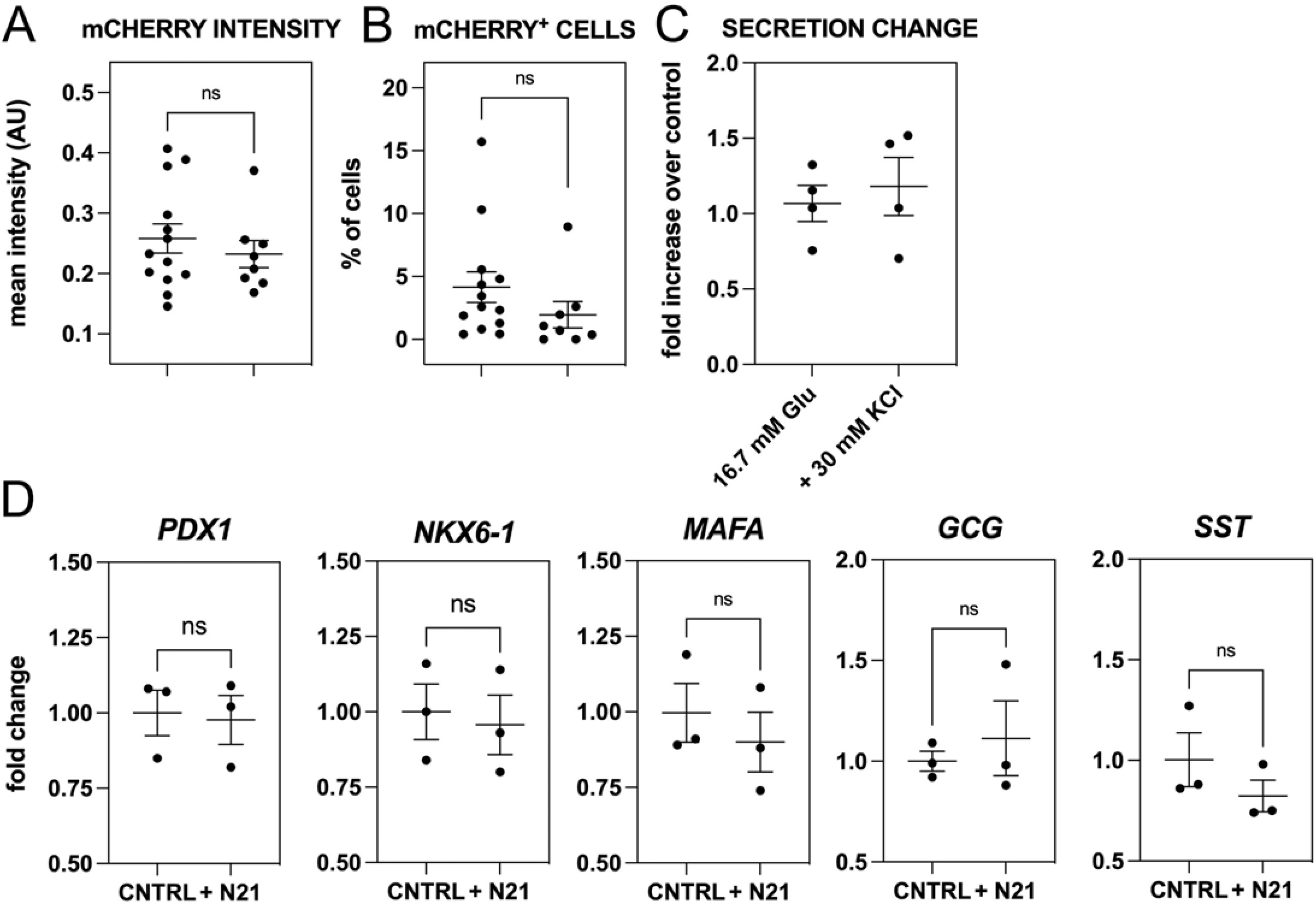
N21 promotes β-cell formation and responsiveness. (A, B) Supplementing S7 with N21 did not change the mean mCHERRY intensity per SC-islet (A) or the % of mCHERRY^+^ cells per islet (B) as compared to control (CNTRL) islets. Dots in A, B correspond to individual SC-islets. (C) Supplementing S7 with N21 did not change the response of SC-islets to either 16.7 mM glucose or to 16.7 mM + 30 mM KCl. Dots represent values from independent differentiation experiments. (D) Changes in the expression of *PDX-1*, *NKX6-1*, *MAFA*, as well as *GCG* and *SOM* upon supplementation of S7 medium with N21 as compared to control conditions (CNTRL). Dots represent values from independent differentiation experiments. Statistical analyses were performed using the non-parametric Kruskal-Wallis test, using the H1 (parental line) as control for the comparison with *p* ≤ 0.05 (*), *p* ≤ 0.005 (**) and *p* ≤ 0.0005 (***). Horizontal lines represent the mean ± SEM. Scale bars correspond to 50 μm.

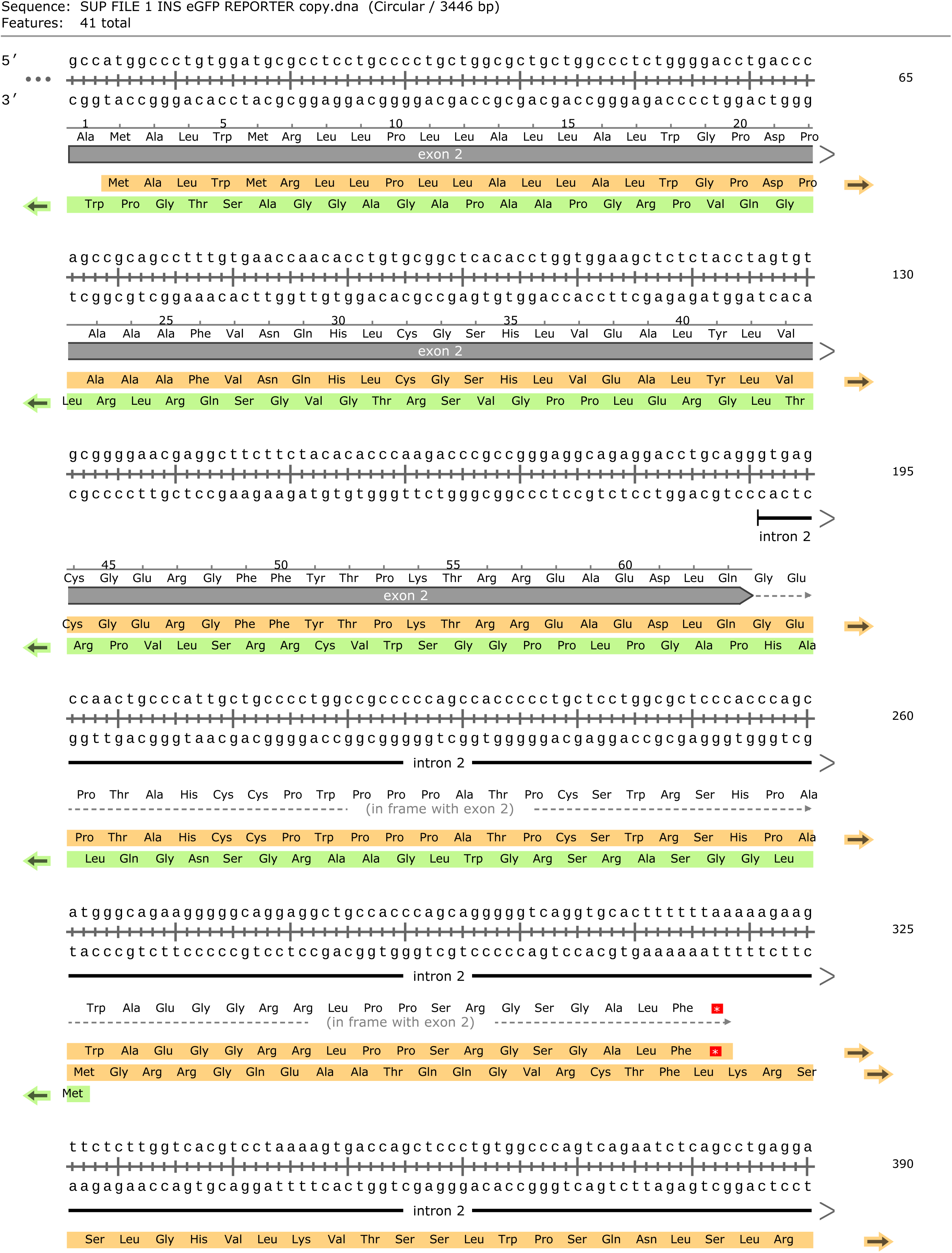
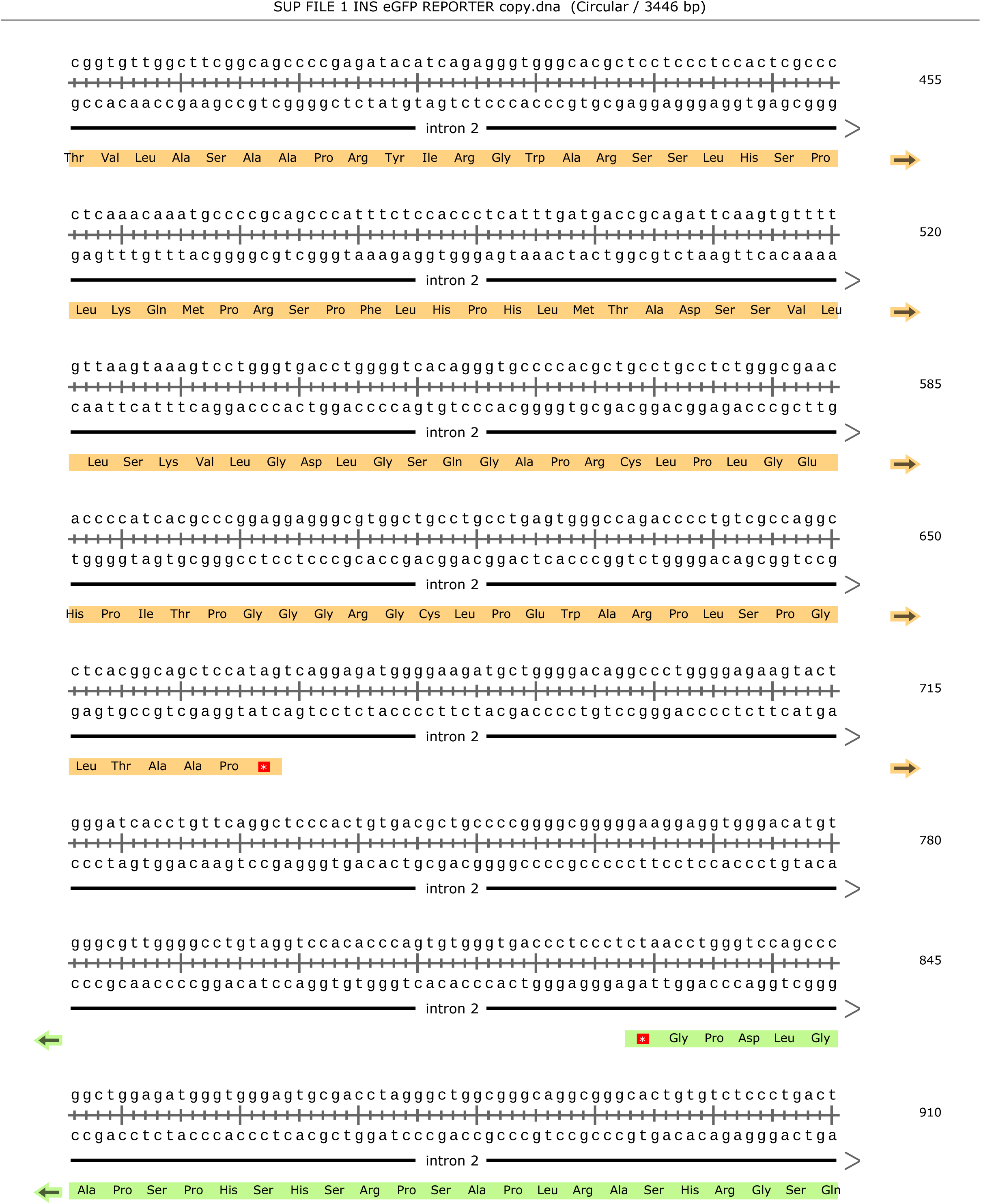
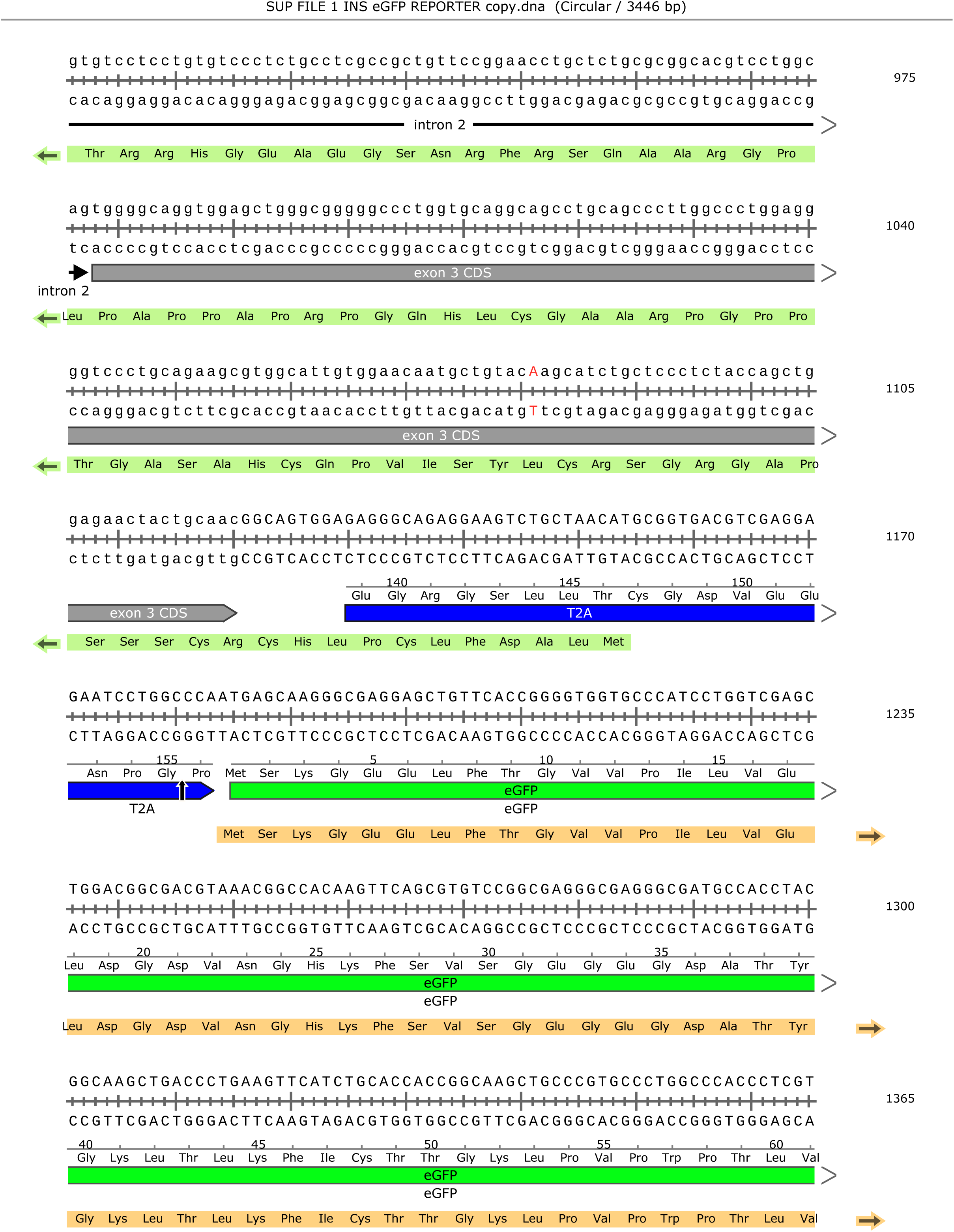
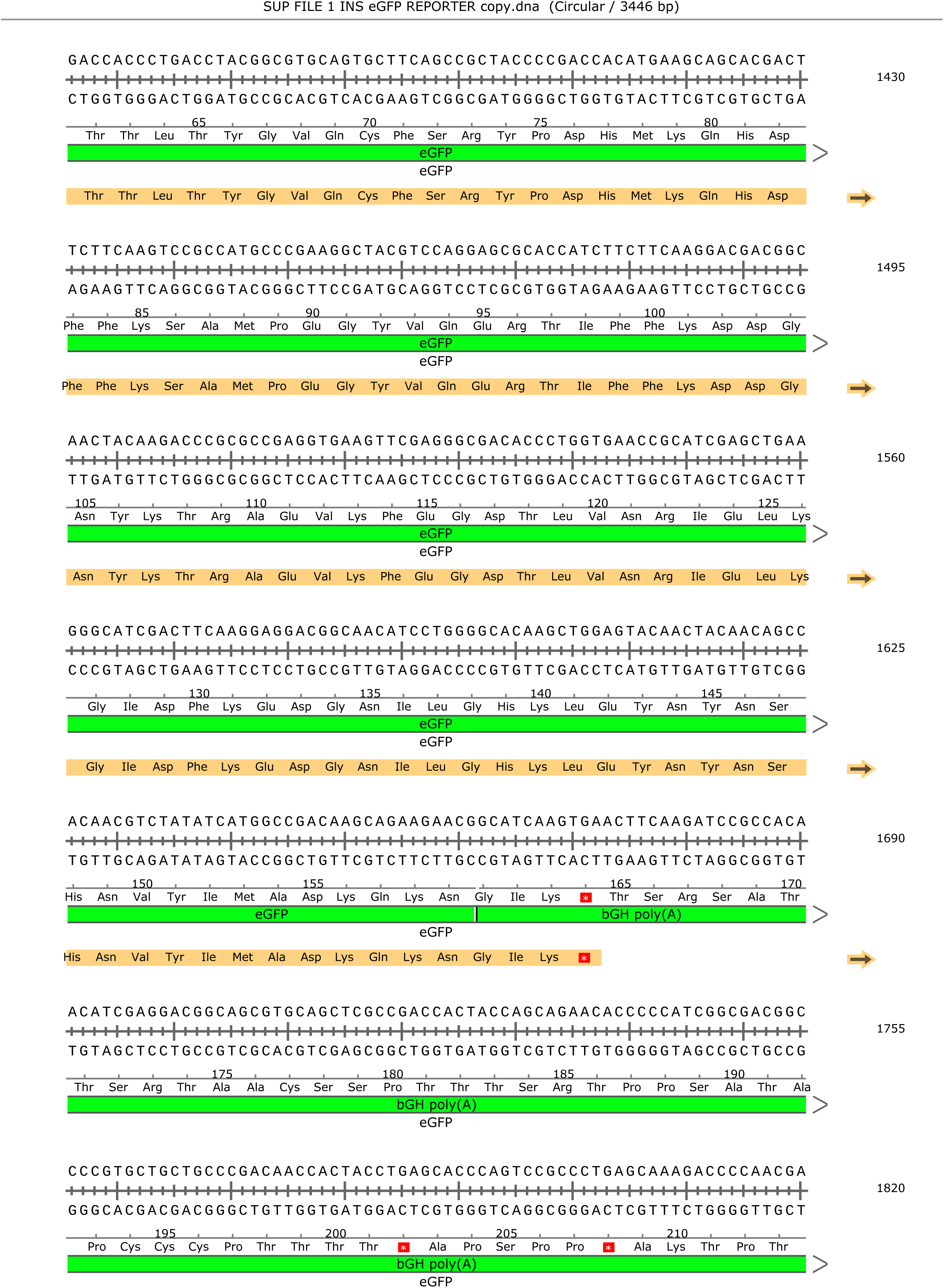
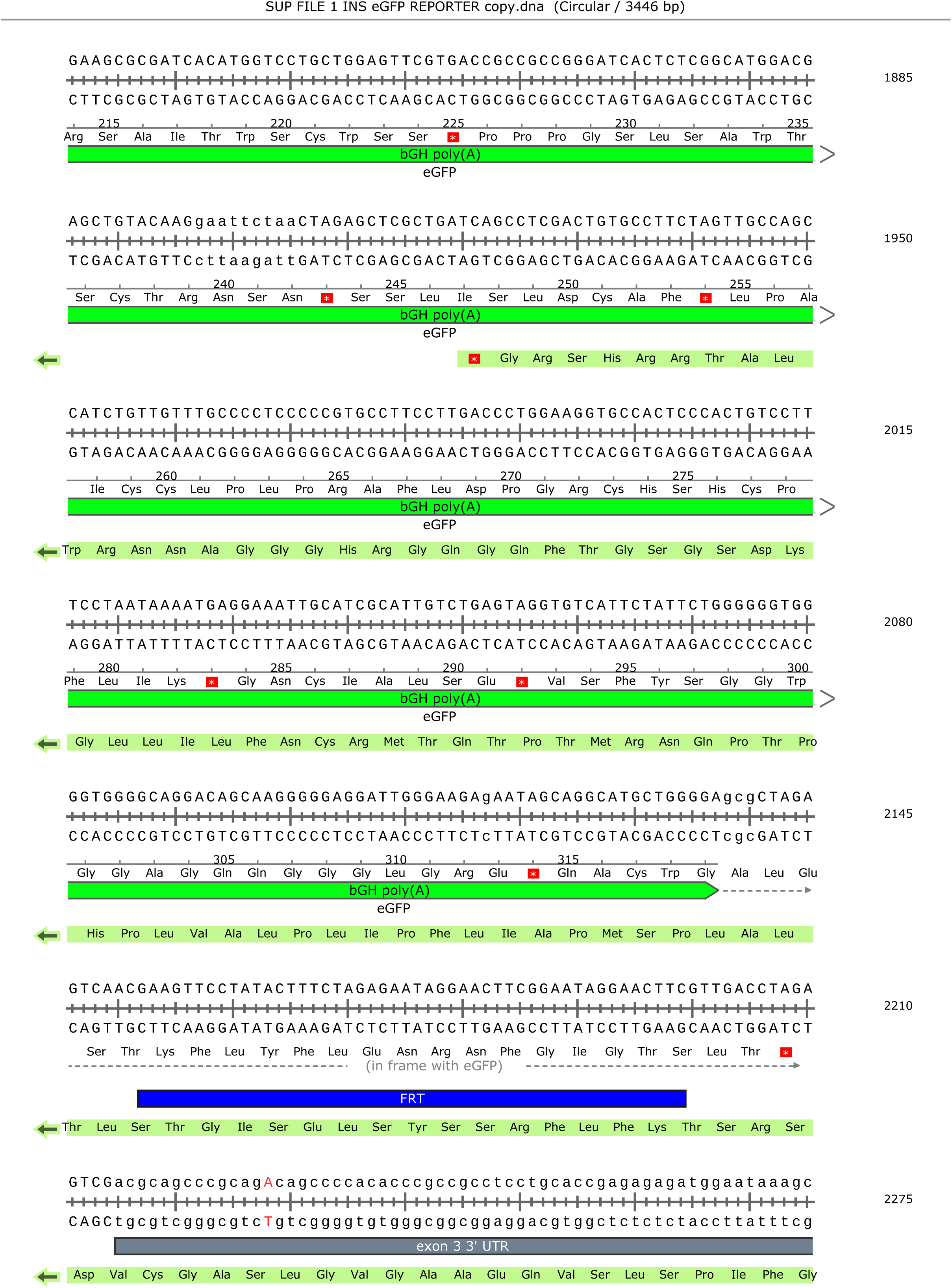
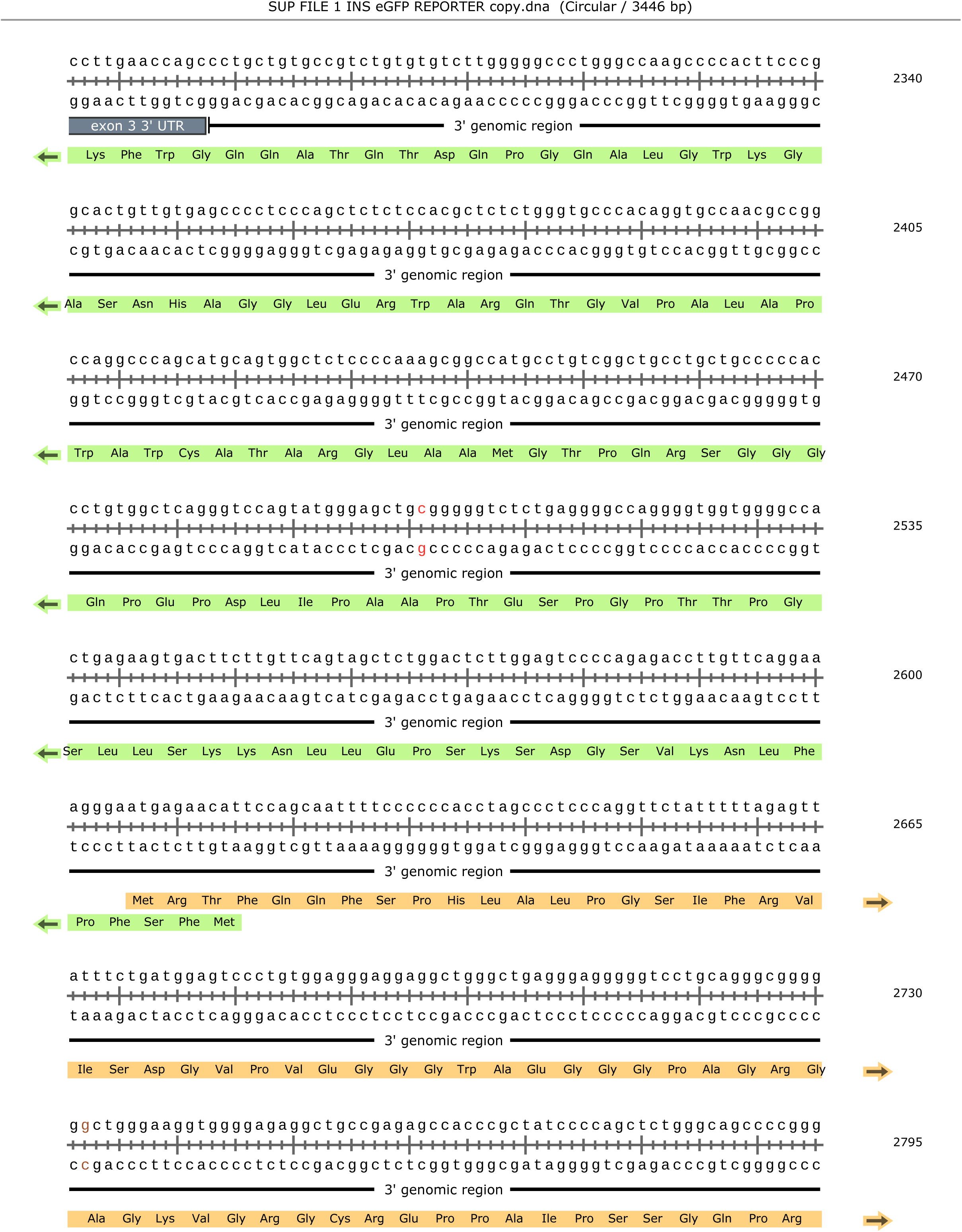
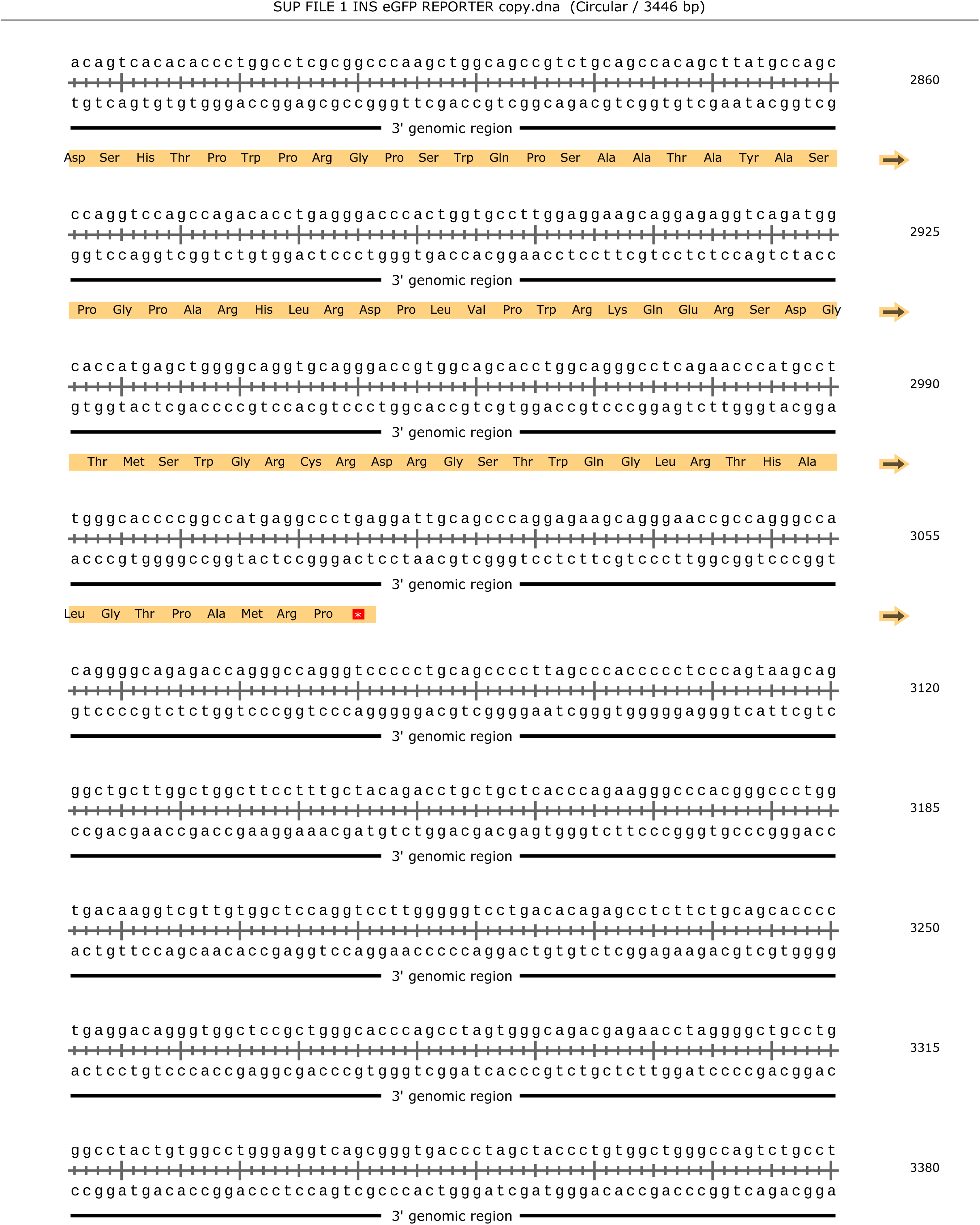
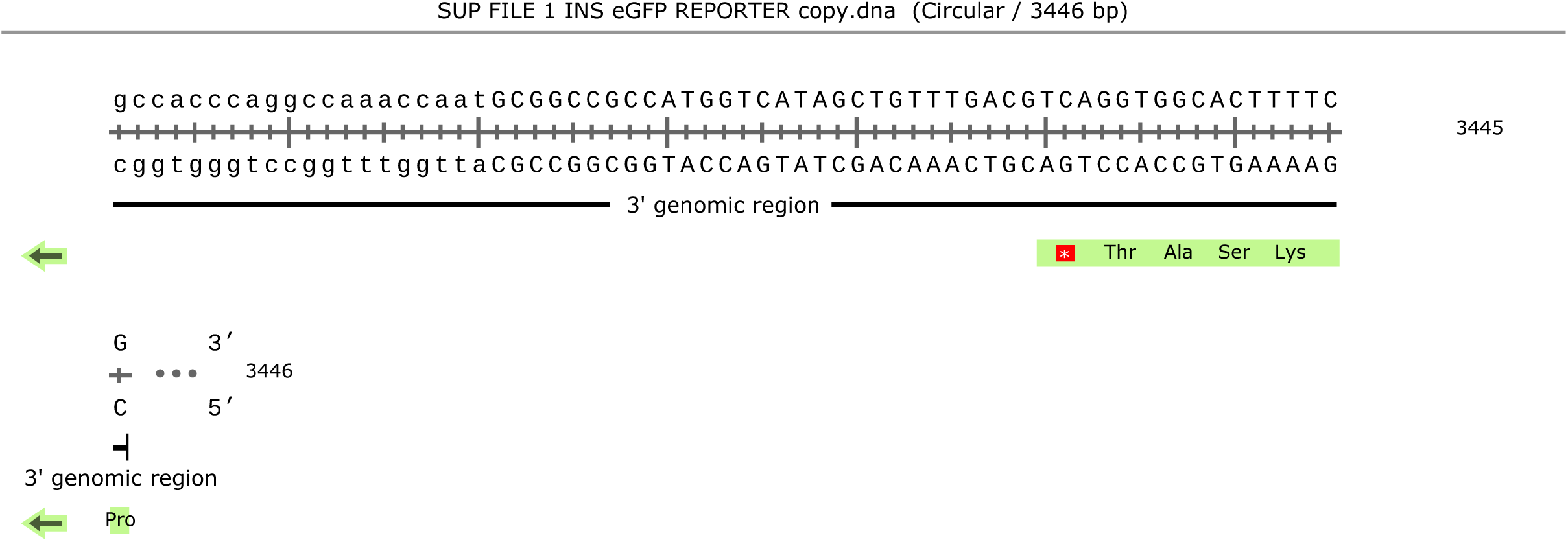

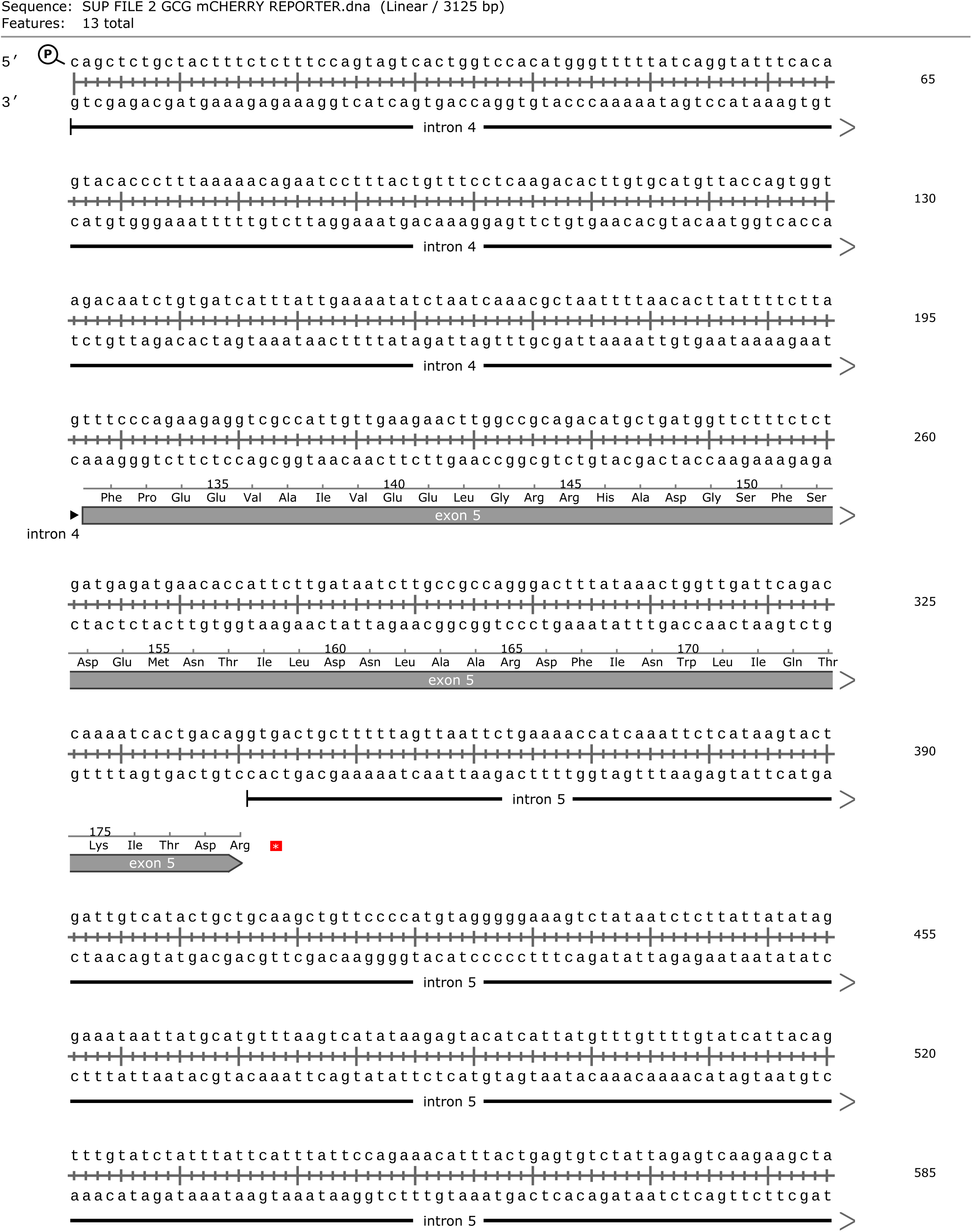
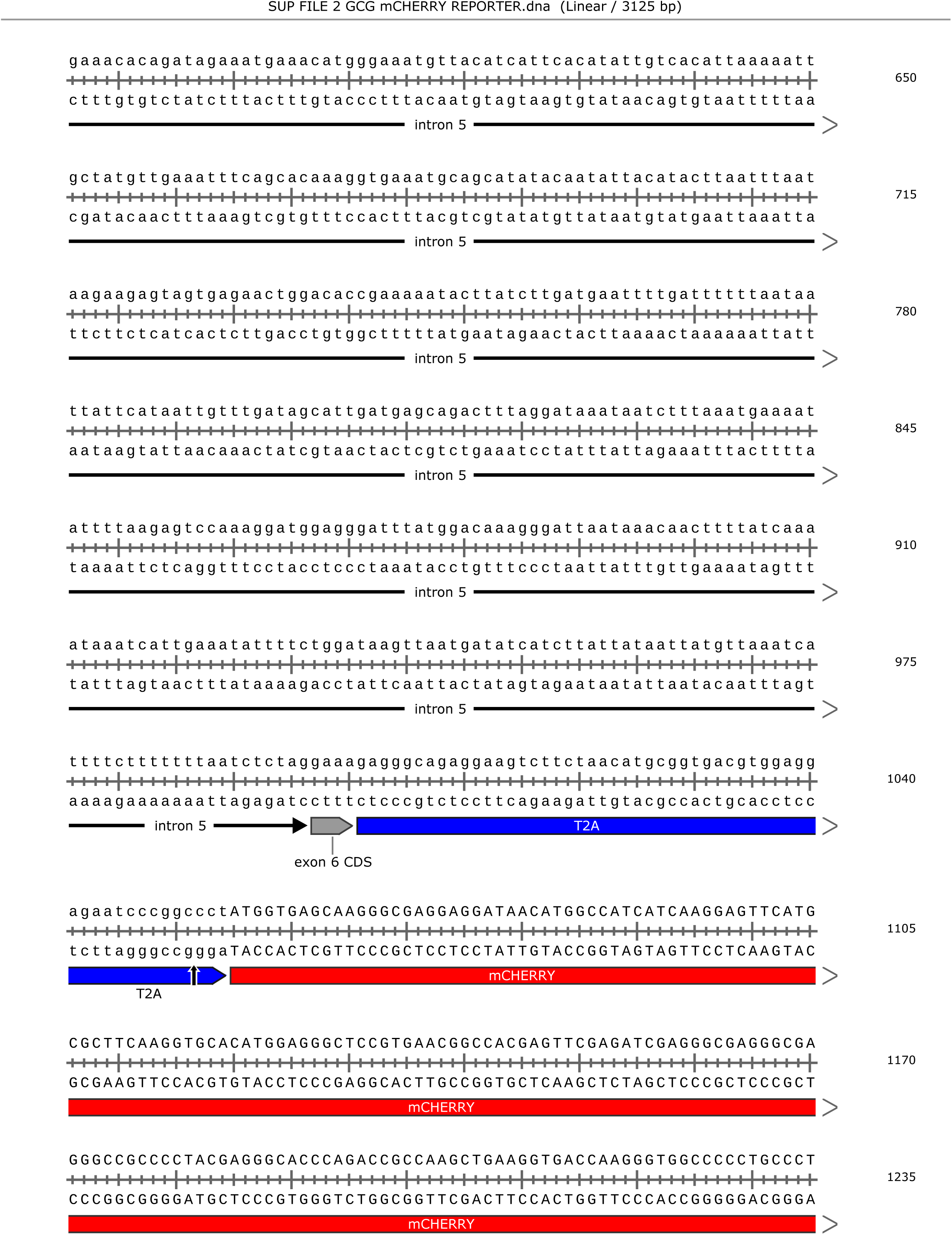
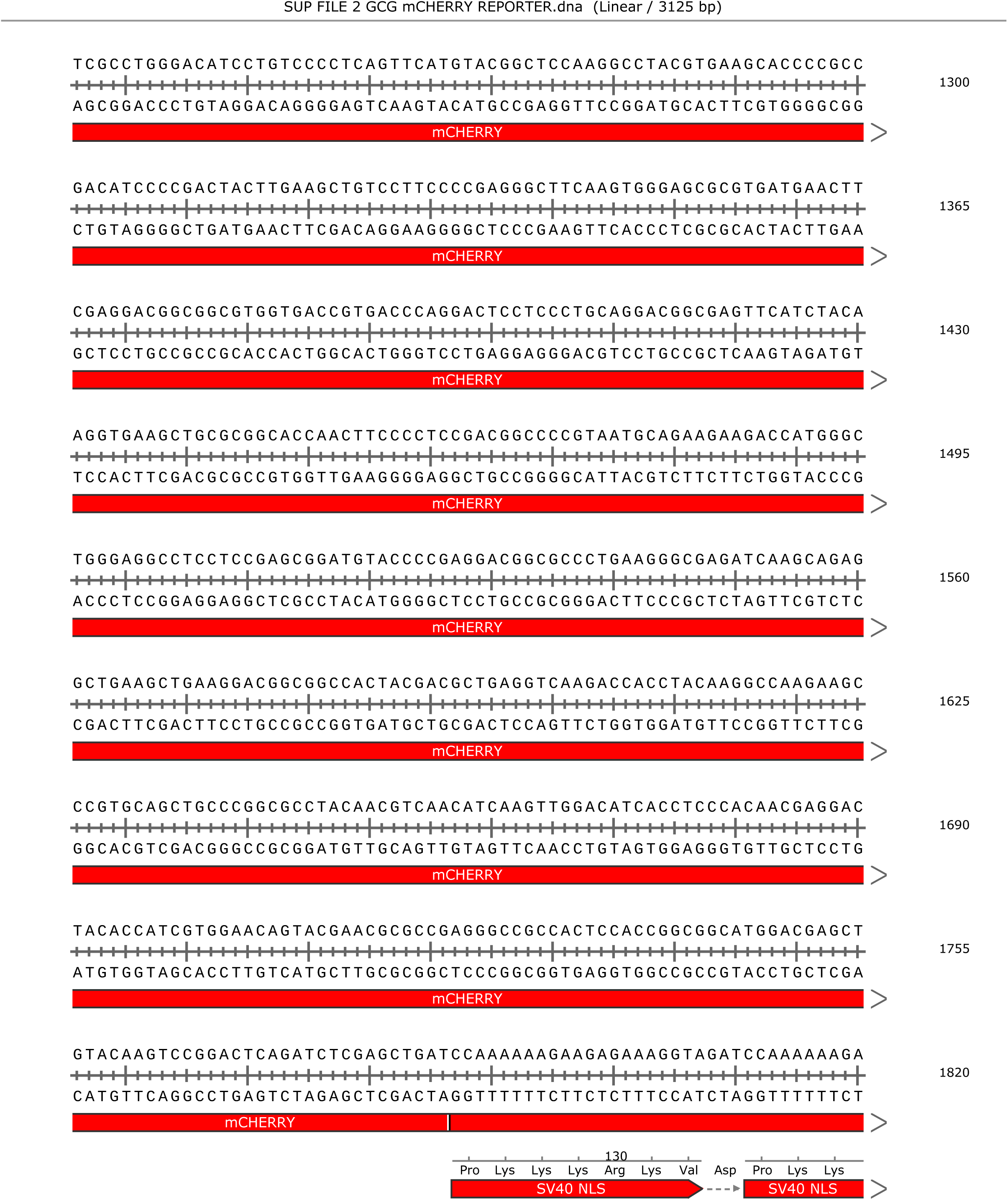
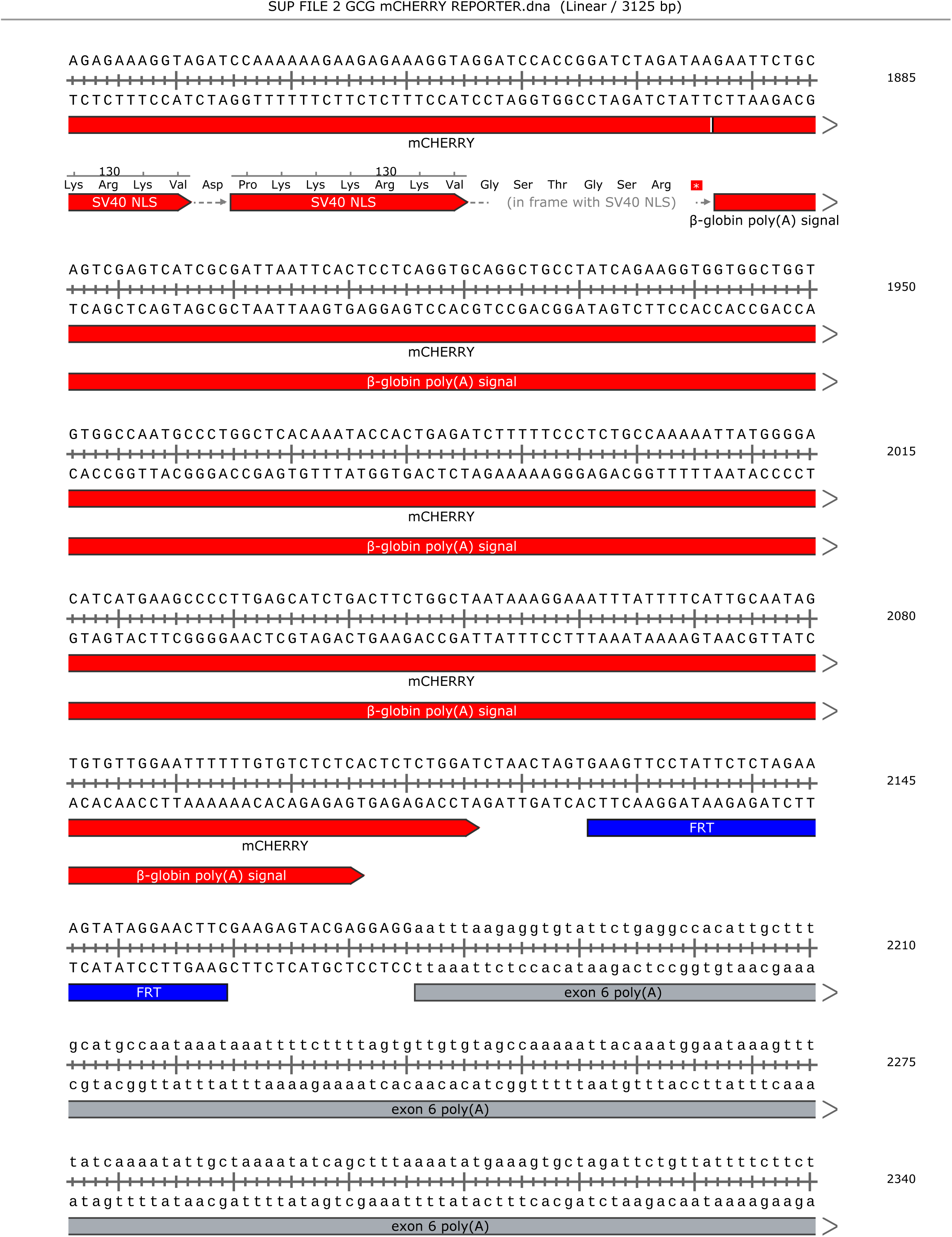
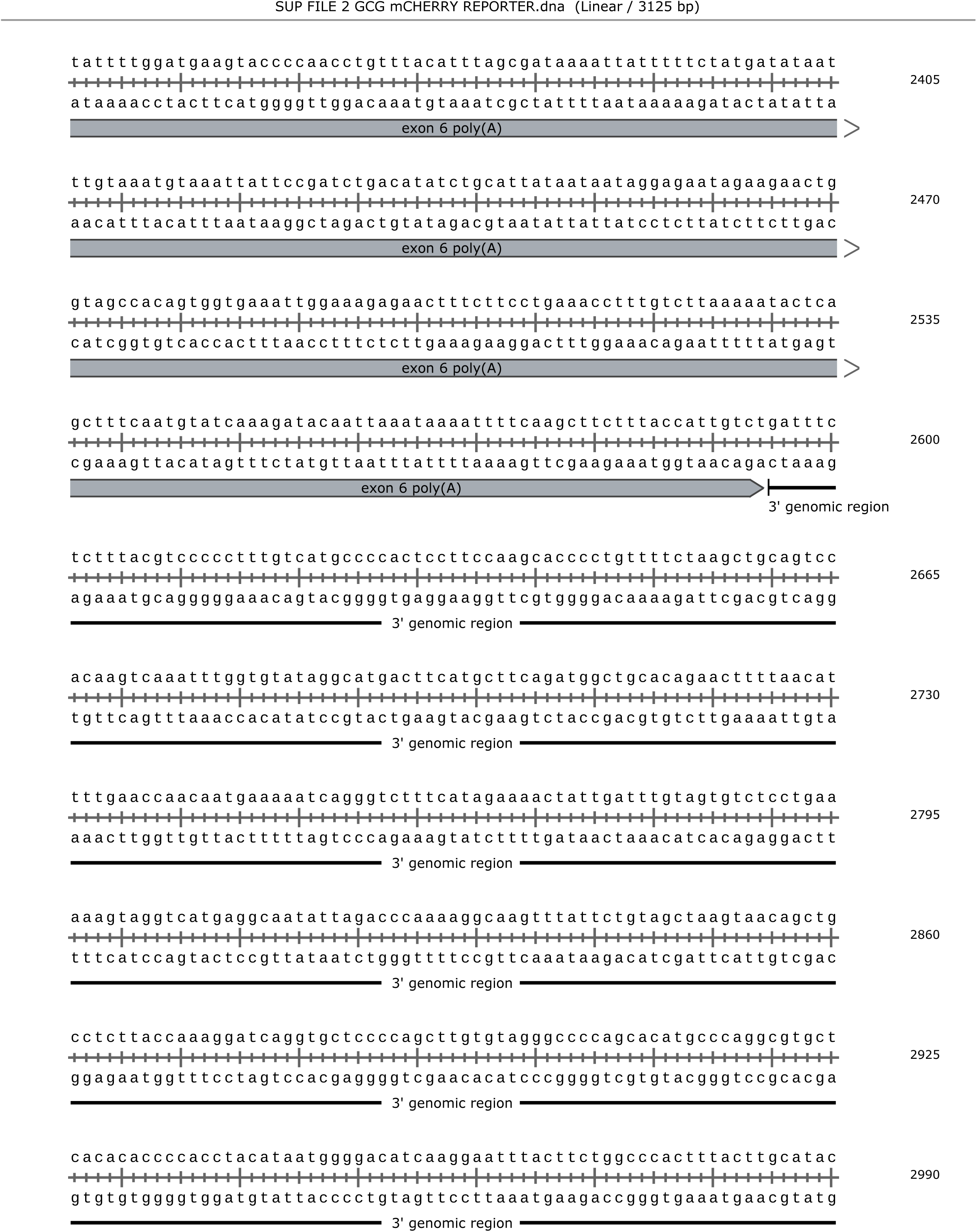
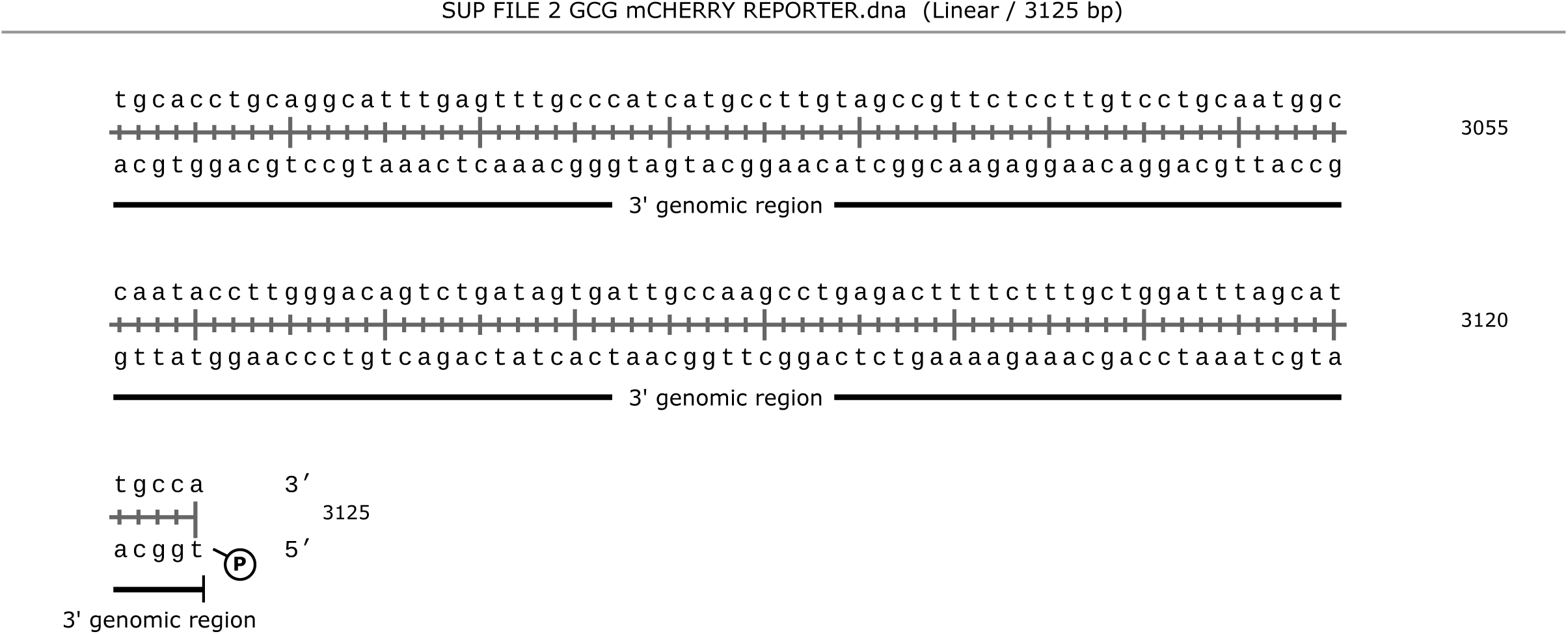
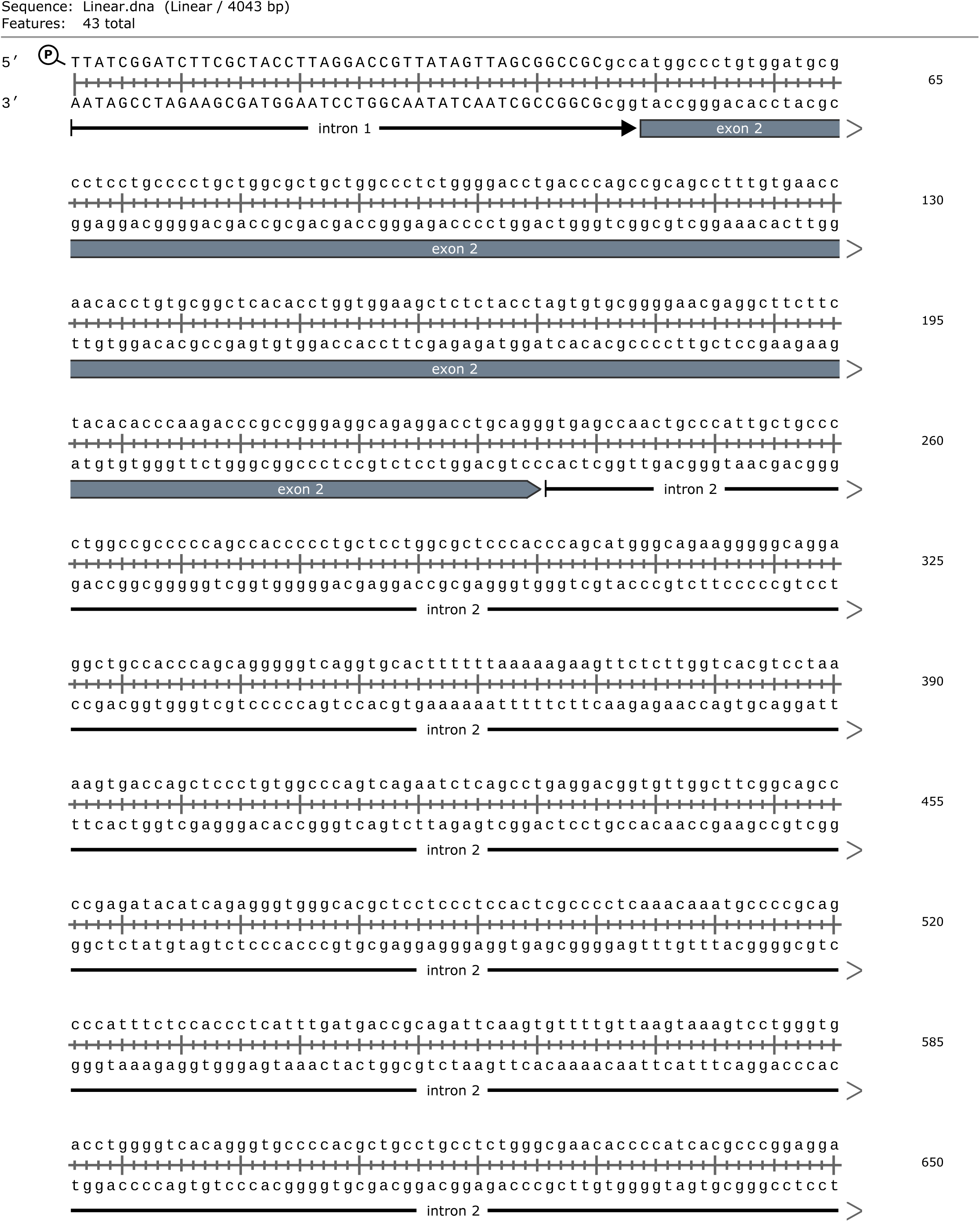
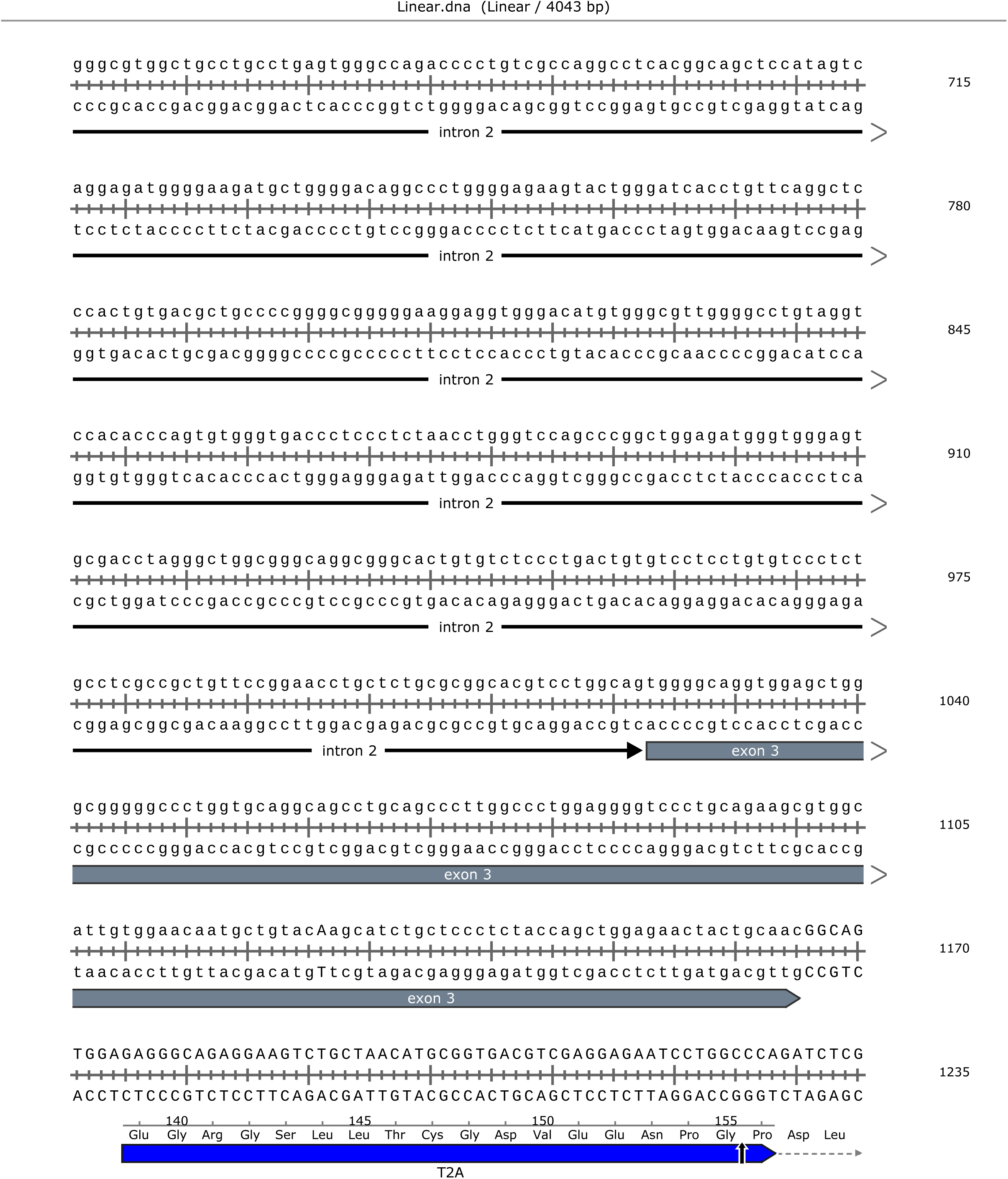
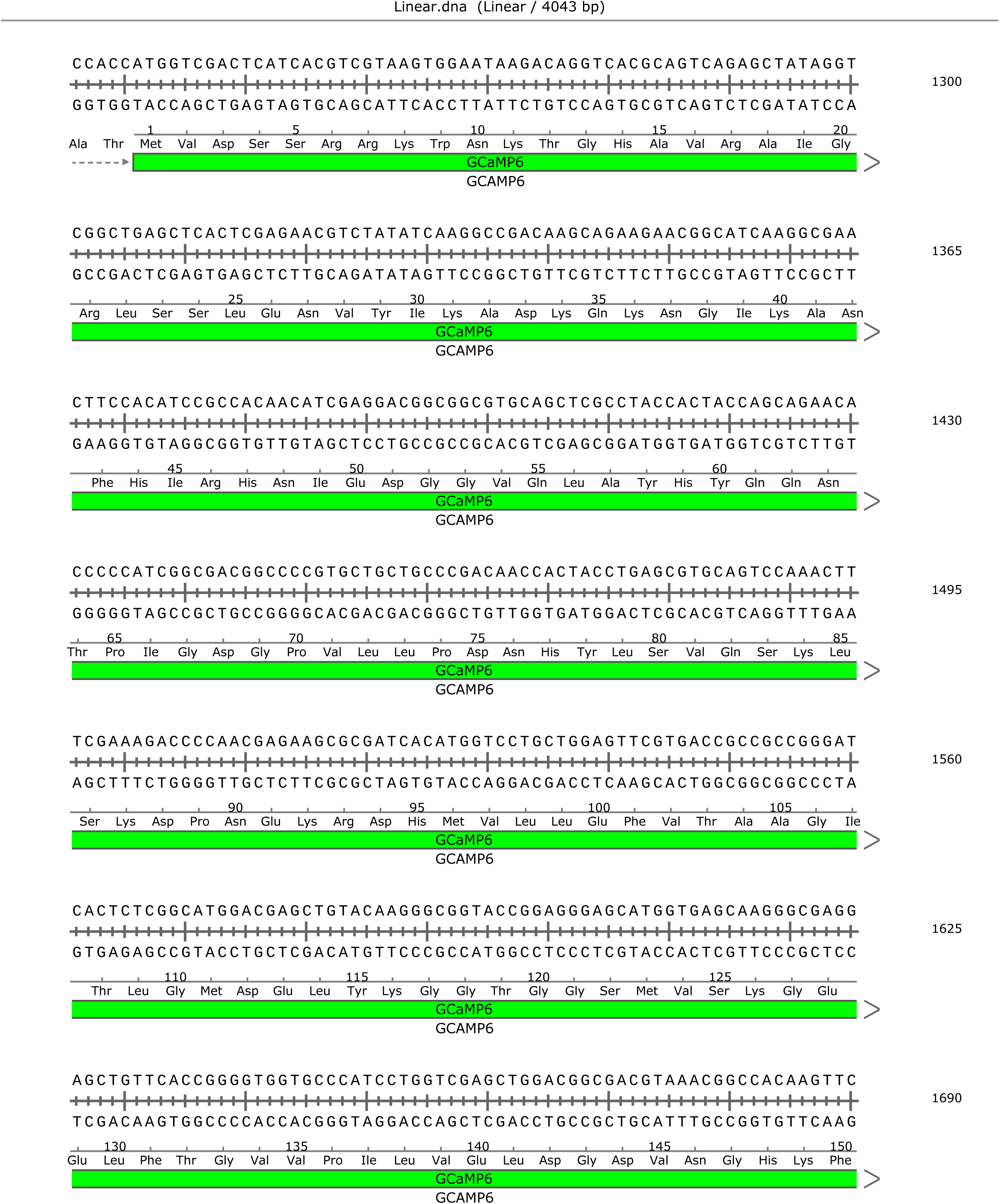
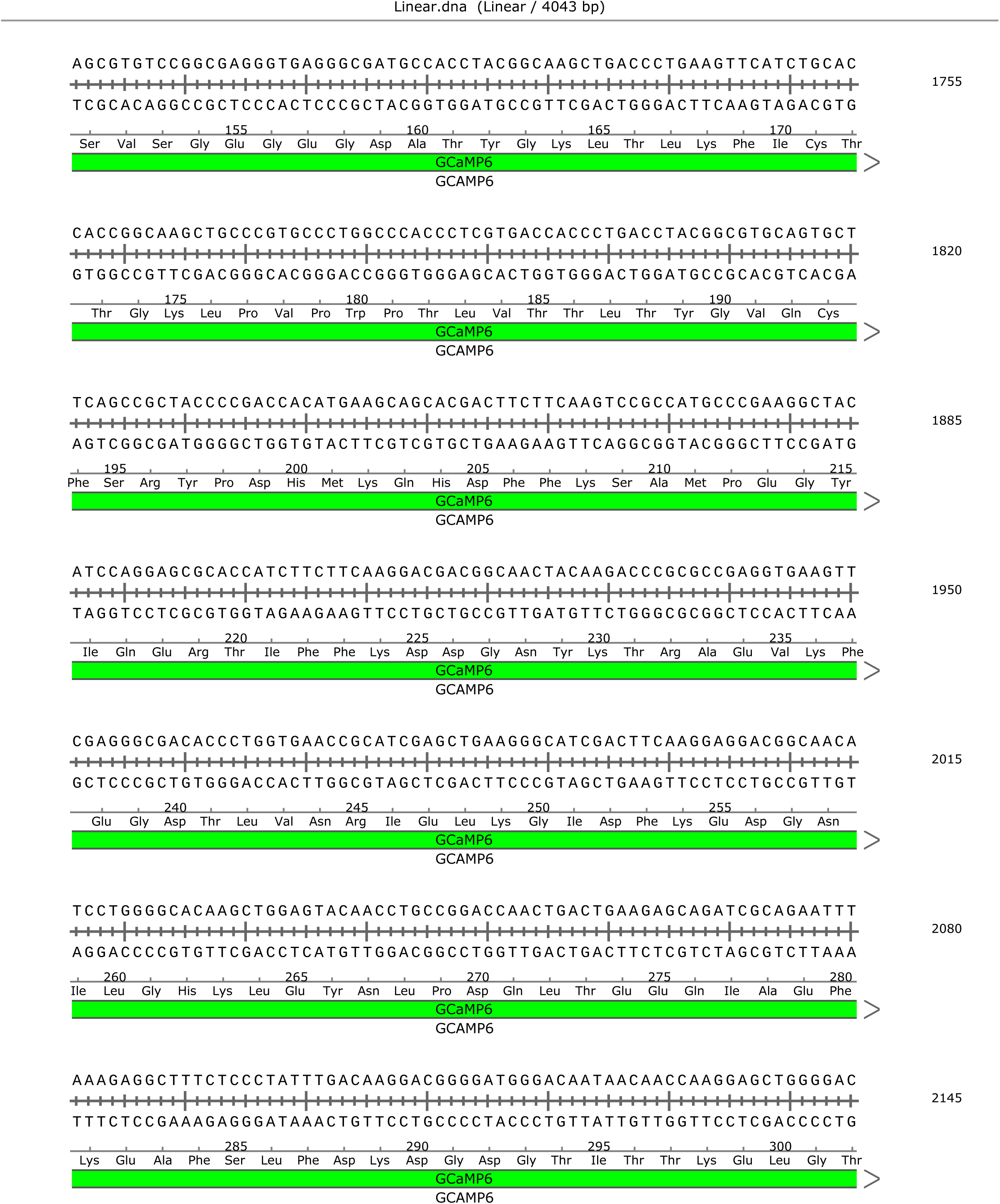
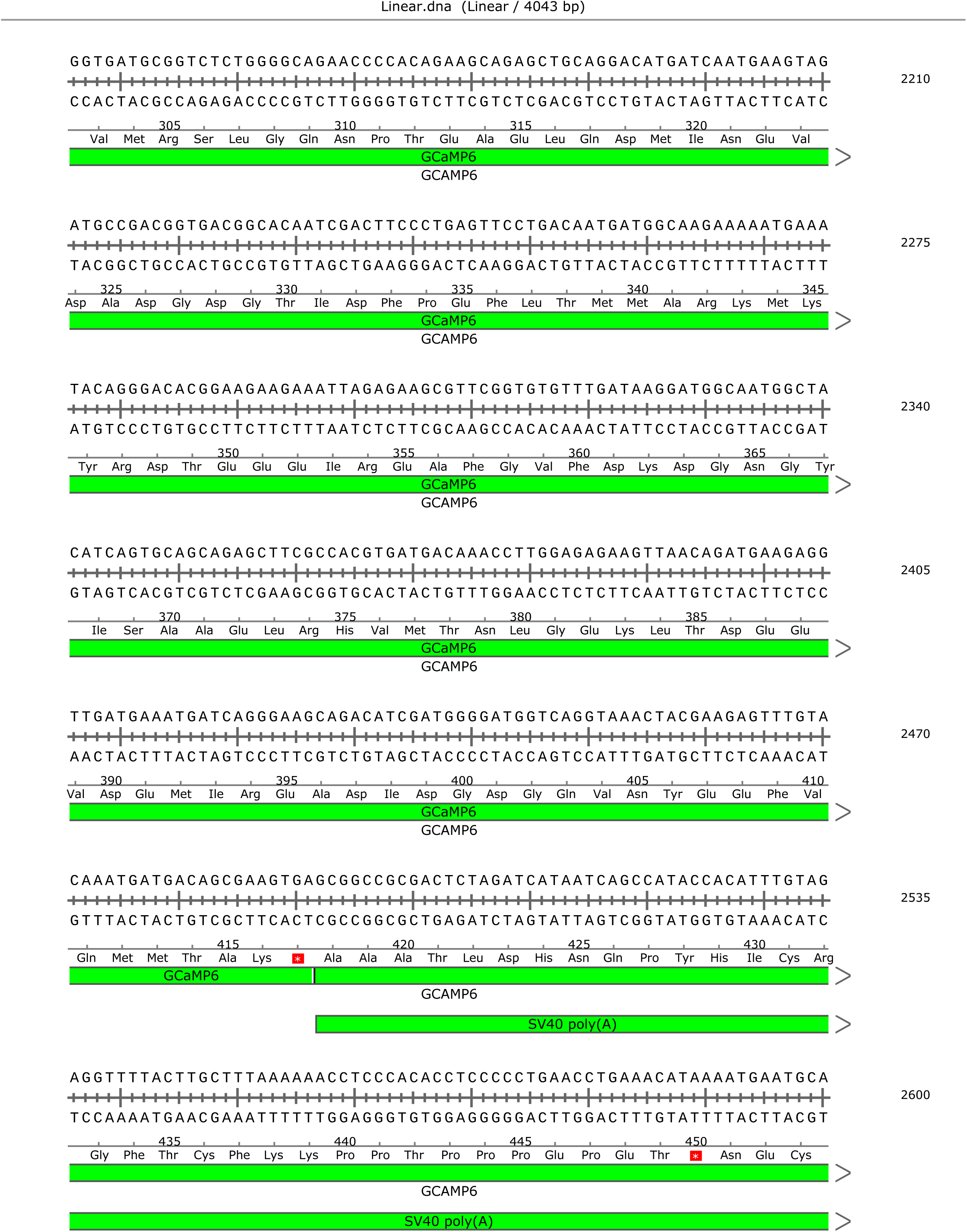
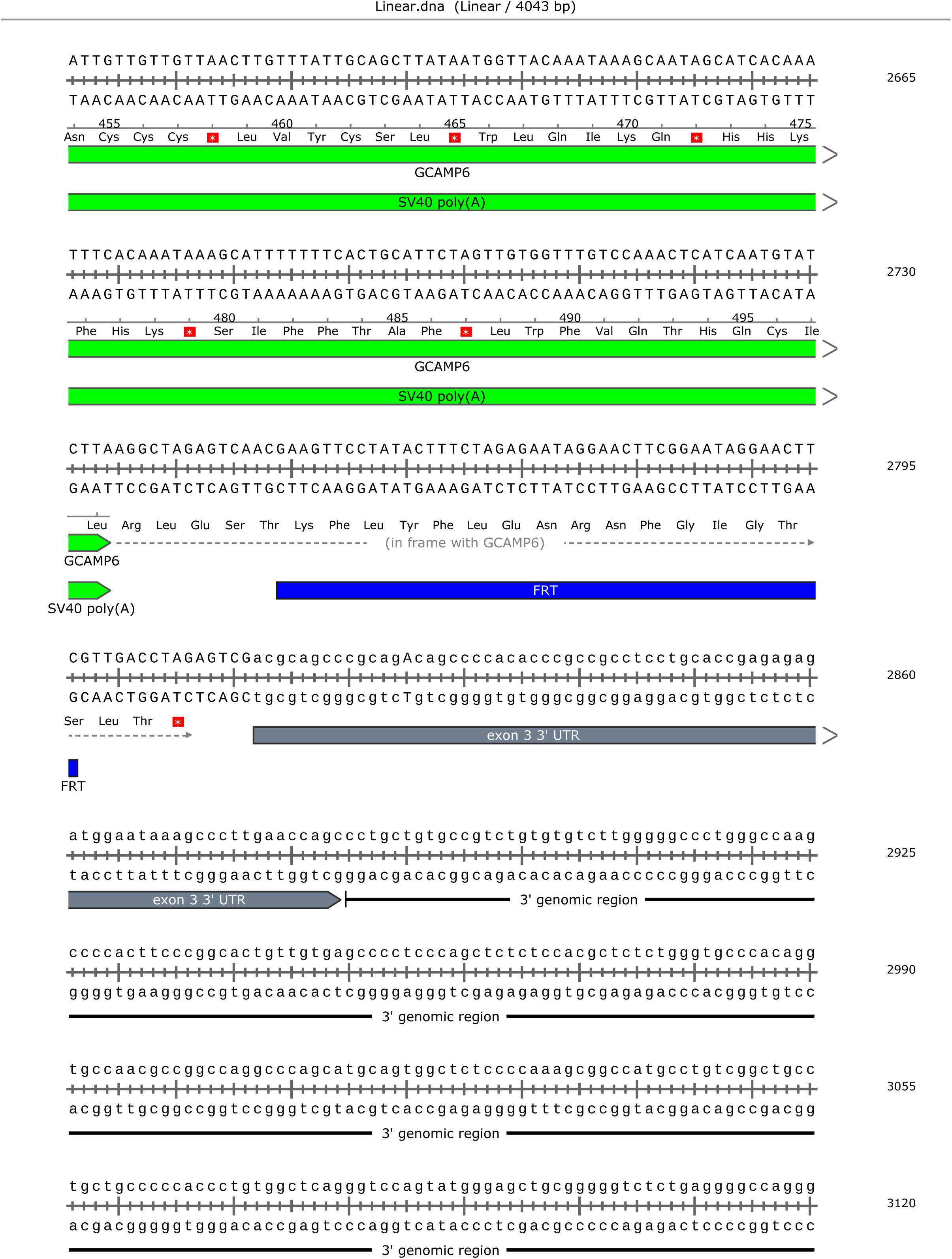
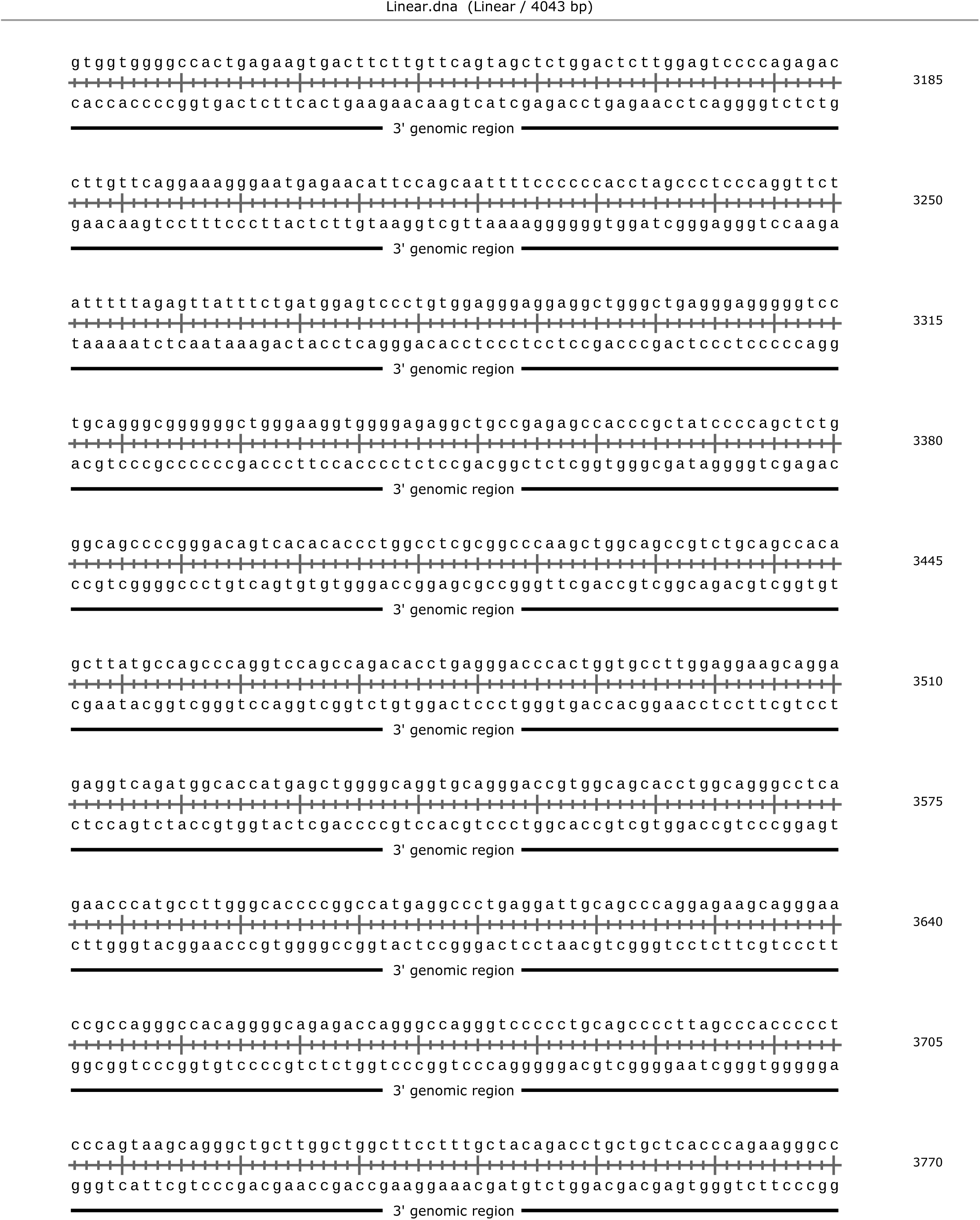
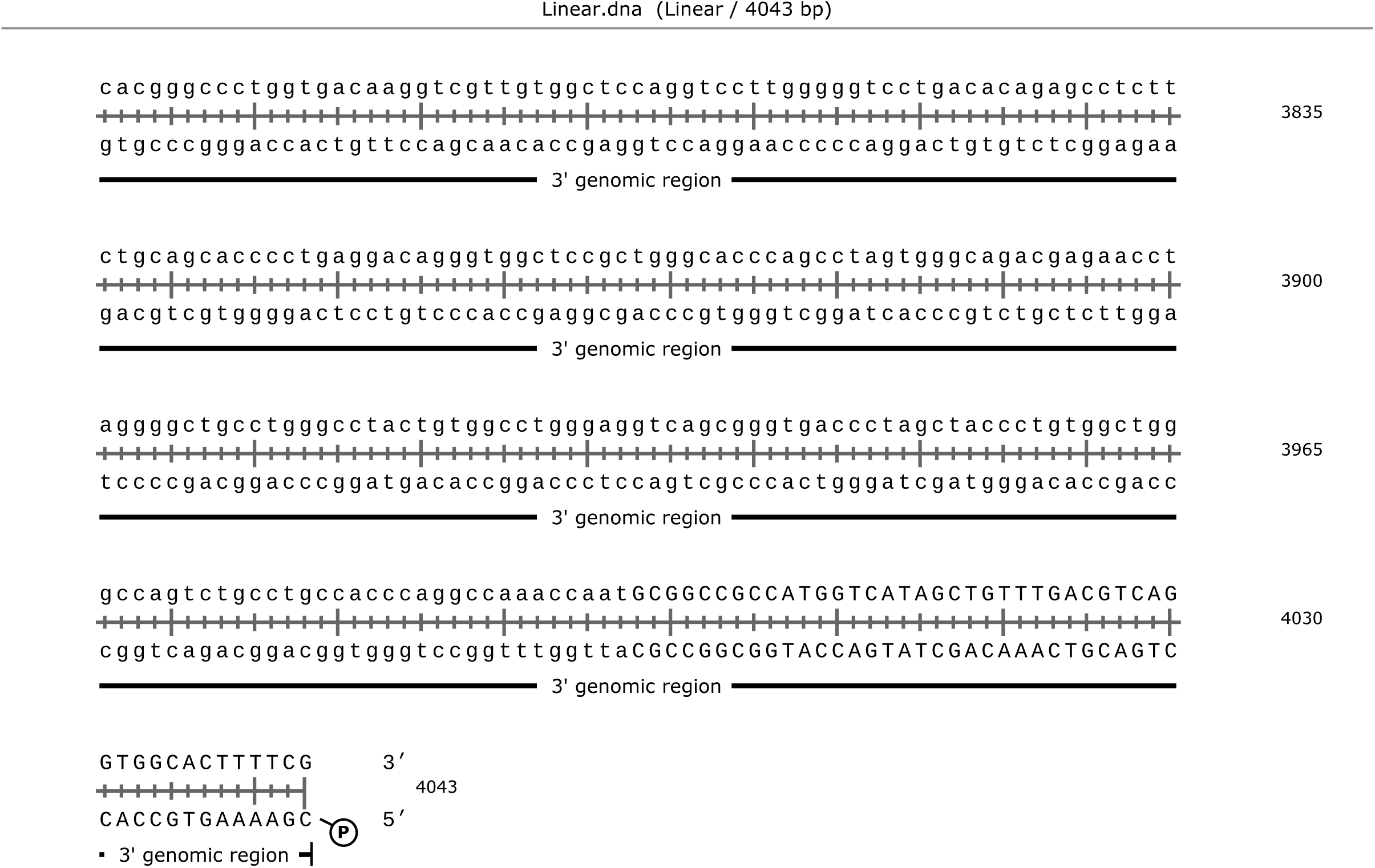

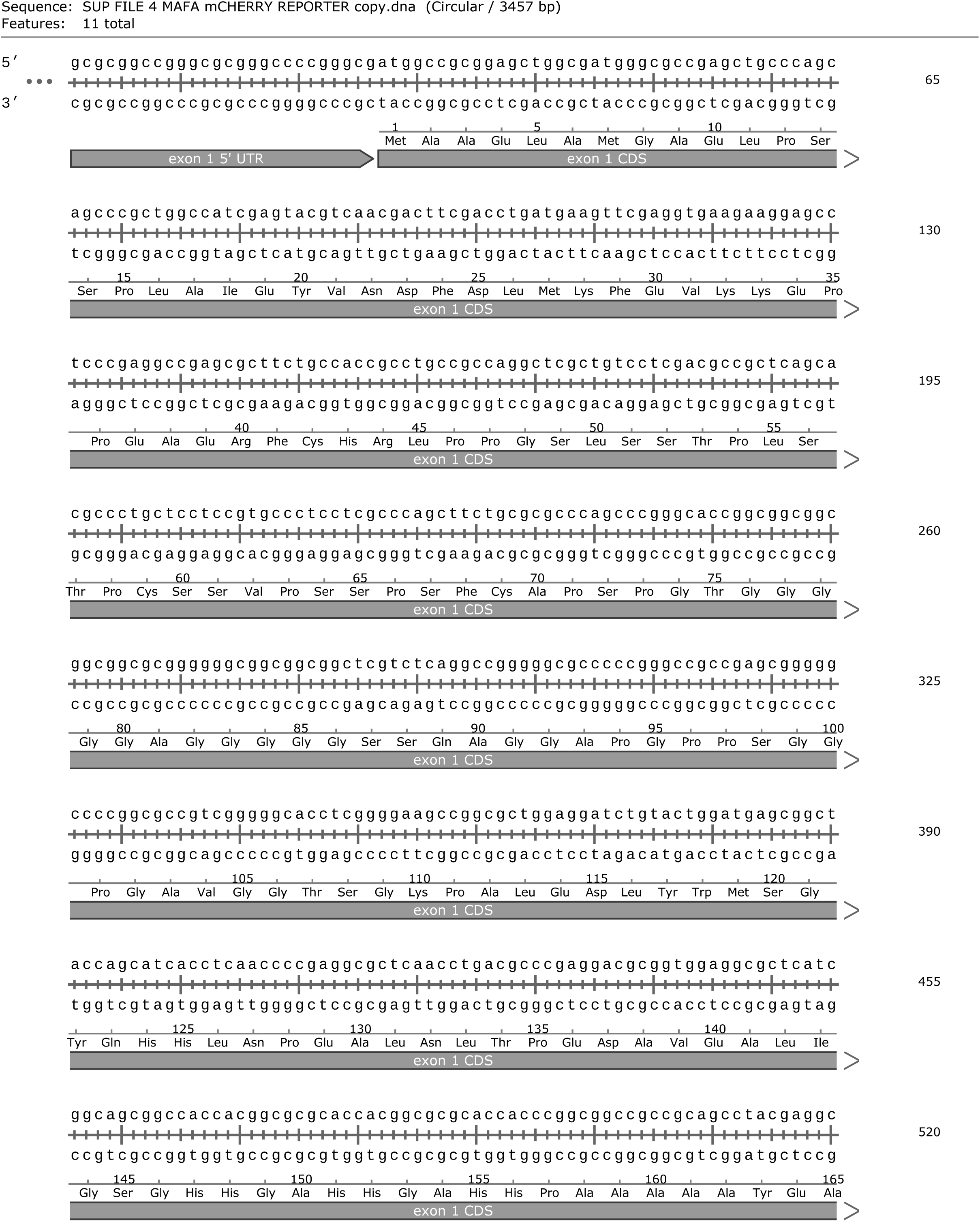
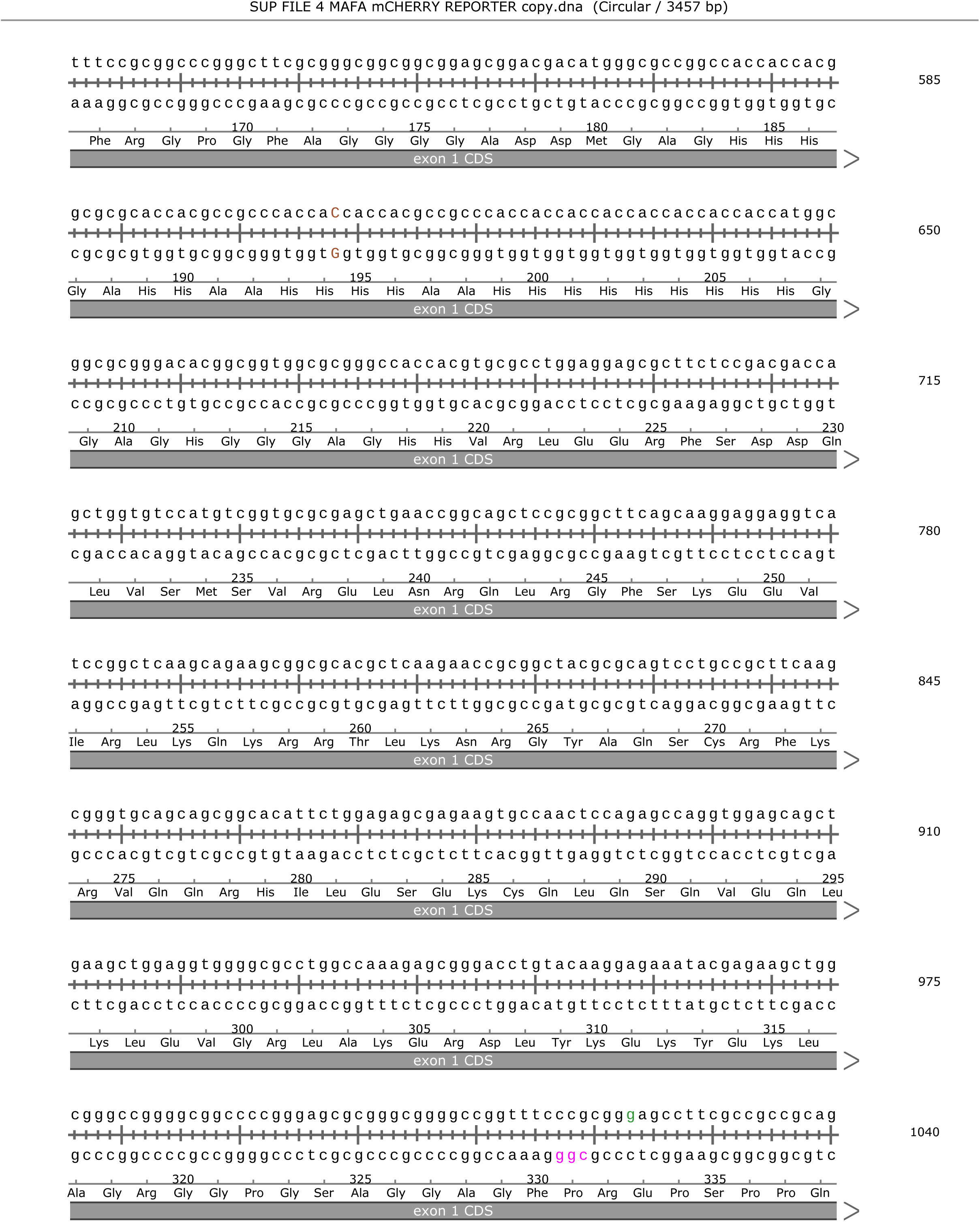
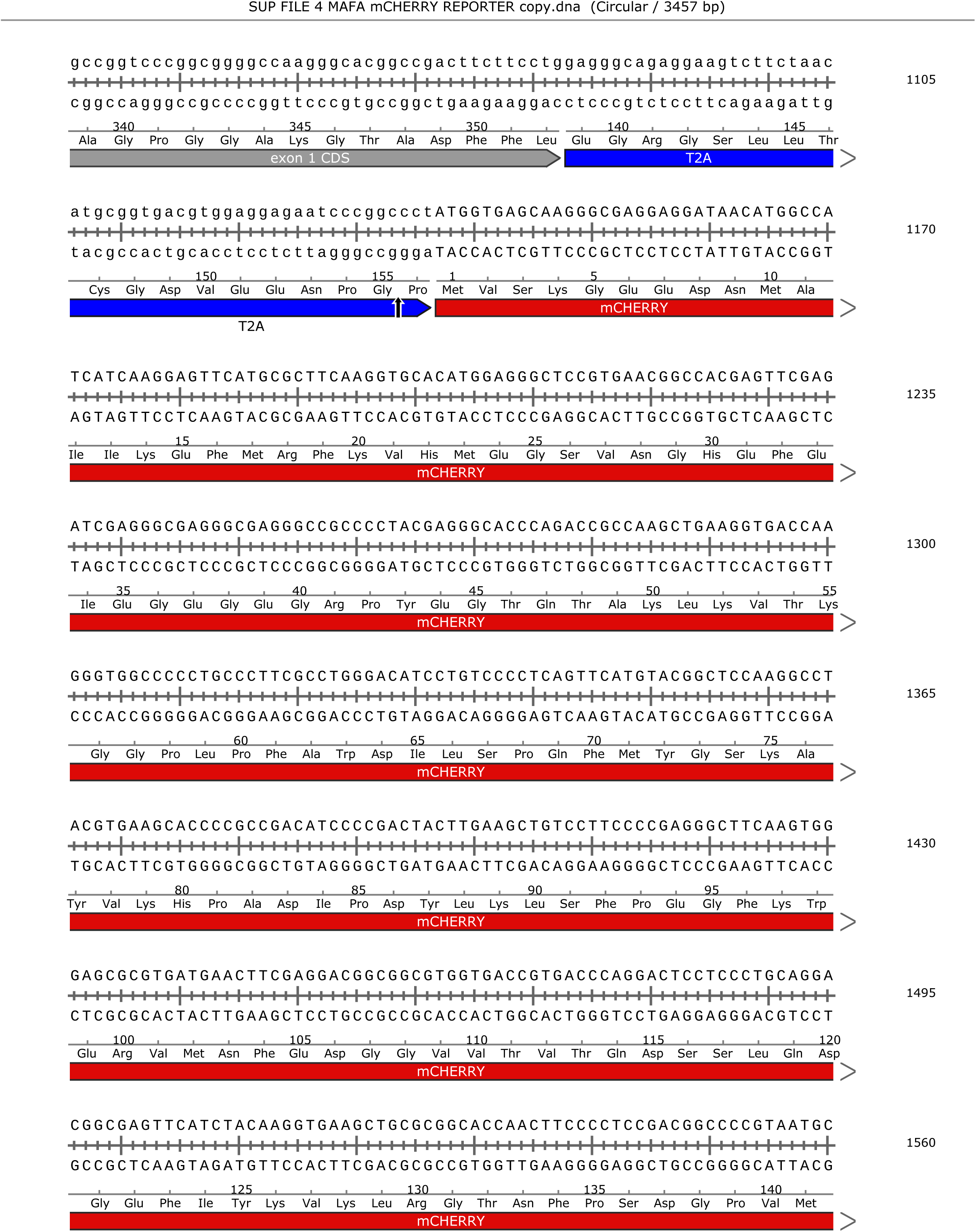
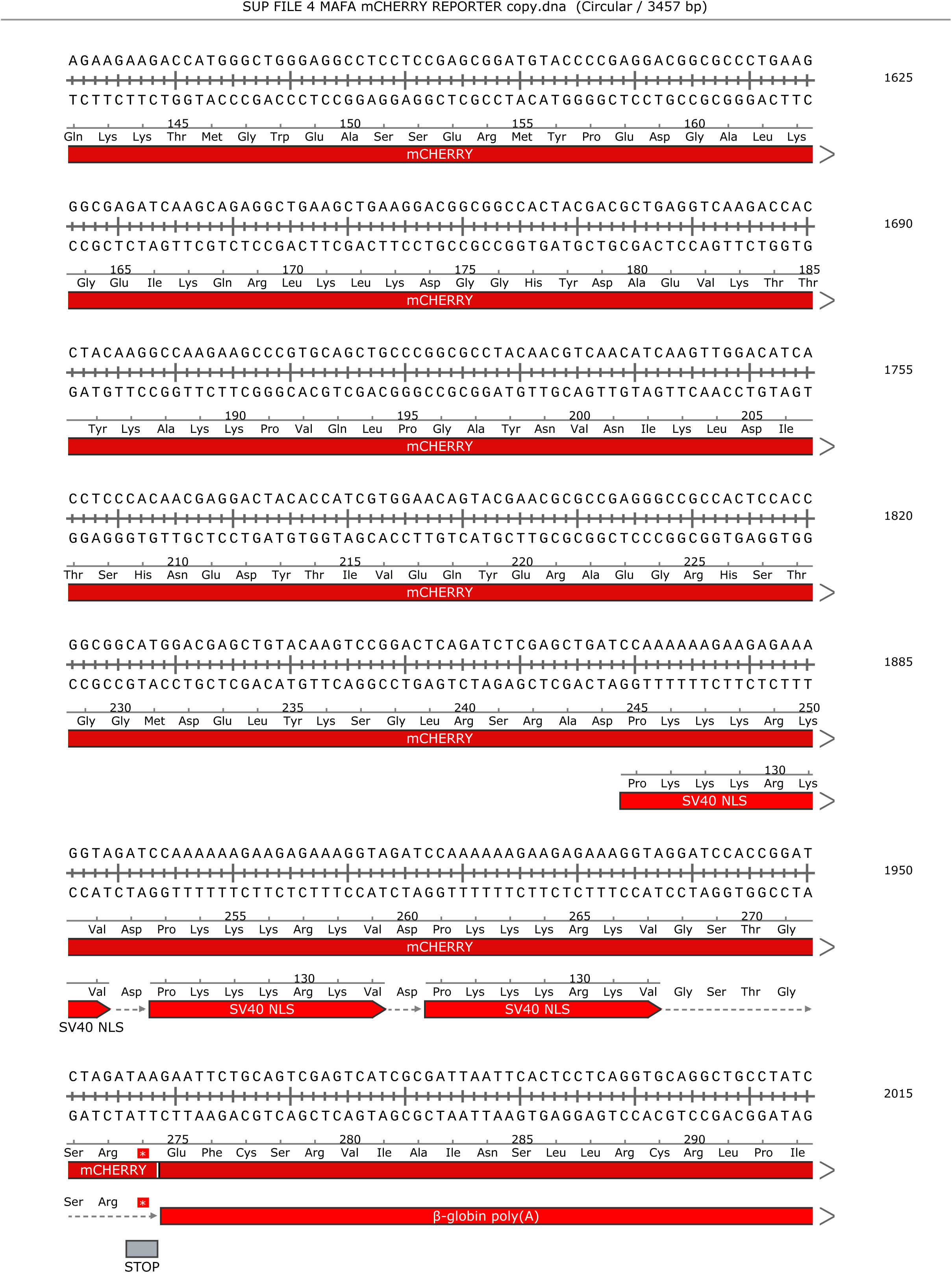
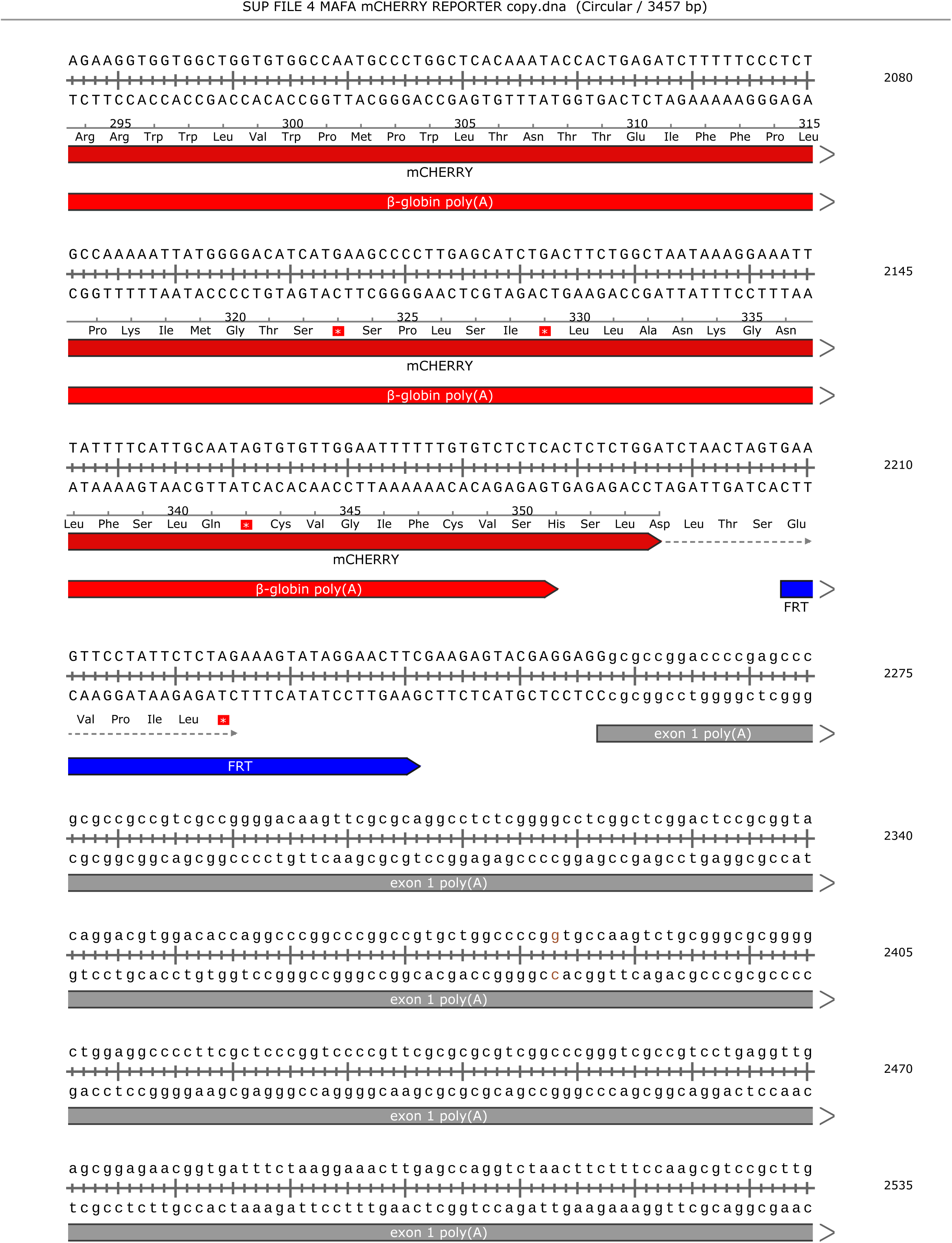
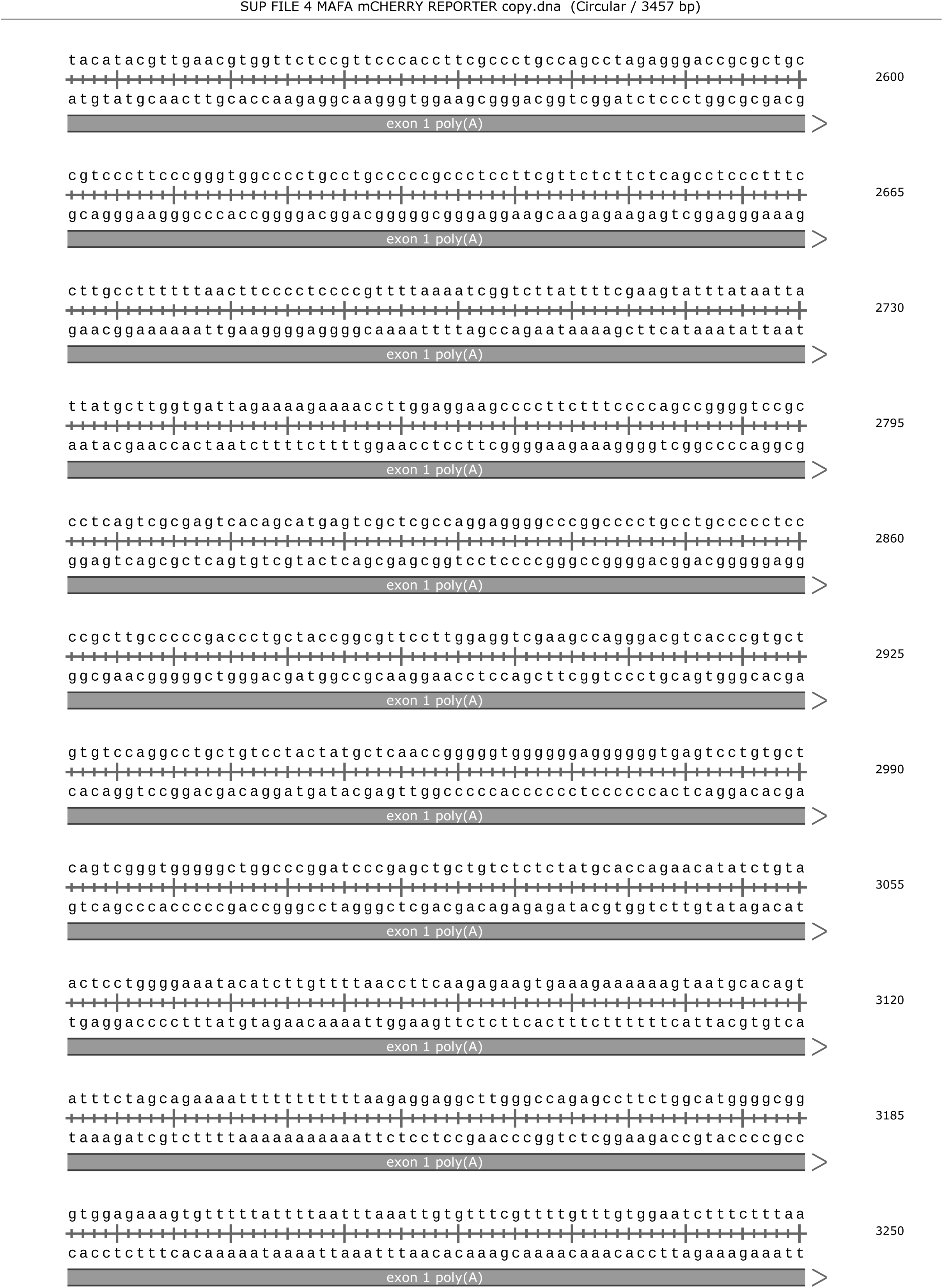
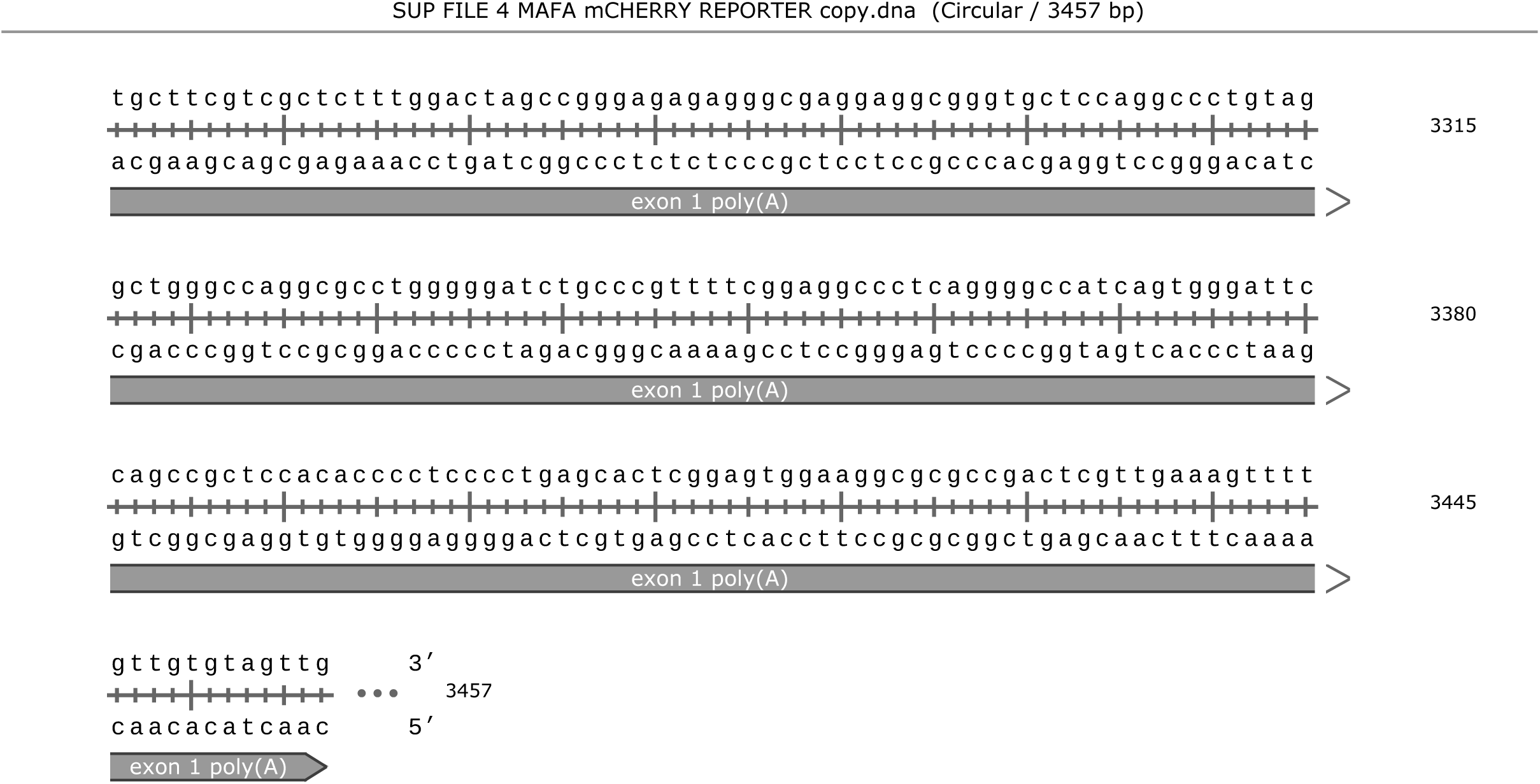
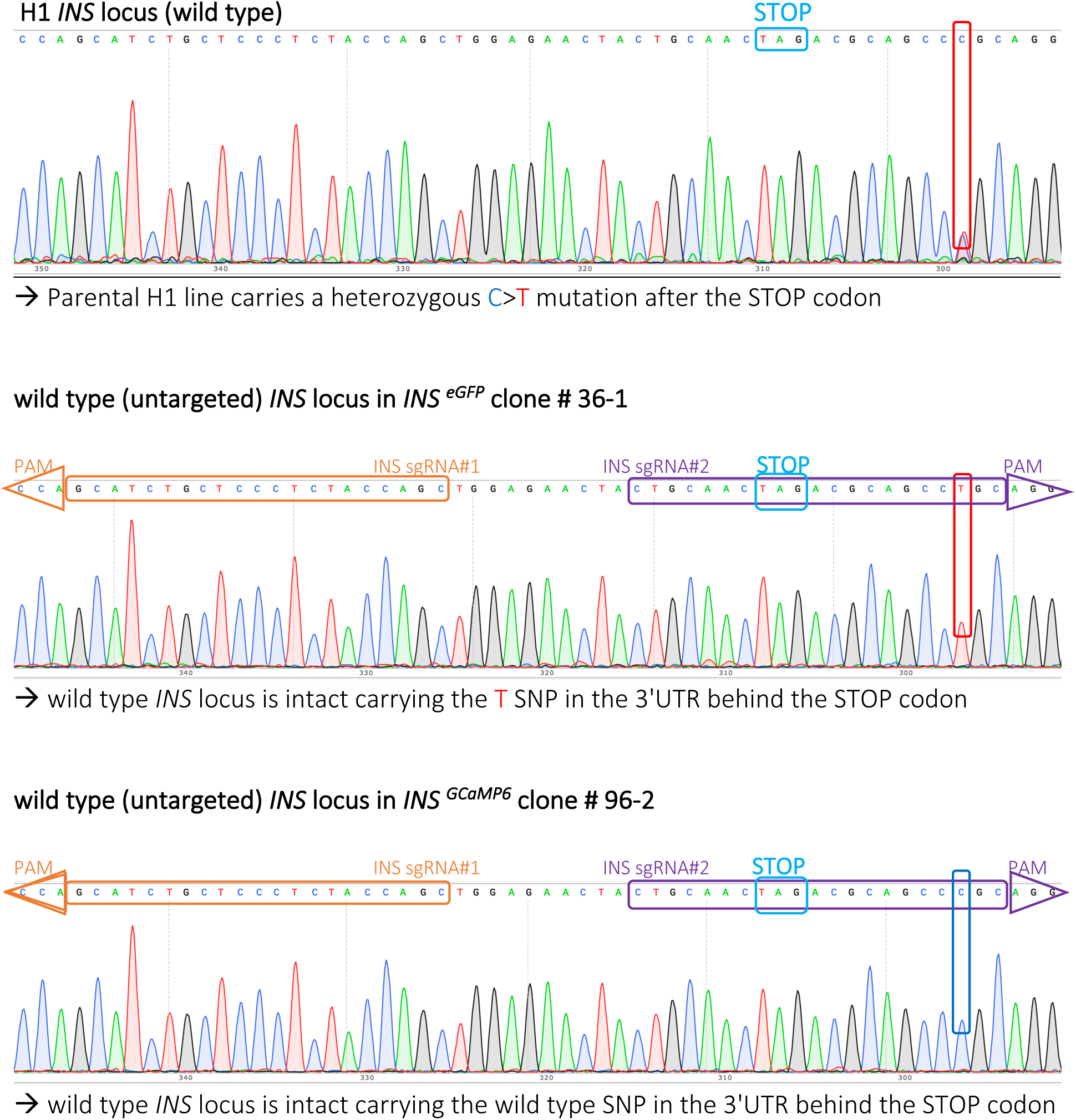
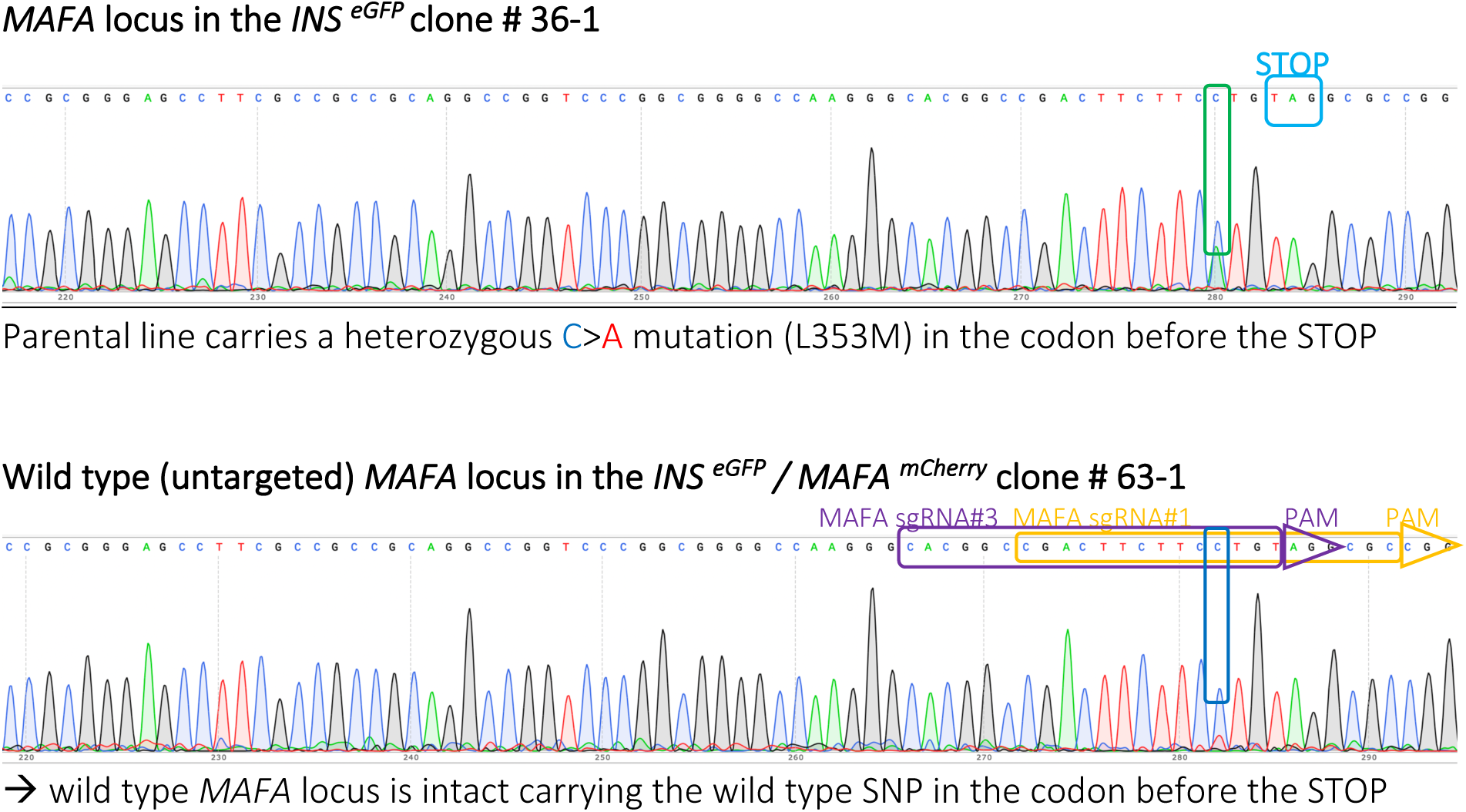
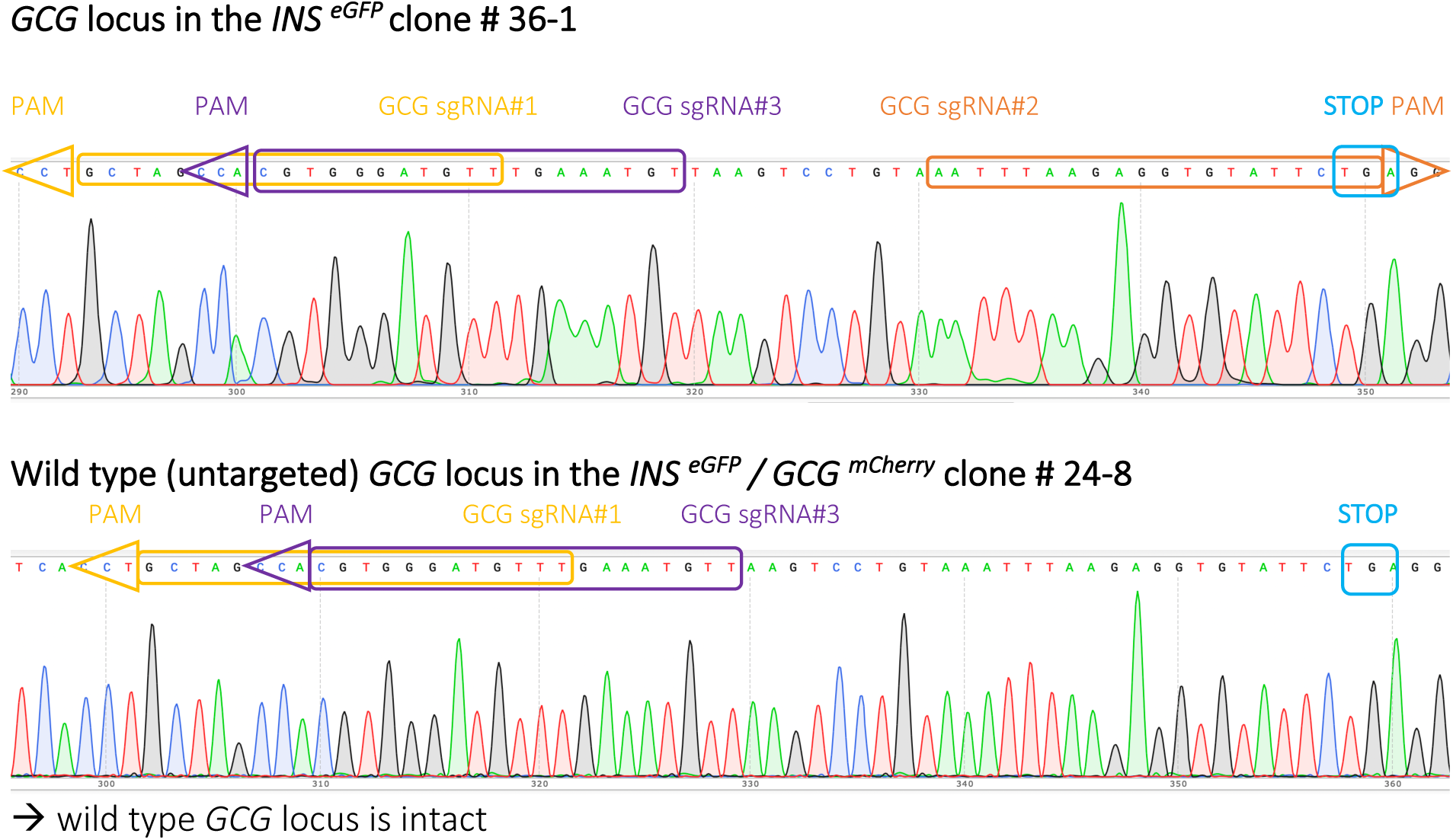

**Table S1.** Generation and / or origin of the DNA fragments used for the Gibson assembly of the targeting constructs.

| Allele | DNA fragments | Plasmid | Reference | Provider | Form |
| --- | --- | --- | --- | --- | --- |
| *INS*<sup>eGFP</sup> | pDTA-TK (plasmid backbone)<br>5' INS HA<br>T2A-eGFP<br>PGK-Neomycin<br>3' INS HA | pDTA-TK<br>hINS 5pHA<br>SpCas9-2A-GFP<br>PGKneoF2L2DTA<br>hINS 3pHA | Nam and Benezra 2009<br>this work<br>Nam and Benezra 2009<br>Hoch and Soriano 2006<br>this work | Addgene 22677.<br>custome made, IDT<br>Addgene 48138.<br>Addgene 13445.<br>custome made, IDT | NotI linearised<br>restriction fragment<br>PCR product<br>PCR product<br>restriction fragment |
| *GCG*<sup>mCHERRY</sup> | pDTA-TK (plasmid backbone)<br>5' GCG HA<br>T2A-mCHERRY<br>PGK-Hygromycin<br>3' GCG HA | modified pDTA-TK*<br>genomic PCR fragment<br>pGEMT_EGFPbait-T2A-Cherry-3xNLS<br>hSpCas9n-2A-Puro<br>genomic PCR fragment | this work<br>this work<br>N/A<br>Nam and Benezra 2009<br>this work | this work<br>this work<br>Dunja Knapp (CRTD)<br>Addgene 62987.<br>this work | NotI linearised<br>PCR product<br>PCR product<br>PCR product<br>PCR product |
| *MAFA*<sup>mCHERRY</sup> | pDTA-TK (plasmid backbone)<br>5' MAFA HA<br>T2A-mCHERRY<br>PGK-Hygromycin<br>3' MAFA HA | modified pDTA-TK*<br>hMAFA 5pHA<br>pGEMT_EGFPbait-T2A-Cherry-3xNLS<br>E2-Crimson-IRES-Puro_PGK-Hygro<br>hMAFA 5pHA | this work<br>this work<br>N/A<br>N/A<br>this work | this work<br>custom made, ThermoFisher<br>Dunja Knapp (CRTD)<br>Shahryar Khattak (CRTD)<br>custom made, ThermoFisher | NotI linearised<br>restriction fragment<br>PCR product<br>PCR product<br>restriction fragment |
| *INS*<sup>GCaMP6</sup> | pDTA-TK (plasmid backbone)<br>5' INS HA<br>GCaMP6<br>PGK-Neomycin<br>3' INS HA | modified pDTA-TK**<br>hINS 5pHA<br>pGP-CMV-GCaMP6s<br>PGKneoF2L2DTA<br>hINS 3pHA | this work<br>this work<br>Chen et al. 2013<br>Hoch and Soriano 2006<br>this work | this work<br>this work<br>Addgene 40753<br>Addgene 13445.<br>this work | NotI linearised<br>PCR product<br>PCR product<br>PCR product<br>PCR product |

\* The bGHpolyA from PGKneoF2L2DTA has substituted the SV40polyA of pDTA-TK as a SbfI / KpnI restriction fragment
\*\* The $\beta$ glob polyA from pGEMT\_EGFPbait-T2A-Cherry-3xNLS has substituted the SV40polyA of pDTA-TK using t

**Table S2.**
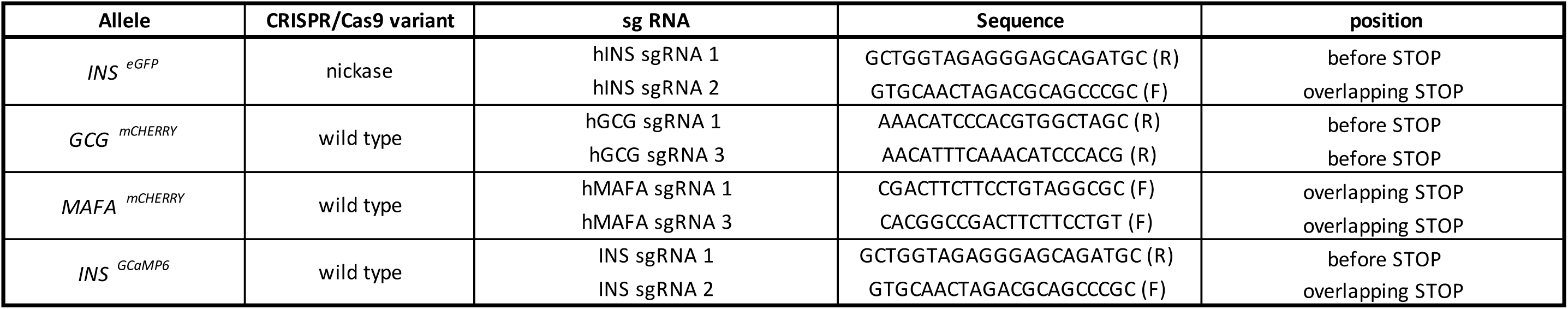
CRISPR Cas9 variant, sequence of the revers (R) or forward (F) sg RNAs used and their position relative to the STOP codon.

**Table S3.**
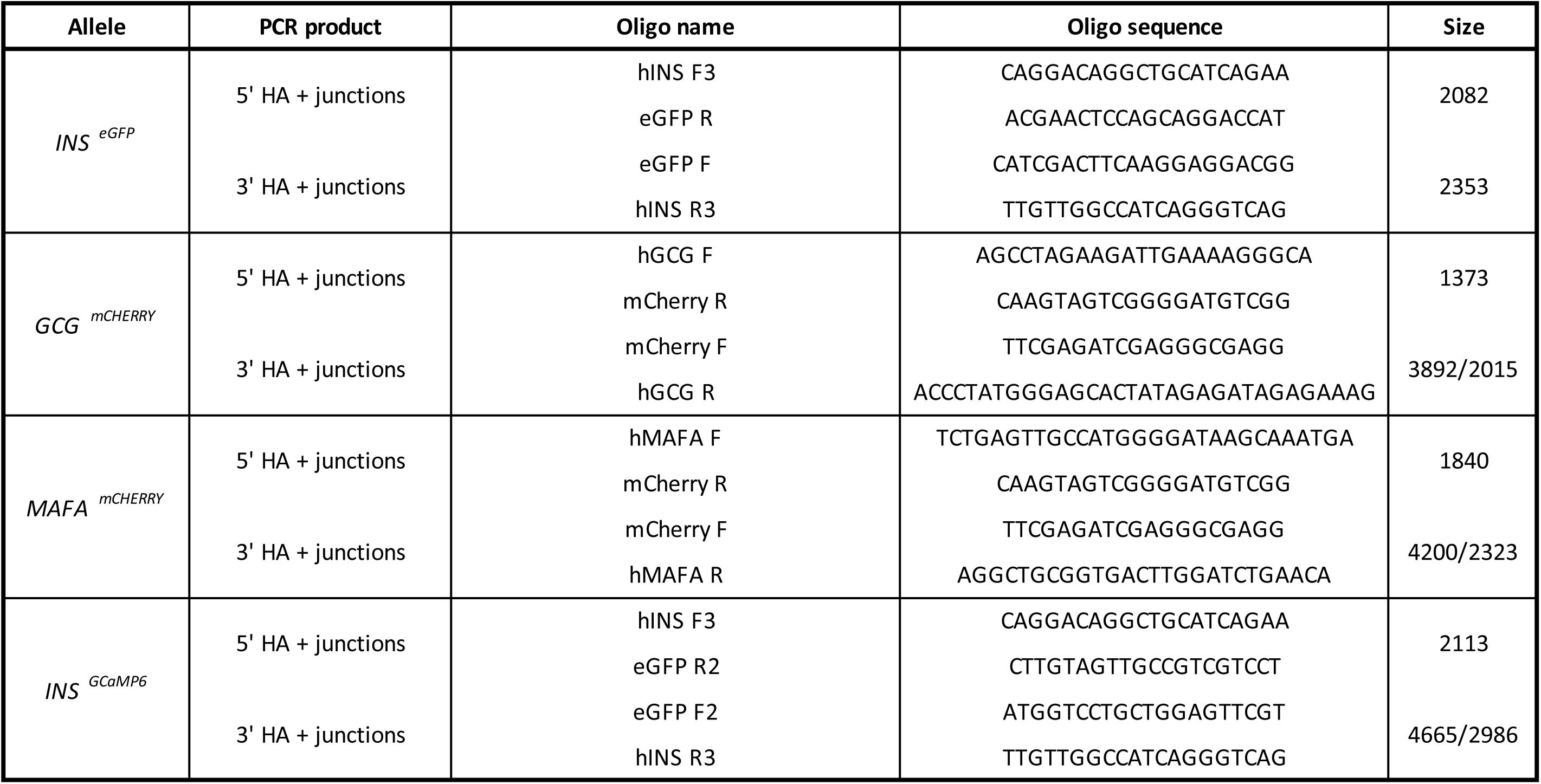
Genotyping and validation strategy of the targeted alleles. The amplified fragments were then Sanger sequenced.

**Table S4.** List of lines and corresponding expanded clones. Used clones are highlighted.

| LINE | CLONES |
| --- | --- |
| *INS*<sup>eGFP</sup> | # 4-1 |
|  | # 32-1 |
|  | # 36-1 |
| *INS*<sup>eGFP</sup> / *GCG*<sup>mCHERRY</sup> | # 24-8 |
|  | # 59-6 |
| *INS*<sup>eGFP</sup> / *MAFA*<sup>mCHERRY</sup> | # 59-1 |
|  | # 63-1 |
| *INS*<sup>GCaMP6</sup> | # 9-1 |
|  | # 96-2 |

**Table S5.**
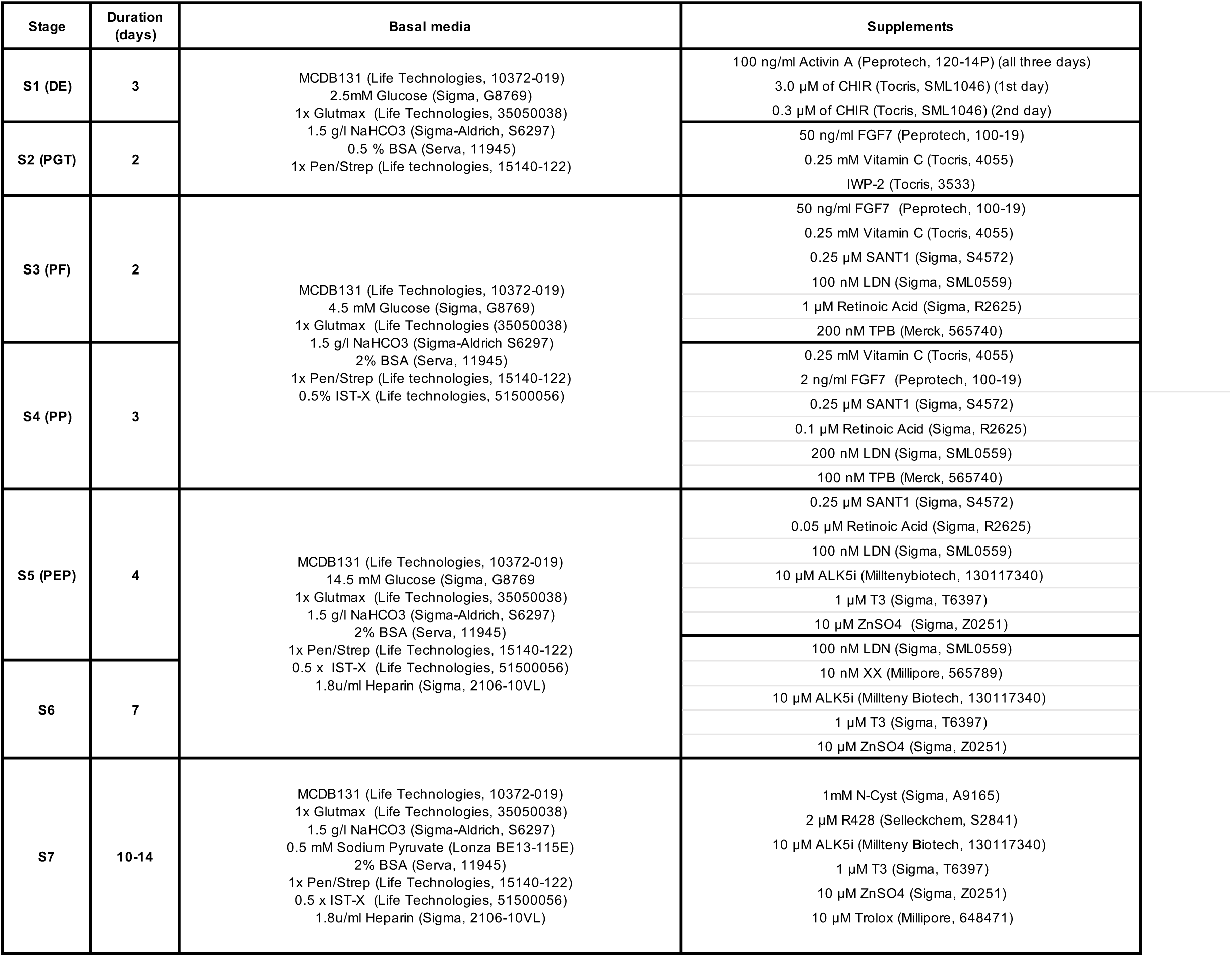
Basal media and supplements for S1-S7.

| Stage | Duration (days) | Basal media | Supplements |
| --- | --- | --- | --- |
| S1 (DE) | 3 | MCDB131 (Life Technologies, 10372-019)<br>2.5mM Glucose (Sigma, G8769)<br>1x Glutmax (Life Technologies, 35050038)<br>1.5 g/l NaHCO<sub>3</sub> (Sigma-Aldrich, S6297)<br>0.5 % BSA (Serva, 11945)<br>1x Pen/Strep (Life technologies, 15140-122) | 100 ng/ml Activin A (Peprotech, 120-14P) (all three days)<br>3.0 µM of CHIR (Tocris, SML1046) (1st day)<br>0.3 µM of CHIR (Tocris, SML1046) (2nd day) |
| S2 (PGT) | 2 |  | 50 ng/ml FGF7 (Peprotech, 100-19)<br>0.25 mM Vitamin C (Tocris, 4055)<br>IWP-2 (Tocris, 3533) |
| S3 (PF) | 2 | MCDB131 (Life Technologies, 10372-019)<br>4.5 mM Glucose (Sigma, G8769)<br>1x Glutmax (Life Technologies, 35050038)<br>1.5 g/l NaHCO<sub>3</sub> (Sigma-Aldrich, S6297)<br>2% BSA (Serva, 11945)<br>1x Pen/Strep (Life technologies, 15140-122)<br>0.5% IST-X (Life technologies, 51500056) | 50 ng/ml FGF7 (Peprotech, 100-19)<br>0.25 mM Vitamin C (Tocris, 4055)<br>0.25 µM SANT1 (Sigma, S4572)<br>100 nM LDN (Sigma, SML0559)<br>1 µM Retinoic Acid (Sigma, R2625)<br>200 nM TPB (Merck, 565740) |
| S4 (PP) | 3 |  | 0.25 mM Vitamin C (Tocris, 4055)<br>2 ng/ml FGF7 (Peprotech, 100-19)<br>0.25 µM SANT1 (Sigma, S4572)<br>0.1 µM Retinoic Acid (Sigma, R2625)<br>200 nM LDN (Sigma, SML0559)<br>100 nM TPB (Merck, 565740) |
| S5 (PEP) | 4 | MCDB131 (Life Technologies, 10372-019)<br>14.5 mM Glucose (Sigma, G8769)<br>1x Glutmax (Life Technologies, 35050038)<br>1.5 g/l NaHCO<sub>3</sub> (Sigma-Aldrich, S6297)<br>2% BSA (Serva, 11945)<br>1x Pen/Strep (Life Technologies, 15140-122)<br>0.5 x IST-X (Life Technologies, 51500056)<br>1.8u/ml Heparin (Sigma, 2106-10VL) | 0.25 µM SANT1 (Sigma, S4572)<br>0.05 µM Retinoic Acid (Sigma, R2625)<br>100 nM LDN (Sigma, SML0559)<br>10 µM ALK5i (Milltenybiotech, 130117340)<br>1 µM T3 (Sigma, T6397)<br>10 µM ZnSO<sub>4</sub> (Sigma, Z0251) |
| S6 | 7 |  | 100 nM LDN (Sigma, SML0559)<br>10 nM XX (Millipore, 565789)<br>10 µM ALK5i (Millteny Biotech, 130117340)<br>1 µM T3 (Sigma, T6397)<br>10 µM ZnSO<sub>4</sub> (Sigma, Z0251) |
| S7 | 10-14 | MCDB131 (Life Technologies, 10372-019)<br>1x Glutmax (Life Technologies, 35050038)<br>1.5 g/l NaHCO<sub>3</sub> (Sigma-Aldrich, S6297)<br>0.5 mM Sodium Pyruvate (Lonza BE13-115E)<br>2% BSA (Serva, 11945)<br>1x Pen/Strep (Life Technologies, 15140-122)<br>0.5 x IST-X (Life Technologies, 51500056)<br>1.8u/ml Heparin (Sigma, 2106-10VL) | 1mM N-Cyst (Sigma, A9165)<br>2 µM R428 (Selleckchem, S2841)<br>10 µM ALK5i (Millteny Biotech, 130117340)<br>1 µM T3 (Sigma, T6397)<br>10 µM ZnSO<sub>4</sub> (Sigma, Z0251)<br>10 µM Trolox (Millipore, 648471) |

**Table S6.** List of primary antibodies for immunofluorescense (IF) and flow cytometry (FC)

| Antigen | Species | Supplier, cat no | Application | Dilution |
| --- | --- | --- | --- | --- |
| OCT4 | Mouse | Santa Cruz, sc-5279 | IF/FC | 1:100/1:200 |
| SOX2 | Rabbit | Abcam, ab97959 | IF/FC | 1:200/1:100 |
| SSEA4 | Mouse | DSHB, MC-813-70 | IF | 1:80 |
| FOXA2 | Rabbit | Merck, 07-633 | IF/FC | 1:200 |
| SOX17 | Mouse | R&D, MAB1924 | IF/FC | 1:50 |
| NKX6.1 | Mouse | DSHB (F55A10) | IF/FC | 1:500/1:250 |
| PDX1 | Goat | R&D, AF2419 | IF/FC | 1:40 |
| SOX9 | Rabbit | Merck, AB5535 | IF/FC | 1:1000 |
| GCG | Mouse | Sigma, G2654 | IF | 1:500 |
| INS | Guinea pig | Abcam, ab7842 | IF | 1:500 |
| MAFA | Rabbit | Abcam, ab264418 | IF | 1:300 |
| GFP | Chicken | Abcam, ab13970 | IF | 1:2000 |
| mCherry | Rat | Chromotek, 5f8 | IF | 1:1000 |

**Table S7.** List of secondary antibodies.

| Antigen | Fluorophore | Species | Supplier, cat no | Dilution |
| --- | --- | --- | --- | --- |
| Goat IgG | Alexa 647 | Donkey | Invitrogen, A21447 | 1:500 |
| Guinea pig IgG | Alexa 647 | Goat | Abcam, ab150187 | 1:500 |
| Mouse IgG | Alexa 568 | Donkey | Invitrogen, A10037 | 1:500 |
| Mouse IgG | Alexa 568 | Goat | Invitrogen, A11004 | 1:500 |
| Chicken IgY | Alexa 488 | Goat | Abcam, ab150169 | 1:500 |
| Rabbit IgG | Alexa 488 | Donkey | Invitrogen, A21206 | 1:500 |
| Rabbit IgG | Alexa 488 | Goat | Invitrogen, A11070 | 1:500 |
| Rat IgG | Alexa 568 | Goat | Invitrogen, A11077 | 1:500 |

**Table S8.**
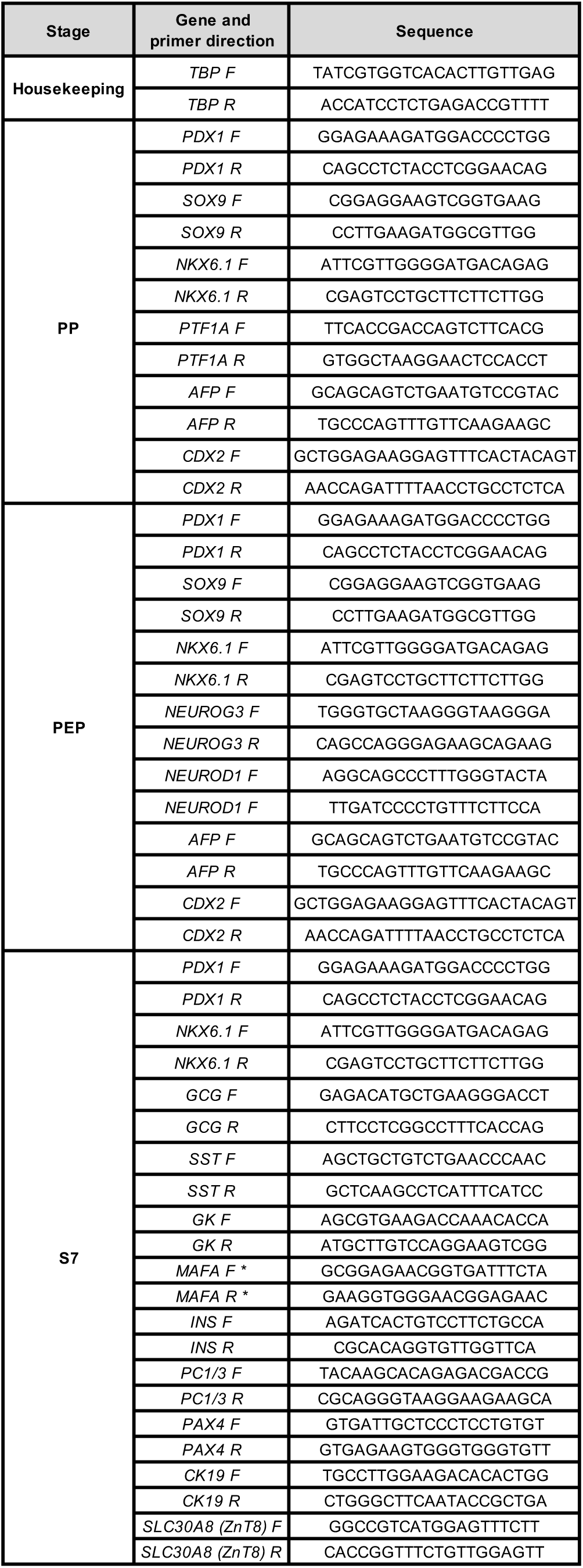
List of forward (F) and reverse (R) primers used for qPCR. * because of the very high GC content of the MAFA coding region the reverse primer had to be designed against the 3’ UTR. As a consequence, this pair of primers does not recognise the targeted allele

